# Discovery of broad-spectrum bacterial polyamine detoxification inhibitors as potential antivirulence agents and antibiotic adjuvants

**DOI:** 10.1101/2024.10.18.618978

**Authors:** Peri B. Moulding, Ronald S. Flannagan, Jonas Wong, Olmo Martín-Cámara, Ahmed M. Soliman, Sonsoles Martín-Santamaría, Wael Elhenawy, David E. Heinrichs, Omar M. El-Halfawy

## Abstract

The alarming rise in antimicrobial resistance reinforces an urgent need for new antimicrobial strategies. Host-or bacteria-derived chemicals, such as polyamines, present at infection sites often influence microbial virulence and antibiotic response. Polyamines are cationic small molecules overproduced by the host during infection, modulating immune responses. Polyamine detoxification by several pathogens correlates with increased virulence. We sought to uncover inhibitors of polyamine detoxification through a high-throughput whole-cell screen against the community-acquired methicillin-resistant *Staphylococcus aureus* (MRSA) USA300, identifying the polyamine analog OES2-0017 and the catechol derivative isoproterenol (OES1-1087), which synergized with polyamines in the low-micromolar range. Chemical genomics and enzymatic assays combined with computational studies revealed that both compounds prevented polyamine detoxification by spermine/spermidine acetyltransferase SSAT (SpeG) and another previously uncharacterized *S. aureus* SSAT (denoted PaiA_Sa_ herein). OES2-0017 directly inhibited the enzymes, whereas OES1-1087 reduced intracellular levels of acetyl-CoA required for SSAT activity. OES2-0017 perturbed the bacterial membrane at higher concentrations, likely causing the observed growth-inhibitory effects, whereas OES1-1087 increased membrane fluidity, likely increasing susceptibility to spermine-mediated membrane perturbation. The inhibitors showed broad-spectrum activity against various Gram-positive and Gram-negative bacteria, as well as *Candida albicans*. OES2-0017 abolished the spermine-mediated protection of MRSA USA300 from antibiotics, including vancomycin, phenocopying the Δ*speG* mutant and suggesting its potential utility as an antibiotic adjuvant. OES2-0017 eradicated SpeG-expressing *Salmonella* Typhimurium inside murine macrophages, whereas OES1-1087 reduced the *S.* Typhimurium burden in the liver and spleen in a murine gastrointestinal infection model, demonstrating their potential as antivirulence agents. Neither inhibitor exhibited cytotoxic activity against eukaryotic cells at their respective antimicrobial ranges. Small-scale structure-activity relationship experimental and computational analyses identified OES2-0017 analogs with improved specificity for the bacterial enzyme compared to the human SAT1 and no toxicity. This study provides novel antimicrobial compounds with broad-spectrum activity and a novel mode of action for multidrug-resistant priority pathogens.

## Introduction

Bacteria have developed resistance to all currently available antibiotics posing a large public health risk. The spread of antimicrobial resistance (AMR) has exacerbated the threat of bacterial infections, with 1.14 million global deaths directly attributed to bacterial AMR in 2021 alone ^1^. *Staphylococcus aureus* is a leading cause of hospital and community-acquired infections, resulting in considerable morbidity and mortality rates ^2^. *S. aureus* can cause multiple infections, including skin and soft tissue infections (SSTI), infective endocarditis, osteomyelitis, prosthetic-device infections, community-acquired pneumonia, and other systemic infections ^3^. Some strains of *S. aureus* have developed resistance to β-lactam antibiotics, coined methicillin-resistant *Staphylococcus aureus* (MRSA). MRSA strains have spread rapidly, and in 2021 alone, it was estimated that 196,000 deaths were directly attributable to MRSA infections worldwide ^1^. Notably, the MRSA strain USA300 has become the most common infection-causing *S. aureus* strain in Canada and the USA ^4,5^.

While we continue to lose clinically useful antibiotics due to the rise of AMR, the discovery and development of antimicrobials with novel chemical structures or cellular targets have drastically slowed ^6,7^. One of the potential reasons for the antibiotic discovery void is that antimicrobial discovery efforts traditionally relied on antimicrobial susceptibility testing (AST) conditions that use culture media such as Mueller-Hinton broth (MHB) ^8^. These test conditions do not accurately reflect those at the infection site, including the chemicals bacteria may encounter within the host. There is an expansive array of molecules produced by the host, resident microbiota, and the infecting pathogen at the infection site, which may influence microbial virulence and antibiotic responses ^9–11^. Such chemical-mediated effects might alter the treatment outcomes and may offer new targets for antimicrobial intervention. One prominent class of compounds is polyamines, which are biogenic small cationic amines, such as the diamine putrescine, triamine spermidine, and tetramine spermine, required for normal cell functioning in most living organisms ^12^. Polyamines have been observed at higher concentrations in host cells during infection compared to uninfected healthy controls ^13–16^, and can be found at millimolar concentrations in eukaryotes ^17,18^, where they have immunomodulatory roles ^19,20^.

Most bacteria produce polyamines endogenously, with *S. aureus* being one of few exceptions that neither produce spermine or spermidine nor require them for homeostasis ^21–23^. For polyamine producers, polyamine levels are tightly regulated to maintain the level required for homeostasis, balancing uptake and biosynthesis with detoxification and efflux to prevent toxicity resulting from high and low polyamine cellular content ^12,18^. Most *S. aureus* strains, being polyamine non-producers, are typically more susceptible to these compounds; however, USA300 is resistant to exogenous spermine and spermidine ^21^. The virulence and prevalence of USA300 have been partially attributed to the acquisition of the arginine catabolic mobile element (ACME) ^13,24,25^. The ACME was horizontally transferred to *S. aureus* from *Staphylococcus epidermidis*, and encodes SpeG, a member of the GCN5-related *N*-acetyltransferase (GNAT) family ^25^. SpeG is a spermine/spermidine *N*-acetyltransferase (SSAT) that uses intracellular acetyl-Coenzyme A (AcCoA) to acetylate spermine and spermidine rendering them less toxic to the cell, resulting in the observed resistance to exogenous polyamines ^13,21,23,25^. The ACME also encodes the arginine-deaminase (*arc*) system which converts arginine to ornithine while producing ATP and ammonia, allowing USA300 to withstand acidic environments, such as those found on the skin ^13^. The host then converts ornithine produced via the ACME-Arc system to polyamines as shown in a murine SSTI model ^13^. Therefore, by carrying *speG,* USA300 can resist the excessive host polyamine production of the skin, increasing its virulence ^13,26^. While a Δ*speG* mutant showed reduced virulence relative to wild-type USA300 in murine skin infection ^13^, a mutant lacking the entire ACME locus did not exhibit reduced virulence in murine pneumonia or skin infection ^27^. The latter observation likely stems from the lack of the ACME-Arc system that is necessary to drive excessive host polyamine production ^13^. Further, the levels of host polyamines vary across infection sites, which may explain the diminished role of SpeG in virulence in murine sepsis where host polyamine levels are not sufficient to kill *S. aureus* ^13^. Additionally, exogenous polyamines showed contrasting effects on biofilm formation. They were observed to increase biofilm formation, which was partially attributable to the increase in pH upon addition of the polyamine to growth media ^25^. In contrast, exogenous polyamines exerted biofilm inhibitory activity in a dose-dependent manner at neutral pH ^23^. A Δ*speG* mutant was more susceptible to the polyamine biofilm inhibitory effects than the wild-type ^25^.

Polyamine detoxification via SSAT homologs contributed to virulence in other bacterial species. For example, SpeG is required for the survival of *Salmonella enterica* serovar Typhimurium in macrophages ^28^ and the overexpression of PmvE (an SSAT) in *Enterococcus faecalis* increases virulence in a *Galleria mellonella* infection model ^29^. Polyamines are also known to alter bacterial susceptibility to antibiotics ^30–34^ and to influence virulence phenotypes such as biofilm formation ^35–37^, intracellular growth ^38^, colonization ^39^, and resistance to reactive oxygen species ^40^. The observed involvement of polyamine detoxification by pathogens in their virulence and the potential for polyamine-mediated alterations to antibiotic susceptibility suggests that inhibiting bacterial polyamine detoxification is an attractive strategy to reduce virulence and prevent polyamine-mediated antibiotic resistance. Indeed, new treatment strategies to combat the spread of AMR are desperately needed, and antivirulence agents and antibiotic adjuvants have been suggested as potential therapeutic options. Antivirulence agents target processes dispensable for bacterial viability but that the pathogen uses to cause infection; therefore, by targeting these processes the host immune system can more easily clear the infection ^41^. Antibiotic adjuvants are compounds that potentiate the activity of existing antibiotics, restoring their effect against resistant bacteria or improving their efficacy against intrinsically resistant organisms ^42^. Further, drug repurposing, a strategy in which existing therapeutics approved for certain indications are redeployed to treat another condition, may accelerate the drug development process as human safety and pharmacokinetic profiles are already established for these drugs ^43^.

Here, we set out to identify inhibitors of bacterial polyamine detoxification mechanisms through a cell-based high throughput screen. We discovered the polyamine analog OES2-0017 and the catechol derivative, isoproterenol, OES1-1087, which showed potent synergy with polyamines against various Gram-positive and Gram-negative pathogens. Both molecules disrupted the SSAT-mediated acetylation of polyamines, where OES2-0017 directly inhibited the enzymes whereas OES1-1087 acted mostly indirectly by reducing the intracellular concentration of AcCoA. Further, OES2-0017 perturbed the bacterial membrane while OES1-1087 increased membrane fluidity of *S. aureus,* which may have contributed to sensitization to membrane perturbation by spermine. We observed standalone antimicrobial activity *in cellulo* in macrophages (by OES2-0017) and *in vivo* in murine gastrointestinal infection model (by OES1-1087) and that OES2-0017 abolished the polyamine-mediated resistance of MRSA USA300 to vancomycin and other antibiotics. Neither compound is cytotoxic to HepG2 cells or lytic in a hemolysis assay at the same concentration range that results in polyamine synergy. Together, this study describes novel antimicrobial compounds with broad-spectrum activity and potential antivirulence and antibiotic adjuvant applications.

## Results

### A high-throughput chemical screen identifies potential polyamine detoxification inhibitors

To identify inhibitors of polyamine detoxification mechanisms, we undertook two screens against *S. aureus* USA300. We chose to screen against *S. aureus* USA300 because it lacks *de novo* polyamine biosynthesis and thus ensures the observed phenotypes are due to the response to exogenous polyamines. In an unbiased approach, we screened a library of 2560 small molecules (Spectrum collection, MicroSource Discovery Systems, Inc.), comprised of previously approved drugs and natural product derivatives, at 20 μM in the presence of 2.5 mM of spermine [equivalent to 1/4^th^ the minimum inhibitory concentration (MIC) of spermine in 384-well plates] against USA300. In parallel, we took a rationale-based approach by assembling a collection of 83 polyamine analogs with at least 60% structural similarity to substrates and products of polyamine biosynthesis and detoxification enzymes in bacteria, reasoning that such analogs have a higher probability of interacting with, and potentially inhibiting, the polyamine-related enzymes of interest (Supplementary tables 1 and 2). Notably, several polyamine biosynthesis inhibitors are analogous to natural polyamines or substrates of their biosynthetic enzymes ^44–47^, which supports this rationale-based approach. We screened the polyamine analog library at 20 μM (similar to the Spectrum collection screen) in the presence of spermine at 1/4^th^ the MIC (5 mM in 96-well plates). The screen was performed with the hypothesis that a compound inhibiting polyamine detoxification would exhibit synergy with spermine due to build up of toxic levels of the polyamine at relatively lower doses in cells unable to detoxify it. Combined, the screens resulted in 187 primary hits showing at least 80% growth inhibition. We then excluded known antimicrobials in the Spectrum collection yielding 129 hits (Figure 1 A and B). We tested these hits in follow-up dose-response assays in the presence and absence of a single spermine concentration (Supplementary Fig. 1); only 8 compounds showed a reduction in MIC in the presence of spermine (Figure 1A). We then tested these eight compounds in checkerboard assays with spermine; six compounds, OES1-1087, OES1-0507, OES1-1639, OES1-1238, OES2-0052, and OES2-0017, showed synergy with spermine (Figure 1 C-H). Synergy was determined by calculating the fractional inhibitory concentration index (FICI), with an FICI value ≤ 0.5 defined as a synergistic interaction. Simultaneously, synergy scores were calculated using SynergyFinder using the ZIP reference model and a score of < -10 was classified as antagonism, between -10 and 10 additive, and > 10 as synergy ^48,49^. OES1-1087 exhibited the strongest synergistic interaction with spermine with an FICI value of 0.14 ± 0.03 followed by OES1-0507 (0.28 ± 0.08), OES2-0017 (0.29 ± 0.07), OES1-1639 (0.42 ± 0.07), OES1-1238 (0.46 ± 0.14), and finally OES2-0052 (0.50 ± 0), with synergy scores largely following this trend (Figure 1 C-H). OES1-0507, OES1-1087, and OES1-1639 are benzenediols, OES1-1238 is an acridine, and OES2-0052 and OES2-0017 are amines. In addition to synergizing with polyamines, the six hits exhibited growth inhibitory effects against USA300 with OES2-0017 being the most potent at an MIC of 20 μM (Figure 1 C-H). Of note, we prioritized OES2-0017 and OES1-1087 (isoproterenol) for further characterization based on their relative potencies (spermine synergy at FICI < 0.35) and as representatives of different chemical classes (polyamine and catechols, respectively). We confirmed the structures of the two prioritized compounds by NMR (Supplementary Fig. 2).

**Figure 1.**
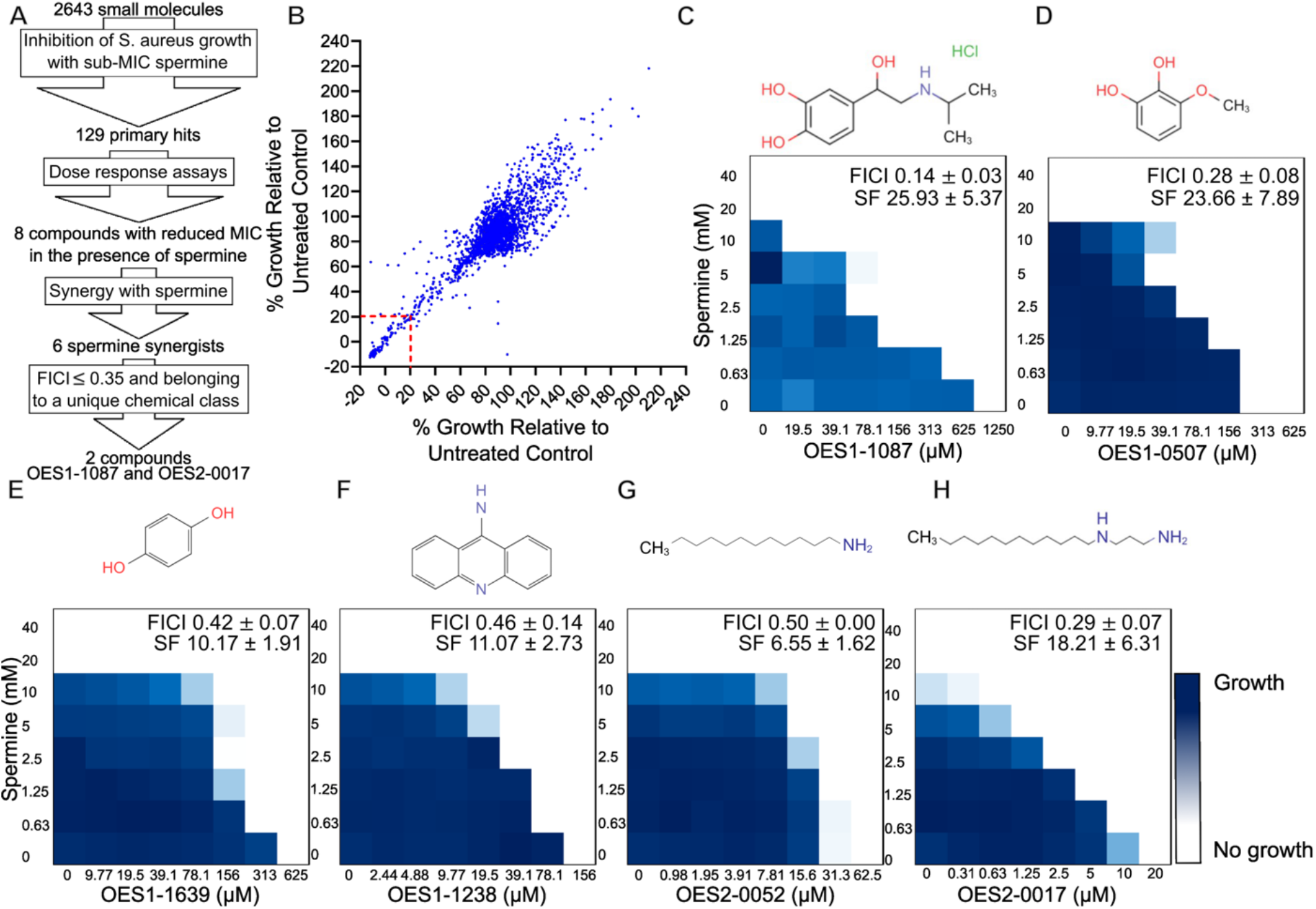
High throughput screening identifies potent spermine synergists. **A)** Workflow undertaken to identify chemical inhibitors of polyamine resistance. **B)** Replica plot of chemical screens performed to identify inhibitors of polyamine detoxification in *S. aureus* USA300. **C-H)** Chemical structures and representative checkerboard assays of the hits OES1-1087 (n=14) **(C)**, OES1-0507 (n= 4) **(D)**, OES1-1238 (n=3) **(E)**, OES1-1639 (n=3) **(F)**, OES2-0052 (n=3) **(G)**, and OES2-0017 (n=16) **(H)** with spermine against *S. aureus* USA300. FICI values and Synergy Finder (SF) scores are reported as mean ± standard deviation of the indicated number of replicates.

### A chemogenomic screen to uncover the putative mechanism of spermine synergy

We undertook a genome-wide approach to identify the putative mechanism of spermine synergy by the screen hits, reasoning that a mutant with disruption in their putative target would exhibit both increased spermine susceptibility and loss or reduction in spermine-hit compound synergy. First, we sought to uncover the determinants of polyamine detoxification whose mutants would exhibit increased susceptibility to spermine by screening the Nebraska Transposon Mutant Library (NTML), a sequence-defined transposon mutant library covering the non-essential genome of *S. aureus* USA300 ^50^, against spermine at 1/8^th^ and 1/16^th^ the wild-type MIC (2.5 and 1.25 mM, respectively). *speG*::Tn was sensitized to spermine, in agreement with previous reports ^21^, thus serving as an internal control for the screen. Mutants in another four determinants not previously linked to the response to polyamines, *tcaA*::Tn, *cls*::Tn, *prmC*::Tn, and *pyrP*::Tn, were also sensitized to spermine (Figure 2 A). Dose-response assays showed an 8-fold reduction in spermine MIC in *speG*::Tn, and a decrease in growth of the other four mutants at ½ MIC relative to the wild-type strain (Supplementary Fig. 3). We constructed an in-frame unmarked deletion of *speG* given that its transposon mutant had the most pronounced spermine MIC shift; Δ*speG* confirmed the observed spermine susceptibility shift (Supplementary Fig. 3). *tcaA* encodes a predicted transmembrane protein and is implicated in teicoplainin resistance ^51^. *cls* encodes a cardiolipin synthase involved in modulating the phospholipid composition of the membrane^52^. *prmC* encodes a release factor methyltransferase, and *pyrP* an uracil-xanthine permease. The molecular basis linking these determinants to polyamine response warrants further investigation, which falls beyond the scope of this study. Notably, *sbnB*::Tn, disrupted in a gene encoding an enzyme involved in the siderophore staphyloferrin B biosynthesis ^53^, was initially identified in the screen and showed an 8-fold spermine MIC reduction; however, a Δ*sbnB* mutant was not sensitized to spermine (Supplementary Fig. 3). Whole-genome sequencing and subsequent PCR analysis of the *sbnB*::Tn mutant revealed an additional truncation of ∼46.8 kb comprising the SCC*mec* cassette and the ACME locus, including *speG* and *mecA* (Supplementary Fig. 3); hence, *sbnB* was excluded from the putative determinants list.

**Figure 2.**
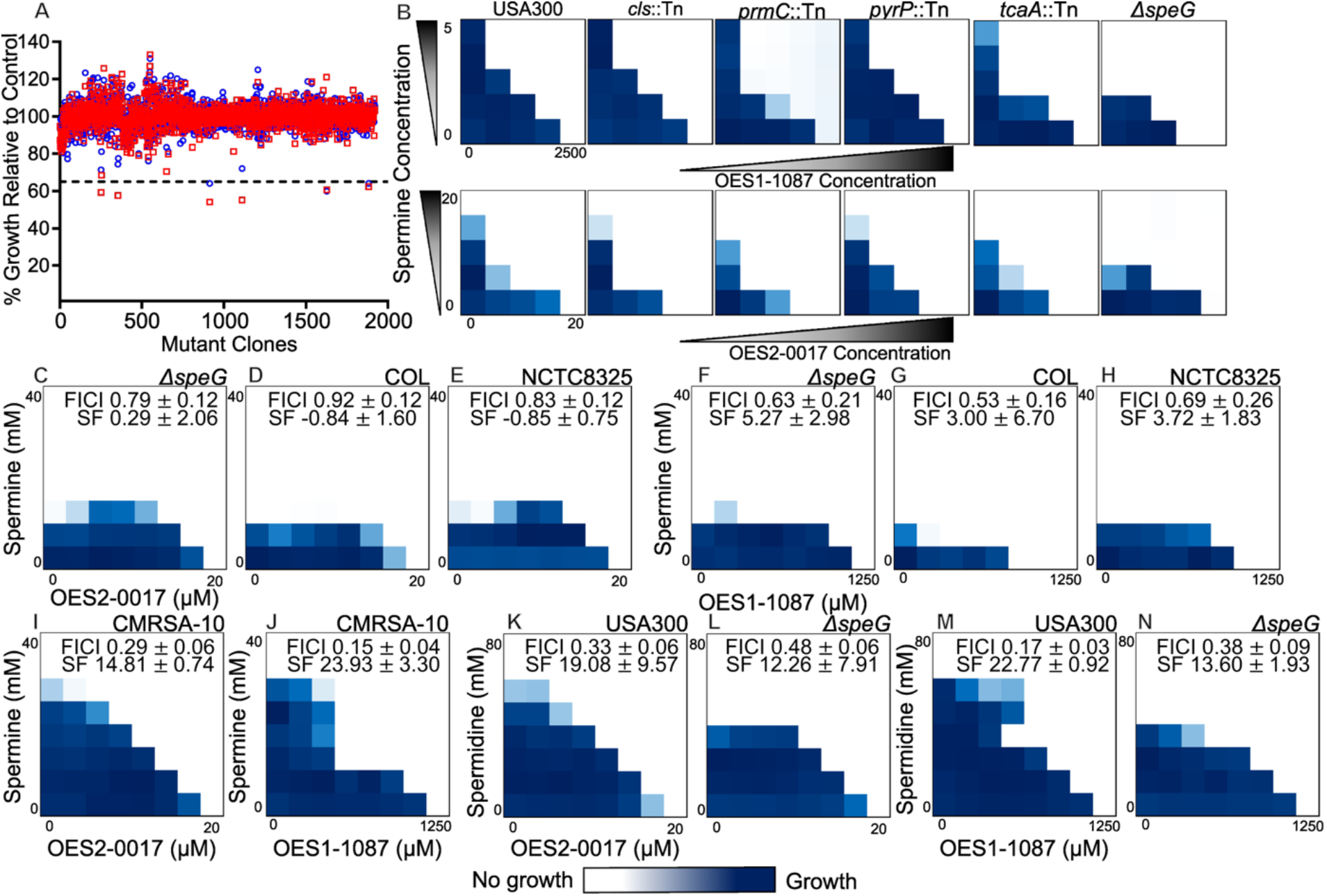
Chemogenomic screen identifies *speG* as the putative target of OES2-0017 and OES1-1087. **A)** Chemogenomic screen of the Nebraska Transposon Mutant Library with spermine at 1/8^th^ (red squares) and 1/16^th^ (blue circles) the MIC. **B)** Mini-checkerboard assays performed by testing 2-fold dilutions of each of spermine and OES1-1087 or OES2-0017 against polyamine resistance determinants uncovered in **(A)** including *S. aureus* USA300, *cls*::Tn, *prmC*::Tn, *pyrP*::Tn, *tcaA*::Tn, and Δ*speG*. **C-N)** Representative checkerboard assays performed by testing 2-fold dilutions of each test compound. **C-E)** spermine and OES2-0017 against *S. aureus* Δ*speG* (n=11) **(C)**, COL (n=3) **(D)**, NCTC8325 (n=3) **(E)**. **F-H)** Spermine and OES1-1087 against *S. aureus* Δ*speG* (n=9) **(F)**, COL (n=4) **(G)**, NCTC8325 (n=4) **(H)**. **I)** OES2-0017 and spermine against CMRSA-10 (n=3), **J)** OES1-1087 and spermine against CMRSA-10 (n=3). OES2-0017 and spermidine against **K)** USA300 (n=7) and **L)** Δ*speG* (n=6). OES1-1087 and spermidine against **M)** USA300 (n=3) and **N)** Δ*speG* (n=3). Checkerboards against Δ*speG* were performed on the same day as their counterparts against the wild-type strain shown in Figure 1. FICI values and Synergy Finder (SF) scores are reported as mean ± standard deviation of the indicated number of replicates.

Next, we resupplied the hit compounds and conducted mini-checkerboard assays with spermine against the five identified polyamine resistance determinants to detect loss or reduction in synergy. The spermine synergistic interaction with OES2-0017 and OES1-1087 was lost or reduced in the Δ*speG* mutant (Figure 2 B), suggesting that SpeG is the putative target of these compounds. Notably, spermine synergy was also lost in the other four de-prioritized hit compounds against Δ*speG* mutant (Supplementary Fig. 4), suggesting that all six hits uncovered from the screens share a similar molecular target for spermine synergy.

We performed checkerboard assays with spermine and OES2-0017 or OES1-1087 against other *speG^+^* and *speG^-^S. aureus* strains. The tested strains included another USA300 strain CMRSA-10 that encodes *speG* as well as two strains that do not encode *speG* including the MRSA strain COL and the methicillin-sensitive *Staphylococcus aureus* (MSSA) strain NCTC8325. The *speG^-^*strains, whether MRSA or MSSA, displayed a loss or reduction in synergy between spermine and OES2-0017 or OES1-1087, phenocopying the USA300 Δ*speG* mutant (Figure 2 C-H). For both OES2-0017 and OES1-1087, CMRSA-10 displayed a similar synergistic phenotype to that observed in the other wild-type USA300 strain, JE2 (Figure 2 I and J). The Δ*speG* mutant complemented with the low copy number plasmid pKK22 expressing *speG* under the control of the *fba* (fructose bisphosphate aldolase) promoter phenocopies wild-type USA300 carrying the control vector in checkerboard assays between spermine and OES2-0017 or OES1-1087 (Supplementary Fig. 5). These results further suggest that OES2-0017 and OES1-1087 are interfering with SpeG activity, and that strains must encode *speG* to observe the spermine synergistic phenotype.

Since SpeG is a spermine/spermidine acetyltransferase, we also tested spermidine with either OES2-0017 or OES1-1087 in checkerboard assays. As expected, OES2-0017 and spermidine synergized with an FICI of 0.33 ± 0.06 and Synergy Score of 19.08 ± 9.57, such synergy was decreased in the Δ*speG* mutant (FICI = 0.48 ± 0.06; Synergy Score = 12.26 ± 7.91) and *speG^-^*strains but not completely lost (Figure 2 K and L and Supplementary Fig. 6). The same is observed with spermidine and OES1-1087 where the Δ*speG* mutant (FICI = 0.38 ± 0.09; Synergy Score = 13.60 ± 1.93) or *speG^-^*strains showed reduced synergy compared to wild-type USA300 (FICI = 0.17 ± 0.03; Synergy Score = 22.77 ± 0.92) (Figure 2 M and N and Supplementary Fig. 6). The results suggest that synergy between these compounds and spermidine is mediated, at least in part, by SpeG inhibition.

### OES2-0017 inhibits SpeG resulting in spermine synergy

We sought to confirm SpeG inhibitory activity of OES2-0017 in an *in vitro* assay using purified His-tagged recombinant SpeG. We used a colorimetric SSAT assay based on the quantification of CoA-SH formed during the transfer of the acetyl group from AcCoA, which is proportional to the amount of the acetylated substrate (*N*-acetylspermine) ^54^. Under these experimental conditions, we observed acetyltransferase activity for the native and not the heat-inactivated SpeG when both spermine and spermidine were used as the test substrates (Supplementary Fig. 7). Based on our results and previous analysis by Tsimbalyuk *et al.* ^55^, we fit SpeG to a Michaelis-Menten and allosteric model with spermine as a substrate and an allosteric model with spermidine as the substrate and estimated catalytic efficiency of each as K_m_ and S_0.5_, respectively. The SpeG K_m_ with spermine fit to a Michaelis-Menten model was 1312 ± 348.1 μM and S_0.5_ with spermine fit to an allosteric model was 757.5 ± 63.35 μM (Supplementary Fig. 7). The S_0.5_ of spermidine was estimated as 6.078 ± 0.583 mM; however, testing spermidine concentrations higher than 10 mM was not achievable due to pH sensitivity of the assay and thus, the S_0.5_ estimated for spermidine may be an overestimation due to the curve not reaching a plateau (Supplementary Fig. 7). We also utilized high performance liquid chromatography (HPLC) to detect substrate depletion and product formation by quantifying spermine and *N*^1^-acetylspermine based off the method in ^56^. HPLC analysis of a 10-minute SpeG reaction with spermine and AcCoA (1 mM each) confirms SpeG spermine acetylation activity (Supplementary Fig. 7). We observed no activity when putrescine was tested as a substrate (Supplementary Fig. 7). These results agree with previous reports showing that spermine is the preferred substrate of SpeG, followed by spermidine and that SpeG does not acetylate putrescine ^23,55^.

Next, we tested if OES2-0017 could inhibit acetyltransferase activity of SpeG *in vitro,* first by performing colorimetric enzymatic inhibition assays testing a dilution series of the compound in the presence of either a single concentration or multiple concentrations of substrate estimating IC_50_ and K_i_, respectively. In these first set of assays, we incubated SpeG with OES2-0017 at room temperature for 30 min before adding the substrate and assay reagents to allow for equilibration; such pre-incubation may be needed for slow-binding inhibitors ^57^. The IC_50_ of OES2-0017 against SpeG was 34.82 ± 6.810 μM at ∼ 1 x K_m_ of spermine (1500 μM) and the K_i_ was 38.70 ± 6.55 μM when using a fixed saturating AcCoA concentration of 1 mM (Figure 3 A and Supplementary Fig 8). In both colorimetric and HPLC-based SSAT assays, SpeG did not acetylate the polyamine analog OES2-0017 (Supplementary Fig 9); hence, spermine remains the only substrate for SpeG in the enzymatic inhibition assays (i.e., when OES2-0017 is added to the reaction mixture) and thus the detected CoA-SH correlates to the level of spermine acetylation (or reduction thereof) in these assays. Notably, we observed reduced SpeG inhibitory activity by OES2-0017 when we performed the enzymatic assay without pre-incubating the enzyme with the inhibitor (Supplementary Fig 8), suggesting that OES2-0017 is likely binding the allosteric site, thus the K_i_ calculation is based on fitting the data to the noncompetitive inhibition model (Supplementary Fig. 8). Importantly, the estimated IC_50_ (34.82 ± 6.810 μM) and K_i_ (38.70 ± 6.55 μM) values are higher than expected when compared to the concentrations exhibiting spermine synergy interactions in whole-cell assays (0.6–5 μM; Figure 1 H). However, this could be attributed to the *in vitro* cell-free experimental conditions (i.e., buffer concentration, pH, temperature, etc.) that do not perfectly match the conditions *in vivo,* which may be more optimal for catalysis. Of note, we use a high buffer concentration (100 mM Tris-HCl) to prevent polyamine pH-mediated cleavage of Ellman’s reagent (the colorimetric enzymatic assay reagent), which is sensitive to high pH ^58^. We further confirmed the inhibitory activity of OES2-0017 against SpeG by detecting spermine, *N*^1^-acetylspermine, and OES2-0017 by HPLC. OES2-0017 at 300 μM shows a significant decrease in the amount of *N*^1^-acetylspermine produced by SpeG after a 10-minute reaction, without pre-incubation with the inhibitor, compared to a DMSO treated control (Figure 3 B and C and Supplementary Fig. 8). These data confirm that OES2-0017 inhibits SpeG, preventing detoxification of spermine and thus resulting in synergy between OES2-0017 and spermine (Figure 3 A-C and Supplementary Fig. 8).

**Figure 3.**
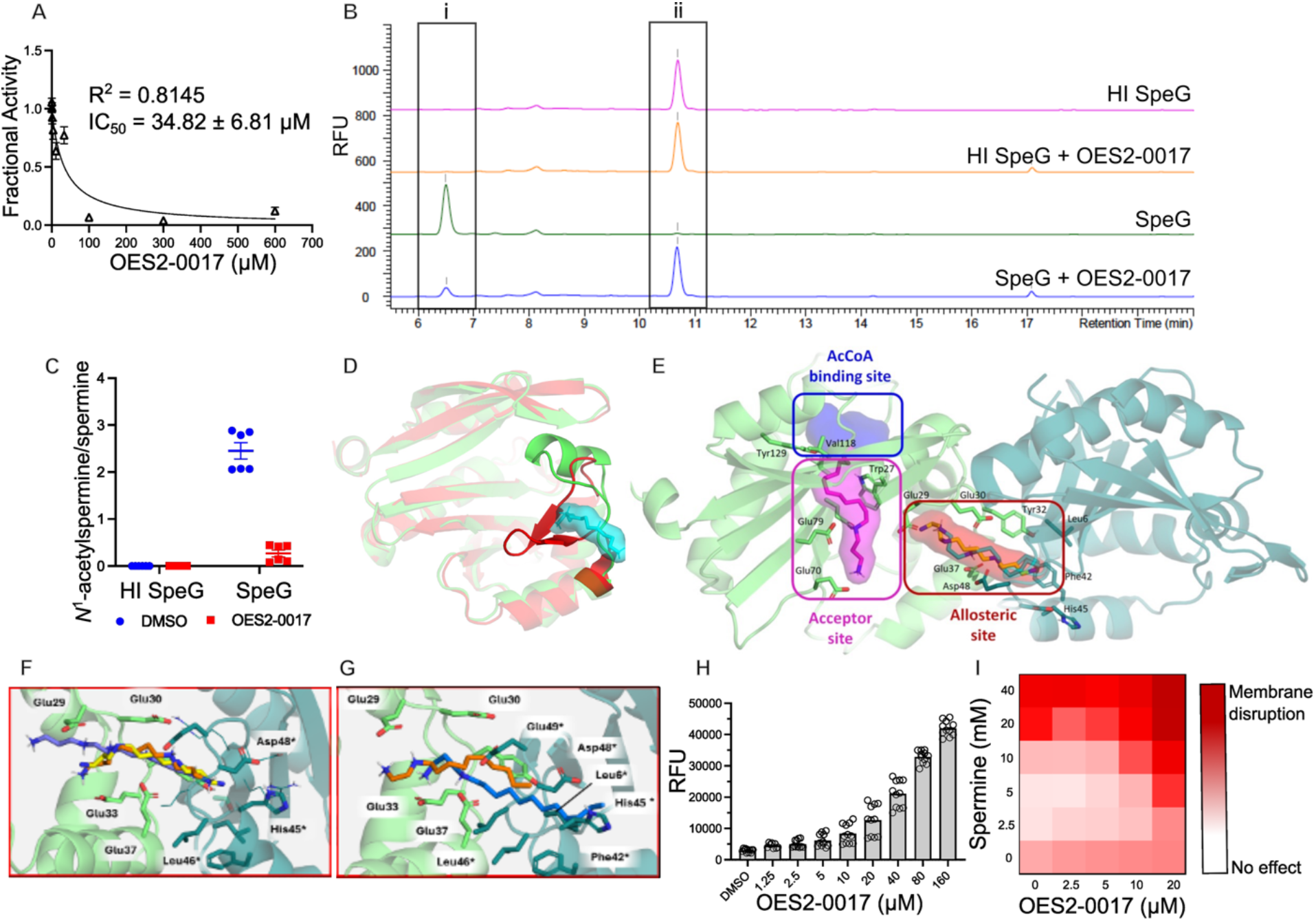
OES2-0017 directly inhibits SpeG and disrupts *S. aureus* membrane integrity. **A)** Colorimetric enzymatic inhibition assay conducted with 1 mM Acetyl Coenzyme A and 1500 μM spermine in the reaction mixture to evaluate the inhibition of OES2-0017 (n=7). Data is represented as mean ± SEM for seven independent experiments. **B)** Representative chromatograms of heat-inactivated SpeG, and SpeG, treated with 300 μM OES2-0017, or an equivalent volume of DMSO in a 10-minute reaction with 1 mM spermine and 1 mM AcCoA detecting **i** *N*^1^-acetylspermine and **ii** spermine. Primary amines were derivatized with *o*-phthalaldehyde (OPA) and *N*-acetyl-L-cysteine (NAC) and detected by fluorescence, represented in relative fluorescence units (λex 340nm, λem 450nm). **C)** Ratio of quantified *N*^1^-acetylspermine and spermine by HPLC analysis of the reaction represented in B. Results are grouped from two independent experiments reported as mean ± SEM, n=6. **D)** Superimposition of SpeG 3D structures, the apo (5ix3, red) and holo (8fv1, green) forms. The change in the conformation of the loop can be observed (amino acids from 21 to 23) from a disordered structure to a partial α-helix, allowing the spermine (cyan) to access to its binding site. **E)** General view of the SpeG dimer of *S. aureus* SpeG (PDB 8fv1). Monomers A and B are depicted in green and aquamarine blue, respectively. The allosteric site is marked with a red box, acceptor site with a magenta box, and AcCoA binding site with a blue box. Site surfaces are also represented with the same colours. The X-Ray crystallographic pose of spermine (PDB 8fv1) is depicted as yellow sticks; the predicted docked poses of OES2-0017 in the allosteric site is depicted as cyan sticks and in the acceptor site as magenta sticks. **F and G)** Selected binding poses obtained from docking calculations of spermine and OES2-0017 into SpeG dimer (PDB ID 8fv1). **F)** spermine (crystallographic bound pose of spermine shown in yellow as reference) and **G)** OES2-0017, bound inside the spermine binding site of the SpeG dimer in the docked pose (orange) and the pose after MD simulations (blue). **H)** DiSC_3_(5) membrane permeabilization assay of OES2-0017 against *S. aureus* USA300. Results represented as mean relative fluorescence units (RFU) ± SEM for three independent experiments (n=11). **I)** A representative DiSC_3_(5) membrane permeabilization assay performed as a mini-checkerboard evaluating the combined effects of OES2-0017 and spermine.

*S. aureus* SpeG forms a dodecameric structure composed of two hexamers stacked together in a ring-like structure ^55^. SpeG has two distinct polyamine binding sites, the allosteric site and active (acceptor) site ^59^, thus the dodecamer comprises a total of twelve allosteric and twelve active sites ^55,60^. Crystallographic data of SpeG from *S. aureus*, *Vibrio cholerae*, and *E. coli* show spermine and spermidine bound in an allosteric site formed by two adjacent monomers ^55,59–61^. When polyamines bind the allosteric site, a conformational change is observed where a mobile loop near the N-terminus of the protein becomes a more ordered alpha helix ^55,59,60^. Polyamines bound in the allosteric site are not acetylated since this site is not close enough to AcCoA for the acetyl transfer, rather another polyamine molecule binds the active site and is acetylated ^55,59,60,62^. Alignment of the crystal structures of *S. aureus* SpeG with and without bound spermine depicts this conformational change and the spermine binding site (PBD ID: 8fv1 and 5ix3, respectively; Figure 3 D).

To have a 3D perspective of the binding of OES2-0017 to SpeG, we carried out *in silico* studies combining docking calculations with MD simulations. OES2-0017 and spermine, as reference, were docked into the SpeG dimer from PDB 8fv1 (see Methods Section and Supplementary Table 3). The docking was performed into the allosteric site, which is placed between monomers A and B (Figure 3 E). OES2-0017 is predicted to bind with the N_1_-N_5_ diammonium group oriented towards the Glu29 (A) and Glu33 (A) carboxylate groups and the carbonyl group of Glu30 (A) backbone, establishing networks of ionic interactions and hydrogen bonds, and orienting the aliphatic chains towards the cavity of monomer B to establish hydrophobic interactions with aliphatic side chains of residues Leu6, Phe42, Leu46, Asp48, Glu51 and Arg53 (Figure 3 G). To confirm the stability of the docked poses, we performed molecular dynamics (MD) simulations of SpeG in complex with OES2-0017 and spermine as reference (see Methods, Figure 3 F and G and Supplementary Fig. 10). Both complexes remained stable with the ligands bound in the starting docked/crystallographic poses throughout the simulations. OES2-0017 maintained the aliphatic tail moving deeper into the non-polar cavity and reallocated the N_1_-N_5_ polar head towards Glu33 (A) and Glu37 (A) or Asp48 (B) carboxylate groups (Figure 3 G and Supplementary Fig. 10). The computational results show OES2-0017 binds in a similar orientation to spermine (Figure 3 F and G).

We also performed docking studies of OES2-0017 into the acceptor site of SpeG (PDB 8fv1) based on the observed binding of *N*^1^-acetylspermine into the acceptor site of *V. cholerae* SpeG ^59^. The resulting docked complexes showed a lower predicted binding energy (-9.57 kcal·mol^-1^) compared to the binding energy obtained in the allosteric site (-11.23 kcal·mol^-1^), suggesting that OES2-0017 may preferentially bind to the allosteric site rather than to the acceptor site (Supplementary Fig. 11). OES2-0017 established two ionic interactions of N_1_ with Glu70, N_5_ with Glu79, one hydrogen bond of N_5_ with the side chain of Gln81, a CH-π with the aromatic ring of Trp27, and several non-polar interactions between the aliphatic chain of the ligand with the side chains of Ile82 and Val118. For all 30 predicted docked poses, we observed that the polar head of OES2-0017 points outside instead of towards the catalytic Tyr129 while the aliphatic chain is oriented towards the bottom of the catalytic region and thus is incompatible with acetylation, which agrees with our experimental data showing the inability of SpeG to acetylate OES2-0017 (Supplementary Fig 9).

We then performed a thermal shift assay of the recombinant SpeG with OES2-0017 and spermine as a control. Ligands, including inhibitors, can bind to proteins within their native, non-native (such as an intermediate state), or unfolded states ^63,64^. Generally, when ligands bind to protein native states, T_m_ increases, and when bound to non-native states, T_m_ decreases ^63^. A titration with spermine increased SpeG T_m_ but only at the highest concentration tested (Supplementary Fig. 12), which is expected since spermine binding to the allosteric site leads to increased interactions of adjacent protomers ^62^. In contrast, OES2-0017 titration decreased SpeG T_m_ (Supplementary Fig. 12), indicating there is a binding interaction between OES2-0017 and SpeG but this interaction differs from that of spermine. The decrease in T_m_ suggests OES2-0017 binds preferentially to a non-native state of SpeG rather than the native state. Together, we hypothesize that OES2-0017 is binding to both the allosteric and active site, with preference to the allosteric site, although it is unclear whether OES2-0017 induces the conformational change to open the active (acceptor) site to substrate binding. However, given that SpeG has two binding sites for natural substrates ^59^ and that OES2-0017 was predicted to bind both sites, the mechanism of inhibition is likely more complicated than the standard competitive or noncompetitive inhibition models, which warrants further in-depth characterization. For this study, we fit data of OES2-0017 SpeG inhibition in the colorimetric assays with a noncompetitive inhibition model to align with our experimental assay design and results (of pre-incubation vs. no pre-incubation of enzyme and inhibitor discussed above; Figure 3 and Supplementary Fig. 8) to estimate approximate inhibition constants.

### OES2-0017 exhibits growth inhibitory effects via membrane perturbation

OES2-0017 exhibits growth-inhibitory effects at concentrations higher than those required to synergize with spermine; therefore, we hypothesized that this activity occurs via a second mode of action. Given its growth inhibitory activity at low micromolar range, we initially hypothesized this activity was mediated through an essential protein target. We confirmed activity against *Bacillus subtilis,* a Gram-positive organism with high degree of genetic conservation with *S. aureus* (Supplementary Fig. 13). Then, we screened a knock-down library covering the essential genes of *B. subtilis* against three sub-inhibitory concentrations of OES2-0017, spanning 2.5–10 μM or 1/16^th^–1/4^th^ the *B. subtilis* wild-type MIC (Supplementary Fig. 13). While some mutants showed reduced growth in the presence of one or more of the tested sub-inhibitory concentrations, follow-up dose-response assays revealed that none of those mutants was more susceptible to OES2-0017 relative to the wild-type, suggesting the absence of an essential protein target.

Next, we hypothesized that the growth-inhibitory activity of OES2-0017 is due to an effect on membrane integrity in the absence of an essential protein target. Using a DiSC_3_(5) membrane integrity assay ^65^, we found that OES2-0017 permeabilized the membrane of USA300 at concentrations starting from its MIC (≥ 20 μM) in a concentration-dependent manner (Figure 3 H and I), suggesting membrane activity at high concentrations as the mechanism of growth inhibitory effects of OES2-0017.

Next, we sought to check whether membrane disruption contributed to the observed synergy between OES2-0017 and spermine. We conducted a checkerboard-style DiSC_3_(5) assay with a combination of spermine and OES2-0017. OES2-0017 concentrations that correspond to the range of synergy (2.5-10 μM) against USA300 resulted in no observable permeabilization (Figure 3 I). Like OES2-0017, spermine disrupted membrane integrity of USA300 at concentrations starting from its MIC against USA300 (Figure 3 I and Supplementary Fig. 14). We observed similar membrane permeabilization with spermidine and putrescine (Supplementary Fig. 14). Large polyamine molecules were previously shown to permeabilize the outer membrane of Gram-negative bacteria ^66^; however, we found no reports on similar effects on Gram-positive membrane integrity. Importantly, there was no significant effect on membrane integrity in the presence of the combination of OES2-0017 and spermine at concentrations where synergy is observed (Figure 3 I), suggesting that membrane permeabilization does not significantly contribute to the observed synergy. Taken together, we revealed a dual mode of action of OES2-0017 with high concentrations leading to growth-inhibitory effects due to membrane perturbation and low concentrations synergizing with polyamines through inhibiting SSAT activity.

### OES1-1087 indirectly disrupts SpeG activity causing synergy with spermine

We took the same approach to uncover the mechanism of OES1-1087 activity by first testing direct inhibition of SpeG *in vitro* as described above. The IC_50_ of OES1-1087 could not accurately be estimated in the colorimetric enzymatic inhibition assays as the enzyme activity did not reach 100% inhibition but was reduced relative to the uninhibited control in a reaction containing ∼1 x K_m_ of AcCoA (0.75 mM) and ∼ 1 x K_m_ spermine (1500 μM; Figure 4 A and Supplementary Fig. 15). OES1-1087 did not exhibit inhibitory activity under a fixed AcCoA concentration and variable spermine concentrations; however, at a fixed spermine concentration and variable AcCoA, OES1-1087 treatment resulted in a slight reduction of SpeG activity (Supplementary Fig. 15). On the other hand, HPLC quantification of a 5-minute SpeG reaction pre-incubated with 3 mM OES1-1087 did not show any significant differences in the quantity of *N*^1^-acetylspermine detected compared to the untreated conditions (Figure 4 B and C and Supplementary Fig. 15). Given that the OES2-0017 concentration needed for complete inhibition of SpeG *in vitro* is roughly 100X higher than the lowest concentration where synergy is observed in whole cell assays, we hypothesize that a greater inhibitory effect may be observed for OES1-1087 *in vitro* at higher concentrations that would increase the inhibitor to enzyme ratio. However, solubility constraints preclude testing higher concentrations of OES1-1087.

**Figure 4.**
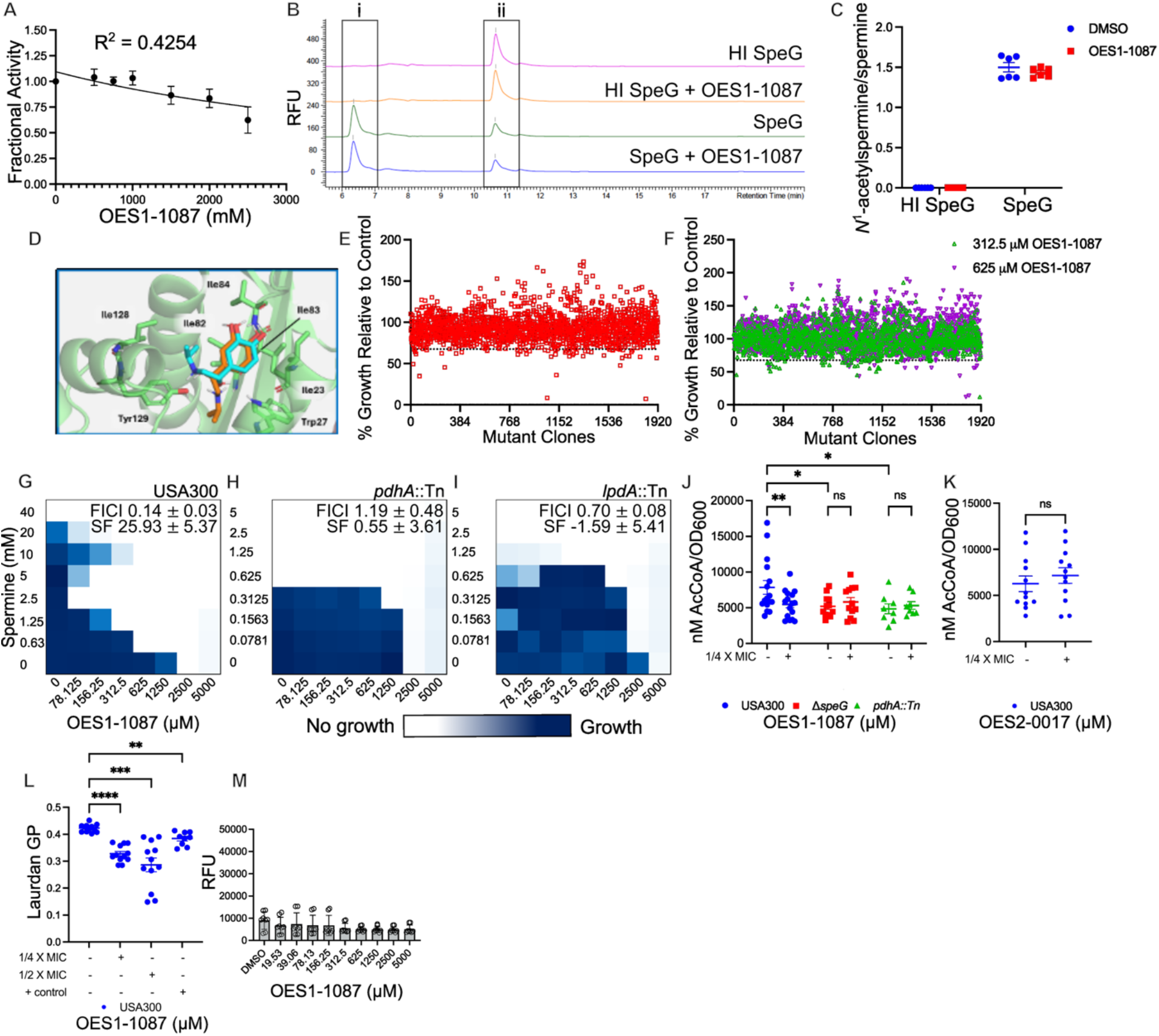
OES1-1087 reduces cellular AcCoA quantity and increases membrane fluidity. **A)**Colorimetric SpeG enzymatic inhibition assays conducted with 0.75 mM Acetyl Coenzyme A and 1500 μM spermine in the reaction mixture to evaluate the inhibition by OES1-1087 (n=3). Data is represented as mean ± SEM for three independent experiments. **B)** Representative HPLC chromatograms of heat-inactivated SpeG, and SpeG, pre-treated with 3 mM OES1-1087 or an equivalent volume of DMSO for 30 min before the addition of 1 mM spermine and 0.75 mM AcCoA substrates and incubating at 37°C for 10-min detecting **i** *N*^1^-acetylspermine and **ii** spermine. Primary amines were derivatized with OPA/NAC and detected by fluorescence, represented in relative fluorescence units (λex 340nm, λem 450nm). **C)** Ratio of quantified *N*^1^-acetylspermine and spermine by HPLC analysis of the reaction represented in B. Results are grouped from two independent experiments reported as mean ± SEM, n=6. **D)** Detail of the computationally predicted interactions of *R*-OES1-1087 and *S*-OES1-1087 (cyan and orange, respectively) docked into the AcCoA binding region of the SpeG dimer (PDB ID 8fv1). **E-F)** Chemogenomic screen of the Nebraska Transposon Mutant Library with **E)** a sub-inhibitory concentration of spermine (5 mM) and **F)** sub-inhibitory concentrations of OES1-1087 (312.5 µM and 625 µM). **G-I)** Representative checkerboard assays of OES1-1087 and spermine against *S. aureus* **G)** USA300 (n=14), **H)** *pdhA*::Tn (n=4), and **I)** *lpdA*::Tn (n=4). FICI values and Synergy Finder (SF) scores are reported as mean ± standard deviation of the indicated number of replicates. **J)** AcCoA quantification of cell lysates of USA300 (n=16), Δ*speG* (n=12), and *pdhA*::Tn (n=8) treated with 1/4 X MIC OES1-1087 (312.5 µM). Results are grouped from eight independent experiments and data represented as mean ± SEM. Statistical significance was determined by ordinary two-way ANOVA with Tukey’s multiple comparisons test. **K)** Acetyl-CoA quantification of USA300 cell lysates treated with 1/4 X MIC OES2-0017 (5 µM; n= 12). Results are grouped from six independent experiments and data represented as mean ± SEM. Statistical significance was determined by a paired t-test. **L)** Laurdan general permeabilization assay performed with USA300 cells treated with 1/4 X and 1/2 X MIC OES1-1087 (312.5 and 625 µM) or 50 mM benzyl alcohol as a positive control. Data is grouped from four independent experiments and represented as mean ± SEM, n=12, except for the positive control treatment where data is grouped from three independent experiments, n=9. Statistical significance was determined by mixed-effects analysis, with Geisser-Greenhouse correction and Dunnett’s multiple comparisons test. **M)** DiSC_3_(5) membrane permeabilization assay of OES1-1087 against *S. aureus* USA300. Results represented as mean ± SEM for three independent experiments (n=8). * represents p ≤ 0.05, ** p ≤ 0.01, *** p ≤ 0.001, **** p ≤ 0.0001, and ns p > 0.05.

We utilized *in silico* studies to predict if OES1-1087 can interact with SpeG resulting in the moderate SpeG-inhibition observed in the colorimetric enzymatic assay. Since OES1-1087 is not a polyamine analog, and rather more closely resembles the structure of AcCoA, we hypothesized it may be binding the same site as AcCoA. Therefore, docking calculations of the *R* and *S* enantiomers of OES1-1087 were performed into the AcCoA binding pocket (Supplementary Table 3). Both enantiomers were predicted to bind into the AcCoA binding pocket, through hydrogen bonds of the catechol hydroxyl groups with the oxygen of the backbone of Ile84 and orienting the isopropyl scaffold towards non-polar regions: the *S* enantiomer towards Ile82, Ile128 and Tyr129 sidechains, and the *R* enantiomer towards Trp28, and Tyr129 sidechains (Figure 4 D). These results suggest a possible competition with AcCoA, which agrees with the moderate effects of OES1-1087 observed in the colorimetric enzymatic assay when varying the concentration of AcCoA (Supplementary Fig. 15). We then tested OES1-1087 in a thermal shift assay and saw no observable effects on the T_m_ of SpeG (Supplementary Fig. 16). While this result may not demonstrate a OES1-1087-SpeG binding interaction within the tested concentration range, it does not preclude the possibility of binding that may occur at higher OES1-1087 concentrations not attainable under the experimental conditions of this assay or that may not induce T_m_ change given that some ligands bind to proteins with relatively high affinity but cause minor changes in T_m_ ^67^.

The whole-cell OES1-1087-spermine synergy phenotype requires SpeG (Figure 1 C), suggesting the inhibitor may be disrupting SpeG activity. Nonetheless, given the relatively modest SpeG inhibition by OES1-1087 in enzymatic assays (Figure 3 A and Supplementary Fig. 15), albeit supported by *in silico* predictions of occupying the AcCoA binding site (Figure 3 D), we sought to explore other mechanisms underlying the OES1-1087-synergy. To that end, we performed a genomic screen against the NTML ^50^ using sub-inhibitory concentrations of spermine (5 mM; less stringent than the screens shown in Figure 2 A) and OES1-1087 (312.5 or 625 μM), selected within the synergistic range. We hypothesized that a knock-out mutant of a gene whose product is involved in the spermine-OES1-1087 synergy will be more susceptible to one or both molecules but will not be further sensitized to the combination; i.e., lose the synergy. The screens identified a total of 69 mutants that showed growth reduction at one of the tested compound concentrations: 40 mutants from the 5 mM spermine screen that included *speG*::Tn (Figure 4 E) and 38 mutants from one or both OES1-1087 screens (Figure 4 F) with 9 mutants identified in both screens. We performed spermine-OES1-1087 mini-checkerboard assays against 69 mutants (Supplementary Fig. 17 and 18). Two mutants, *pdhA*::Tn and *lpdA*::Tn, showed increased spermine susceptibility and a potential loss of synergy (Supplementary Fig. 17). Follow-up full-sized checkerboard assays against both mutants confirmed the loss of synergy with FICI values of 1.19 ± 0.48 (*pdhA*::Tn) and 0.70 ± 0.08 (*lpdA*::Tn) compared to the wild-type FICI of 0.14 ± 0.03 (Figure 4 G-I). The loss of synergy and increased susceptibility to spermine were restored to the wild-type levels in complemented strains (Supplementary Fig. 19). Notably, we and others observed poor growth of these two mutants in their untreated growth controls ^68,69^, which may have hindered their identification from the more stringent spermine screens (Figure 2A). As such, all checkerboard assays against both mutants were performed with an inoculum 10-fold higher than normal. Both genes are part of the pyruvate dehydrogenase complex (PDHc); *pdhA* encodes the alpha subunit of the E1 component of the PDHc and *lpdA* encodes dihydrolipoamide dehydrogenase, the E3 component of the PDHc ^70^. Importantly, *S. aureus* USA300 produces two dihydrolipoamide dehydrogenase enzymes both denoted LpdA; however, herein, we refer to LpdA SAUSA300_0996, which is part of the PDHc. The PDHc converts pyruvate to AcCoA, and a *pdhA*::Tn mutant has impaired AcCoA production ^69^.

AcCoA is required for SpeG-mediated detoxification of polyamines, thus we hypothesized that OES1-1087 may be affecting the production of AcCoA leading to the loss of synergy observed in checkerboard assays with Δ*speG*, *pdhA*::Tn, and *lpdA*::Tn mutants. We quantified AcCoA content from cell lysates and observed a significant reduction (p = 0.0080) in AcCoA quantity in *S. aureus* USA300 cells treated with 1/4 X MIC OES1-1087 (312.5 μM; Figure 4 J). There were significant decreases in AcCoA levels of untreated cells of both Δ*speG* (p = 0.0167) and *pdhA*::Tn (p = 0.0171) mutants relative to the wild-type; however, there was no significant further reduction in the AcCoA content in either mutant upon treatment with 1/4 X MIC OES1-1087 (312.5 µM; Figure 4 J). The *pdhA*::Tn mutant is expected to have reduced levels of intracellular AcCoA due to disruption of the PDHc and we hypothesize that the reduction in AcCoA content in the Δ*speG* mutant is due to metabolic adaptations in response to the loss of the acetyltransferase. As a control, we quantified AcCoA from USA300 cell lysates treated with 1/4 X MIC OES2-0017 (5 µM) and observed no significant changes to intracellular AcCoA content (Figure 4 K). We then hypothesized that if OES1-1087 was acting on the same metabolic pathway to synergize with spermine as OES2-0017 then the two compounds should not synergize further in the presence of spermine. To test for this, we performed checkerboard assays with OES2-0017 and spermine and added OES1-1087 to the media in a 2-fold concentration gradient, effectively performing a 3D checkerboard. When titrating OES1-1087 into the background of OES2-0017-spermine checkerboards we did not observe further synergy suggesting that both compounds are acting in the same pathway (the SpeG-mediated acetylation) to disrupt spermine detoxification (Supplementary Fig. 20).

A *S. aureus pdhA*::Tn mutant has increased membrane fluidity ^69^. *S. aureus* strains with increased membrane fluidity are more susceptible to membrane active antimicrobials, such as daptomycin ^71^ and the cationic antimicrobial peptides LL37 and HNP-1 ^72^. This led us to hypothesize that OES1-1087 may affect cell membrane fluidity, thereby increasing susceptibility to spermine-mediated membrane perturbation. We performed a Laurdan assay to test membrane fluidity of *S. aureus* USA300 when treated with 1/4 and 1/2 X MIC of OES1-1087 (312.5 and 625 μM) and observed an increase in membrane fluidity in a dose-dependent manner (Figure 4 L). Notably, OES1-1087 did not show observable membrane perturbing activity in a DiSC_3_(5) assay (Figure 4 M). Membrane fluidity of *S. aureus* is dependent on the ratio of branched chain to straight chain fatty acids with a higher ratio resulting in increased membrane fluidity ^73,74^. Fatty acid production in *S. aureus* is reliant on the CoA thioester pool composition with higher proportions of acyl-CoAs derived from the branched chain amino acids leucine, isoleucine, and valine leading to increased production of branched-chain fatty acids, while initiation of fatty acid synthesis by AcCoA gives rise to straight chain fatty acids ^73^. Under limited AcCoA conditions, straight chain fatty acid production is reduced and the fluidity of the membrane increases, as observed for the *pdhA*::Tn mutant (Figure 4 L). Of note, OES1-1087 (Isoproterenol) treatment in eukaryotes disrupts fatty acid content including increasing the quantity of free fatty acids in serum ^75,76^ and decreasing fatty acid uptake and β-oxidation of fatty acids ^69,76,77^. Together, the OES1-1087-spermine synergy appears to be largely driven by a reduction in AcCoA content, impairing SpeG-mediated detoxification of spermine, with potential simultaneous contributions of the OES1-1087-mediated increased membrane fluidity that may exacerbate the membrane damaging effects of spermine and a potential modest direct SpeG inhibitory effect.

### Uncovering parallel polyamine acetyltransferase mechanisms in USA300

Checkerboard assays with OES2-0017 and OES1-1087 revealed only a partial loss of synergy in the Δ*speG* mutant with the polyamine spermidine (Figure 2 K-N and Supplementary Fig. 6), suggesting other potential mechanisms may also be at play in the spermidine synergy and that these may also be targeted by the inhibitors. SpeG preferentially acetylates spermine and acetylates spermidine with a lower affinity (Supplementary Fig. 7 and ^23,55^). The lower affinity may contribute to a decreased ability to detoxify spermidine, possibly requiring other detoxification mechanisms that function in parallel to SpeG. Some bacteria encode SSATs distinct from SpeG, such as *B. subtilis* encoding two enzymes by *paiA* and *bltD* ^78,79^ and *Enterococcus faecalis* encoding *pmvE* a homolog of *B. subtilis paiA* ^29,78,79^. The presence of both *paiA* and *bltD* in *B. subtilis* provides evidence for the existence of parallel polyamine detoxification pathways. Further, a recent report observed an accumulation of *N*-acetylputrescine in the MSSA strain ATCC 25923, which does not encode SpeG; however, the enzyme responsible for its production was not identified ^80^. This further suggests that *S. aureus* may encode alternative polyamine acetyltransferases and that they may have activity towards different polyamines. We identified potential additional polyamine acetyltransferases in *S. aureus* by performing protein blast of the *B. subtilis* 168 strain sequences of BltD (Accession NP_390537.1) and PaiA (Accession NP_391095.1) against *S. aureus* to identify potential homologs. Blastp results for BltD revealed two hypothetical proteins of *S. aureus* USA300 FPR3757; SAUSA300_0441 with 29.76 percent identity (query coverage 54%, E value of 2e^-^09) and SAUSA300_2083 with 27.41 percent identity (query coverage 86%, E value of 5e^-^09). PaiA blastp results identified a homolog of *S. aureus* USA300 FPR3757; SAUSA300_2316 (denoted PaiA_Sa_ herein), with 40.35 percent identity (99% query coverage, E value of 3 e^-^39). These *S. aureus* proteins have not been previously characterized functionally; all three are annotated as ribosomal-protein-alanine acetyltransferases whereas SAUSA300_2083 and SAUSA300_2316 (PaiA_Sa_) are also annotated as spermine/spermidine acetyltransferase blt and PaiA, respectively.

To confirm or rule out SSAT activity of SAUSA300_2083, SAUSA300_0441, and PaiA_Sa_, we overexpressed and purified these enzymes to assess their activity *in vitro*. We observed acetyltransferase activity of PaiA_Sa_ when spermine and spermidine are used as substrates but negligible activity with putrescine under the test conditions (Figure 5 A-C and Supplementary Fig. 21). HPLC analysis of a 1-hour PaiA_Sa_ reaction reveals the production of *N*^1^-acetylspermine when 1 mM spermine and AcCoA are used as substrates (Supplementary Fig. 21). Thus, these results suggest that PaiA_Sa_ is able to acetylate both spermine and spermidine. Notably, SAUSA300_2083 and SAUSA300_0441 did not exhibit acetyltransferase activity against the polyamines spermine, spermidine or putrescine under the tested conditions (Supplementary Fig. 22). These data suggest that PaiA_Sa_ is likely to function as a spermine/spermidine acetyltransferase.

**Figure 5.**
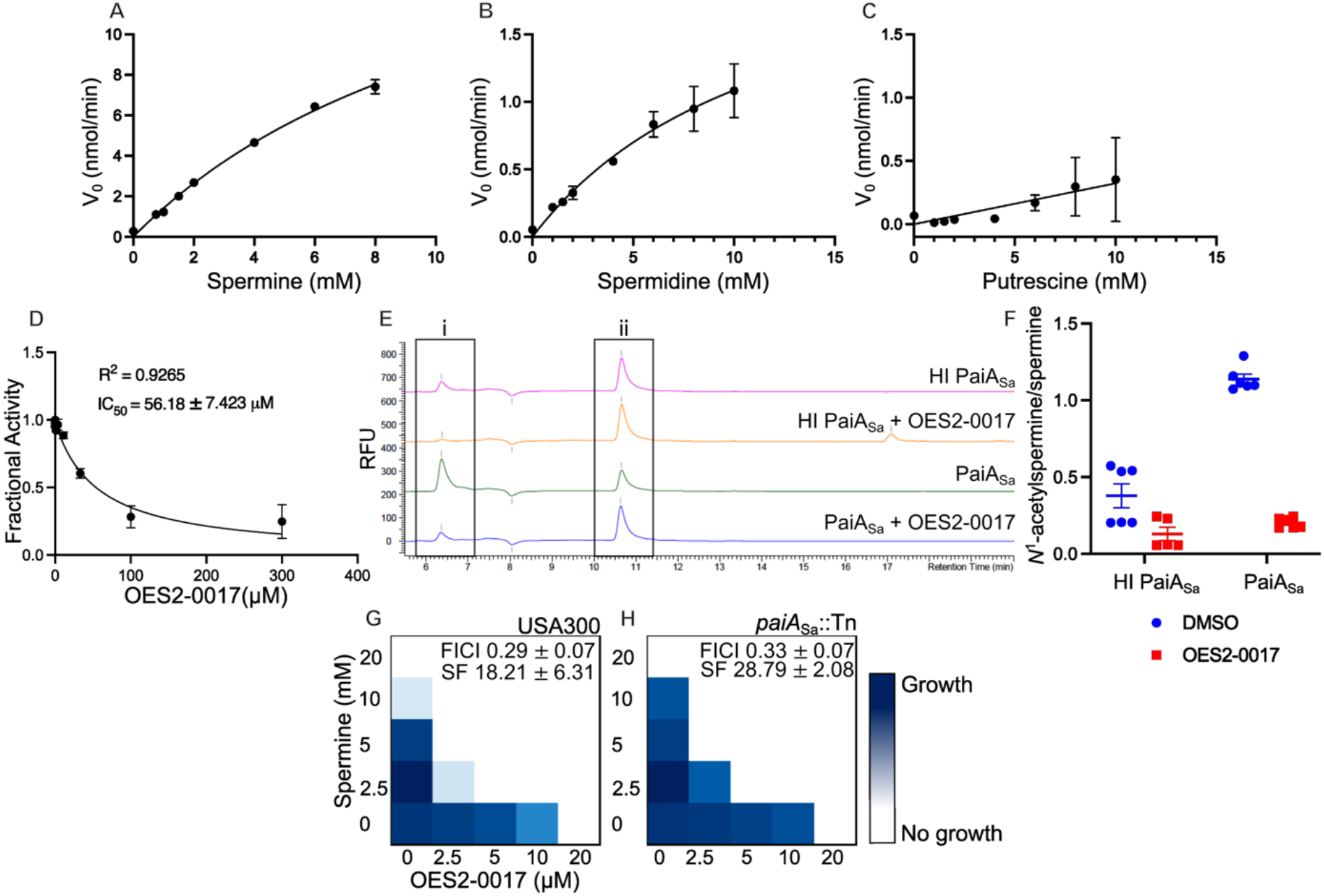
Interaction of polyamines and OES2-0017 uncovers PaiA_Sa_ as a spermine/spermidine acetyltransferase in *S. aureus*. **A-C)** Steady-state enzymatic assays of PaiA_Sa_ evaluate the acetylation activity of PaiA_Sa_. Each reaction contains 3.2 μg of PaiA_Sa_ 1 mM AcCoA and variable concentrations of substrates. Results are represented as a mean ± SEM of two independent experiments **A)** spermine (n=4), **B)** spermidine (n=2), and **C)** putrescine (n=2). **D)** Enzymatic inhibition assay of PaiA_Sa_ containing 0.8 μg of PaiA_Sa_, 2 mM spermine and 1 mM AcCoA and variable concentrations of OES2-0017 (n=3). Results are represented as a mean ± SEM of three independent experiments. **E)** Representative HPLC chromatograms of reactions containing heat-inactivated PaiA_Sa_ and PaiA_Sa_ with and without 1.2 mM OES2-0017 and 1 mM each AcCoA and spermine monitoring i, *N*^1^-acetylspermine and ii spermine. Primary amines were derivatized with OPA/NAC and detected by fluorescence, represented in relative fluorescence units (ex 340nm/em 450nm). **F)** Ratio of quantified *N*^1^-acetylspermine and spermine by HPLC analysis of a 1-hour reaction of PaiA_Sa_ and heat inactivated PaiA_Sa_ with 1 mM AcCoA and spermine and 1.2 mM OES2-0017. Results are represented as a mean ± SEM of two independent experiments n=6. **G-H**) Representative checkerboard assays of spermine and OES2-0017 against *S. aureus* **G)** USA300 (n=16), **H)** *paiA*::Tn (n=3). FICI values and Synergy Finder (SF) scores are reported as mean ± standard deviation.

Next, we tested the ability of PaiA_Sa_ to acetylate OES2-0017, being a polyamine analog. Using both a colorimetric enzymatic assay and HPLC, we observed that PaiA_Sa_ acetylates OES2-0017 (Supplementary Fig. 23). We then tested if OES2-0017 could inhibit PaiA_Sa_ acetylation of spermine; the low spermidine acetylation signal precluded testing OES2-0017 PaiA_Sa_ inhibition with spermidine as a substrate. OES2-0017 inhibited PaiA_Sa_ activity exhibiting an IC_50_ value of 56.18 ± 7.423 μM and estimated K_i_ of 43.83 ± 6.693 μM (Figure 5 D Supplementary Fig. 24). HPLC data similarly show a significant reduction in the quantity of *N*^1^-acetylspermine detected in a 1-hour reaction containing 1 mM spermine and AcCoA when PaiA_Sa_ was treated with 1.2 mM OES2-0017 without pre-incubation with inhibitor, relative to DMSO vehicle treated enzyme (Figure 5 E and F and Supplementary Fig. 24). We hypothesize that since PaiA_Sa_ can acetylate both OES2-0017 and spermine, both substrates compete for the enzyme, resulting in reduced spermine acetylation by PaiA_Sa_. Next, we performed whole-cell spermine-OES2-0017 mini-checkerboard assays against a *paiA*::Tn mutant. Compared to the wild-type USA300, no loss of synergy in the *paiA_Sa_* mutant was observed (Figure 5 G and H). SpeG, hypothesized to be the primary spermine/spermidine detoxification enzyme in *S. aureus*, may have compensated for the absence of PaiA_Sa_, masking any phenotypic changes in the *paiA_Sa_* mutant; a *speG paiA_Sa_* double mutant may be required to assess the contribution of PaiA_Sa_ in whole cells. Further, the lack of increased sensitization of *paiA*::Tn to OES2-0017 suggests it may be inefficient at acetylating OES2-0017 in whole cells, as its inhibitory activity remained the same regardless of an active PaiA_Sa_. Together, our results indicate that polyamine acetylation in *S. aureus* may occur through PaiA_Sa_ in addition to SpeG, serving as complementary acetyltransferase enzymes, and that the SSAT function of both enzymes is inhibited by OES2-0017.

### Broad-spectrum activity of OES2-0017 and OES1-1087 against clinically relevant bacteria and *Candida*

We conducted a BLAST search of spermine/spermidine acetyltransferases using *S. aureus* SpeG as the query, obtaining 94 amino acid sequences comprising eukaryotes, bacteria, and archaea. Sequences were aligned using MUSCLE and a pairwise deletion of ambiguous positions was performed for each sequence pair. The conservation of various SSATs was determined by a maximum likelihood phylogenetic analysis (Supplementary Fig. 25, Supplementary Table 4). Although the main purpose of such an analysis is to infer evolutionary history and relatedness, we used the phylogenetic tree to visualize or infer sequence similarity between known spermine/spermidine acetyltransferases. Our analysis reveals two main branches originating from a common ancestor (Supplementary Fig. 25, labelled 1, 2a, and 2b). Branch 1 shows archaea and (vertebrate) eukaryotic organisms clustering together. This branch includes the human SAT1, rat, and other mammalian SATs. The same branch also included *Acinetobacter baumannii* Dpa, which was recently characterized as more closely resembling eukaryotic SSATs than bacterial ones ^81^. The second branch was divided into two sub-branches; branch 2a contained SpeG homologs in fungi and yeast while branch 2b comprised bacterial SpeG. Branch 2b included SpeG and other polyamine acetyltransferases that have previously been characterized in both Gram-positive and Gram-negative organisms including *Staphylococcus epidermidis* ^25^*, Escherichia coli* ^82^*, V. cholerae* ^60^*, Yersinia pestis* (PDB ID: 6d72), and *Salmonella enterica* serovar Typhimurium ^28^.

Given that SpeG has been implicated in virulence phenotypes in multiple bacteria ^13,28,83^, we tested the identified SpeG inhibitor OES2-0017 against a broad range of Gram-positive and Gram-negative bacteria and the pathogenic yeast *Candida albicans* in checkerboard assays with spermine. The combination of OES2-0017 and spermine exhibited a synergistic interaction or an increase in spermine susceptibility in all tested organisms, namely *B. subtilis, E. faecalis, E. faecium, S. epidermidis, M. smegmatis, B. cenocepacia, E. coli, K. pneumoniae, S.* Typhimurium, and *C. albicans* (Figure 6 A-K). In *A. baumannii* the polyamine acetyltransferase Dpa preferentially acetylates 1,3-diaminopropane ^81^, which is a diamine; therefore, we tested the diamine putrescine and OES2-0017 in combination. OES2-0017 synergized with putrescine against *A. baumannii* (Figure 6 L). As in *S. aureus*, OES2-0017 also inhibited the growth of all tested bacteria and *C. albicans* at the micromolar range (Figure 6). Next, we tested the combination of spermine and OES2-0017 against the respective SSAT mutants of *E. coli* BW25113, *K. pneumoniae* MKP103, *S. enterica* sv. Typhimurium 14028s, and *B. subtilis* 168 (Figure 6 M-P). OES2-0017 no longer potentiated the effects of spermine in the SSAT mutants as opposed to their respective wild-type strains (Figure 6 M-P), suggesting OES2-0017 mediates its broad-spectrum spermine synergy through SSAT inhibition. Similarly, we tested the OES1-1087–spermine combination against other Gram-positive and Gram-negative bacteria. As observed in *S. aureus,* OES1-1087 synergized spermine against *S.* Typhimurium, *K. pneumoniae,* and *S. epidermidis* and the synergistic phenotype is lost in the respective *speG* mutants in *S.* Typhimurium and *K. pneumoniae* and the pyruvate dehydrogenase mutant, *aceE*::Tn30, in *K. pneumoniae* (Figure 6 Q-V). This is consistent with the phenotype observed in *S. aureus* and suggests OES1-1087 is inhibiting polyamine detoxification via a similar mechanism in other bacteria. Together, OES2-0017 and OES1-1087 exhibited broad-spectrum polyamine synergy, including against clinically relevant World Health Organization priority pathogens ^84^.

**Figure 6.**
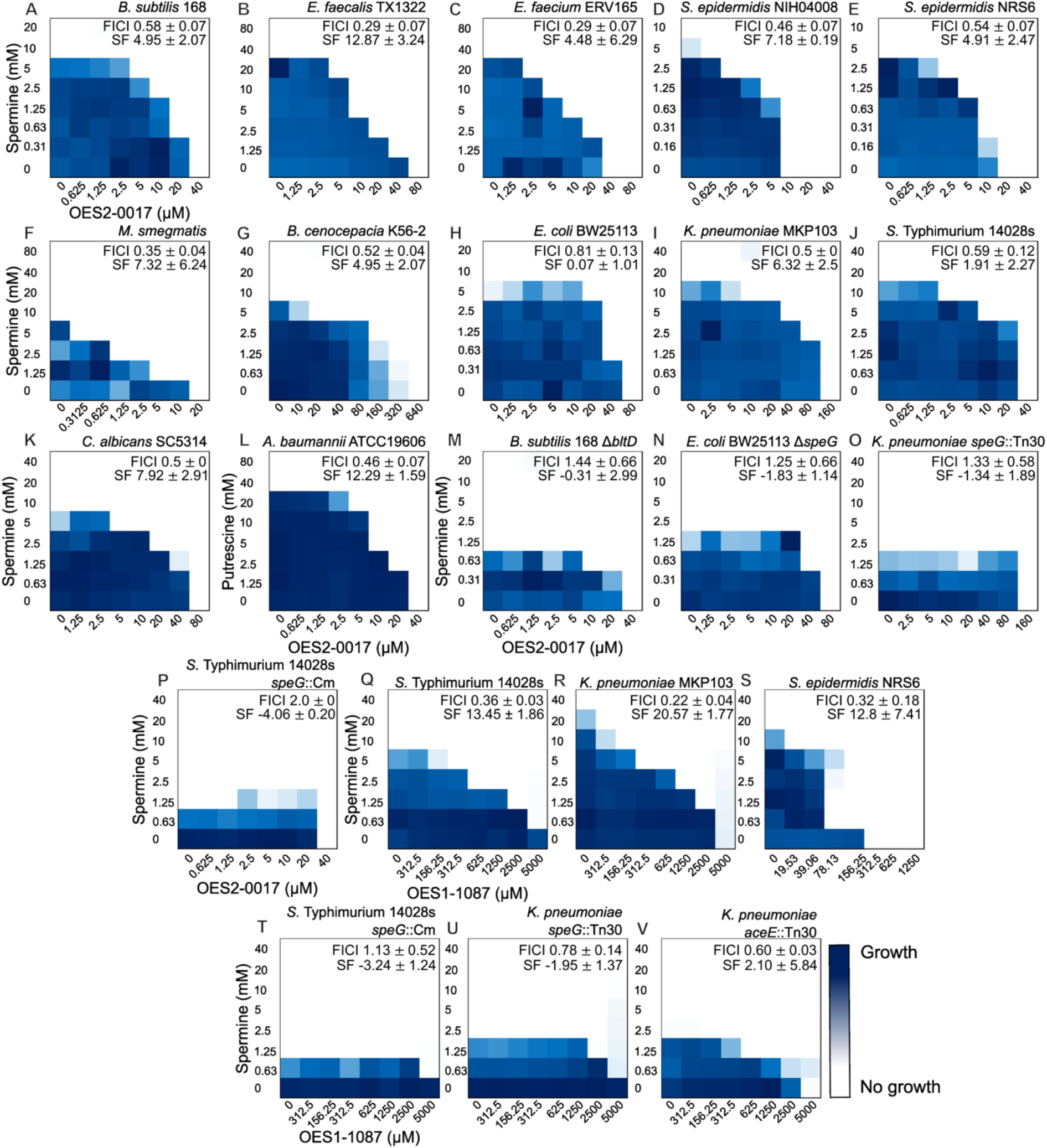
OES2-0017 and OES1-1087 exhibit broad-spectrum activity. Representative checkerboard assays of OES2-0017 and spermine against **A)** *B. subtilis* 168 (n=3), **B)** *E. faecalis* TX1322 (n=3), **C)** *E. faecium* ERV165 (n=3), **D)** *S. epidermidis* NIH04008 (n=3), **E)** *S. epidermidis* NRS6 (n=3), **F)** *M. smegmatis* (n=3), **G)** *B. cenocepacia* K56-2 (n=3), **H)** *E. coli* BW25113 (n=4), **I)** *K. pneumoniae* MKP103 (n=4), **J)** *S.* Typhimurium 14028s (n=4), and **K)** *C. albicans* SC5314 (n=3), **L)** Putrescine and OES2-0017 checkerboard against *A. baumannii* ATCC19606 (n=3). Representative checkerboard assays of spermine and OES2-0017 against spermine/spermidine acetyltransferase mutants **M)** *B. subtilis* Δ*bltD* (n=4), **N)** *E. coli* Δ*speG* (n=3), **O)** *K. pneumoniae speG*::Tn30 (n=3), **P)** *S.* Typhimurium *speG*::Cm (n=3). Representative checkerboard assays of OES1-1087 and spermine against **Q)** *S.* Typhimurium 14028s (n=4), **R)** *K. pneumoniae* MKP103 (n=7), and **S)** *S. epidermidis* NRS6 (n=3). Representative checkerboard assays of spermine and OES1-1087 against spermine/spermidine acetyltransferase or pyruvate dehydrogenase mutants **T)** *S.* Typhimurium 14028s *speG*::Cm (n=4), **U)** *K. pneumoniae speG*::Tn30 (n=4), and **V)** *K. pneumoniae aceE*::Tn30 (n=4). FICI values and synergy scores are represented as mean ± standard deviation for the indicated number of replicates.

### Polyamine-mediated altered antibiotic susceptibility and the potential of OE2-0017 as an antibiotic adjuvant

Checkerboard assays of polyamines and representative antibiotics from seven classes with diverse cellular targets revealed various polyamine-mediated altered antibiotic responses (Supplementary Fig. 26). The three tested polyamines synergized with the macrolide azithromycin while spermine synergized with the β-lactam cefuroxime (Supplementary Fig. 26). The spermine β-lactam synergy matches previous reports of the phenotype in *S. aureus* ^30,32^. Similarly, polyamine synergy with macrolides has been reported in the Gram-negative pathogens *K. pneumoniae* ^34^ and *E. coli* ^32^. In *K. pneumoniae,* the synergistic interaction between azithromycin and putrescine is mediated through the combined effects of putrescine-induced membrane perturbation and protein synthesis inhibition via interaction at the ribosome ^34^. The effects of polyamines on ribosome function are well documented ^85^ and polyamines are known to interact with biological macromolecules, including DNA, RNA, proteins, and phospholipids ^18,86^. In contrast, the three polyamines antagonized rifampicin and kanamycin whereas spermine antagonized with vancomycin, defined as a FICI value ≥ 4 (Supplementary Fig. 26). The mechanisms of the observed polyamine-mediated alterations to antibiotic susceptibility warrant further investigation. On the other hand, we observed no alterations in daptomycin or ciprofloxacin susceptibility in combination with the tested polyamines (Supplementary Fig. 26).

Interestingly, the MIC of vancomycin is increased 4-fold when combined with spermine (FICI = 4.69 ± 1.57), but the interaction is lost in a Δ*speG* mutant (FICI = 2.14 ± 0.24), suggesting that *S. aureus* requires a functional SpeG to use exogenous polyamines to resist vancomycin (Figure 7 A and B). Supplementing a sub-inhibitory concentration of OES2-0017 (2.5 μM) to a vancomycin-spermine checkerboard against *S. aureus* USA300 phenocopied the loss of antagonism observed against the Δ*speG* mutant (FICI = 1.59 ± 0.62; Figure 7 C). This phenotype is not due to a synergistic interaction between OES2-0017 and vancomycin (Supplementary Fig. 27). Vancomycin is one of the first-line antibiotics for the treatment of severe MRSA infections ^87^, suggesting the antagonism between polyamines and vancomycin may be detrimental to MRSA treatment. Similarly, the polyamine antagonistic interactions with rifampicin and kanamycin in USA300 were lost in the Δ*speG* mutant and with supplementation of a sub-inhibitory concentration of OES2-0017 to the USA300 checkerboard (Figure 7 D-I). The loss of antagonism was not due to synergy between OES2-0017 and kanamycin or rifampicin (Supplementary Fig. 27). While kanamycin and rifampicin are not considered first-line monotherapies for MRSA infections, they are often used in conjunction with vancomycin ^88,89^. We also identified a synergistic interaction between OES2-0017 and cefuroxime, where OES2-0017 lowered the MIC of the antibiotic to <16 μg/mL from 1024 μg/mL (FICI = 0.35 ± 0.13; Figure 7 J). OES2-0017 and azithromycin also exhibited a slight potentiation effect against USA300 (FICI = 0.72 ± 0.21; Figure 7 K). OES2-0017 similarly synergized with azithromycin against *K. pneumoniae* ^34^. These results suggest that OES2-0017 may serve as an antibiotic adjuvant by preventing *S. aureus* from utilizing exogenous polyamines to resist antibiotics (e.g., glycopeptides, aminoglycosides, and rifamycins) or by potentiating the effects of antibiotics phenocopying the polyamine-mediated sensitization to certain antibiotics (e.g., macrolides and β-lactams).

**Figure 7.**
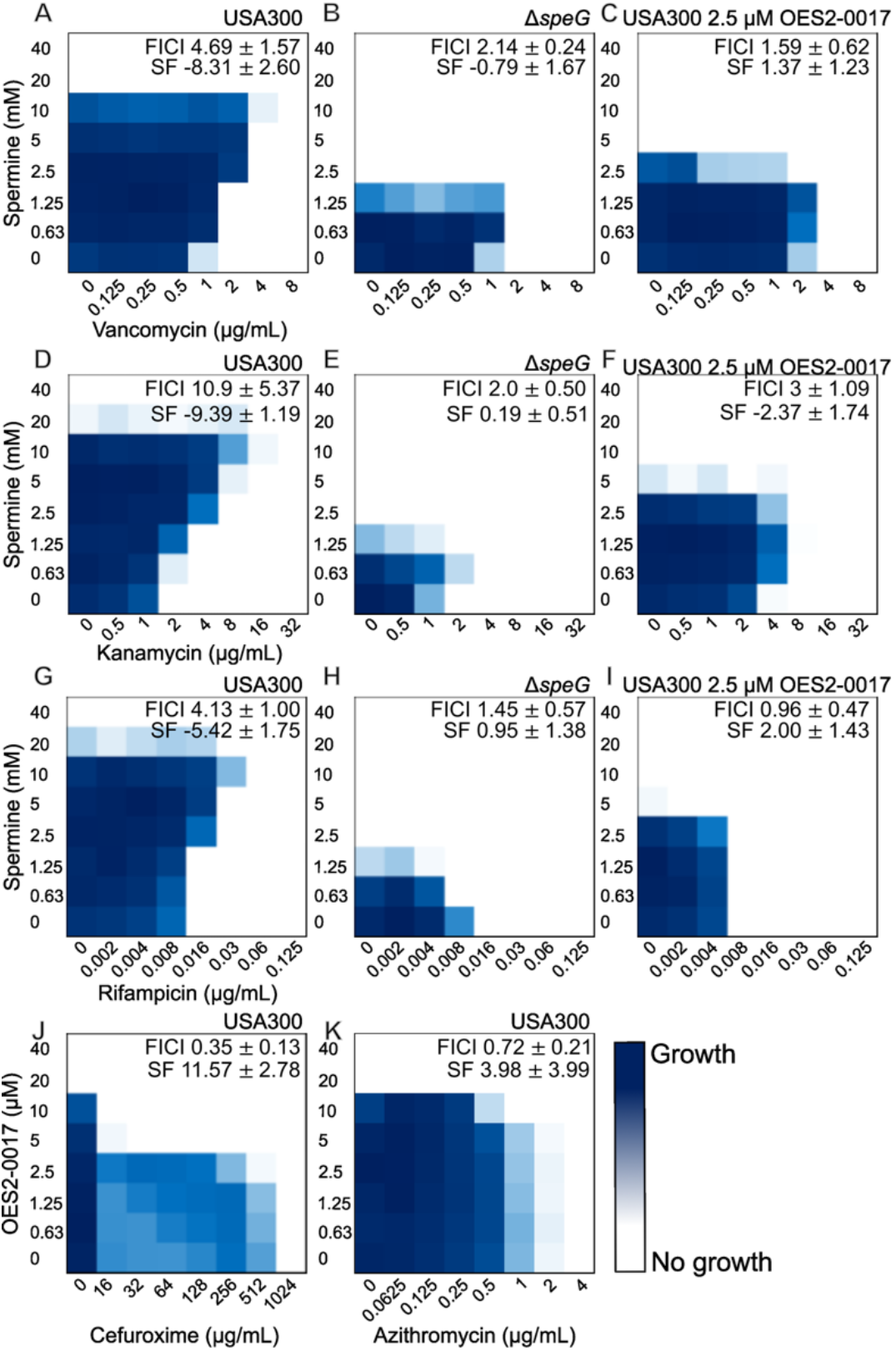
The potential of OES2-0017 as an antibiotic adjuvant. Representative checkerboard assays of spermine and vancomycin **(A-C)**, kanamycin **(D-F)**, and rifampicin **(G-I)** against *S. aureus* **A, D, G)** USA300, **B, E, H)** Δ*speG*, and **C, F, I)** USA300 supplemented with a sub-inhibitory concentration of OES2-0017 where OES2-0017 abolishes the protective effect of polyamines from the antibiotics. Representative checkerboard assay of OES2-0017 and cefuroxime **(J)** and azithromycin **(K)** against *S. aureus* USA300, where OES2-0017 potentiates the antibiotic activity. FICI values and Synergy Finder (SF) scores are represented as mean ± standard deviation for the indicated number of replicates: **A)** n=10, **B)** n=7, **C)** n=4, **D)** n=5, **E)** n=5, **F)** n=3, **G)** n=8, **H)** n=5, **I)** n=3, **J)** n=3, and **K)** n=4.

### The potential of OES2-0017 and OES1-1087 as antivirulence agents: *In cellulo* and *in vivo* activities

We tested the ability of OES2-0017 to clear *S.* Typhimurium infection of murine macrophages since SpeG was previously shown to be necessary for the intracellular replication of *S.* Typhimurium, where a *speG* knockout was attenuated in macrophages ^28^. Bacterial load was significantly reduced in a gentamicin protection assay at both 4-and 15-hours post-infection in the *speG::*Cm infected macrophages compared to the wild-type (p = 0.0286 and 0.0286, respectively; Figure 8 A and B). OES2-0017 treatment, whether pre-or post-infection, reduced *S.* Typhimurium bacterial load in macrophages, similar to the levels of recovered bacteria in the *speG*::Cm condition (Figure 8 A and B). Thus, OES2-0017 eradicated intracellular macrophage infection of the wild-type *S.* Typhimurium reducing its virulence.

**Figure 8.**
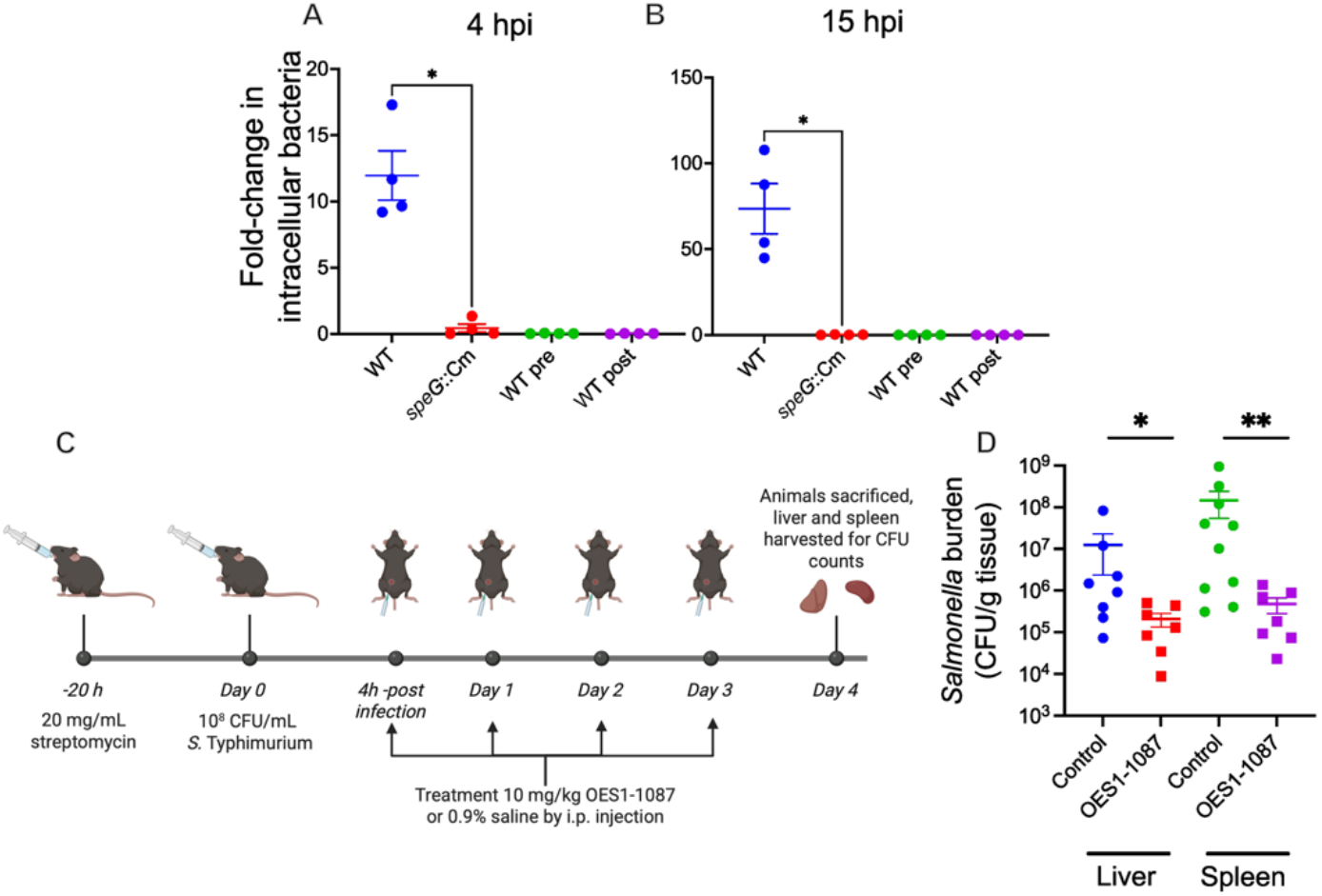
Antivirulence activities against *S.* Typhimurium: OES2-0017 abrogates bacterial intracellular survival and OES1-1087 reduces bacterial burden *in vivo*. A and **B)** murine macrophages treated with OES2-0017 before or after infection with *S.* Typhimurium 14028s or *speG*::Cm and bacterial colony forming units counted at **A)** 4 hours post-infection and **B)** 15 hours post-infection. Data represent 3 independent experiments, and statistical analysis was performed by Mann-Whitney test. **C)** Infection and treatment timeline of the murine *in vivo* experiment and **D)** the bacterial burden (CFU/g tissue) of female C57BL/6J mice 4 days post-infection that were orally infected with 10^8^ CFU/mL *Salmonella enterica* serovar Typhimurium 14028s and treated daily with 10 mg/kg OES1-1087 or saline (0.9% NaCl) i.p. Liver and spleen tissues were harvested and plated on LB agar for bacterial counting and PCR performed with specific *ssrB* primers were used to confirm the recovered bacteria was *S.* Typhimurium. Data represents two independent experiments each consisting of 5 animals per condition. Outliers were removed using Grubb’s test and statistical analysis was performed by Mann-Whitney test in GraphPad Prism 11.0 (* signifies p < 0.05 and ** p < 0.01). Created in BioRender. El-halfawy, O. (2026) https://BioRender.com/404qyln

We also tested the ability of OES1-1087 to act as a stand-alone antimicrobial *in vivo*. OES1-1087 (isoproterenol) is a previously approved drug with established dosage and safety profiles. A murine gastrointestinal infection with *S.* Typhimurium 14028s showed a significant reduction in bacterial burden in the liver (p=0.0401) and spleen (p=0.0046) following treatment with 10 mg/kg (i.p.) OES1-1087 relative to animals treated with a vehicle control (Figure 8 C and D). Of note, OES1-1087 did not exhibit growth inhibitory effects against *S.* Typhimurium *in vitro*, even at the highest tested concentration (5000 µM; Figure 6 Q). Together, this illustrates the potential of OES1-1087 as an antivirulence agent *in vivo*, making it a good candidate for drug repurposing and future SAR studies to further enhance its antibacterial and antivirulence properties.

### Evaluation of OES2-0017 and OES1-1087 potential eukaryotic activity

Given the membrane activity of OES2-0017 against *S. aureus* (Figure 3 H and I), we sought to evaluate its potential effects on mammalian cell membranes. To that end, we assessed the hemolytic activity of OES2-0017 against sheep red blood cells. OES2-0017 exhibited hemolytic effects, which are first observed at 160 μM with an increase at 640 μM reaching an absorbance of roughly 1/2 that of the positive control (Triton X100) (Supplementary Fig. 28). The hemolytic effects occur at a concentration significantly higher than those required to inhibit bacterial growth (8-fold higher than the MIC) or to synergize with spermine through SpeG inhibition (512-fold higher than the lowest point of synergy with spermine). Notably, OES1-1087 did not exhibit hemolytic effects at the concentrations tested (Supplementary Fig. 28). To further evaluate the relative toxicity of OES2-0017 and OES1-1087 toward eukaryotic cells, we exposed HepG2 cells to the compounds at concentrations equal to their MIC against *S. aureus*, which are already higher than their respective spermine synergizing concentrations, and evaluated cell damage via a lactate dehydrogenase (LDH) assay. Neither compound exhibited significant cytotoxicity towards the eukaryotic cells compared to a DMSO-treated control with both showing < 20% LDH release (Supplementary Fig. 29). Altogether, these data indicate that at the concentrations employed in our study significant toxicity towards host cells would not occur.

Next, we assessed the activity of OES2-0017 and OES1-1087 against the human spermine/spermidine acetyltransferase, SAT1. SAT1 is a dimer in solution with one monomer seemingly self-acetylating a lysine residue and the other binding polyamines for acetylation ^90^. SpeG and SAT1 are located on two separate branches of the phylogenetic analysis (Supplementary Fig. 25), suggesting a distant relation. First, we overexpressed and purified SAT1 enzyme and confirmed its acetyltransferase activity *in vitro* with spermine. SAT1 demonstrated higher substrate specificity (K_M_ = 73.10 ± 8.328 μM; Supplementary Fig. 30) towards spermine than SpeG (K_M_ = 1312 ± 348.1 μM; Supplementary Fig. 7), in agreement with previous reports ^23,91^. We then tested the ability of OES2-0017 and OES1-1087 to inhibit SAT1 utilizing the above-described colorimetric and HPLC enzymatic assays. The IC_50_ of OES2-0017 against SAT1 was 26.39 ± 6.411 μM and a significant reduction in *N*^1^-acetylspermine concentration was detected in a 10-minute reaction containing 1 mM spermine and AcCoA without pre-incubating SAT1 with 300 μM OES2-0017 (Supplementary Fig. 30). We also tested whether SAT1 could acetylate OES2-0017 and observed no acetylation activity towards OES2-0017 when used as a substrate (Supplementary Fig. 30). OES1-1087 did not inhibit SAT1 under any of the tested conditions (Supplementary Fig. 30). Although, OES2-0017 can inhibit SAT1 activity, this may not prove detrimental to human cells. Multiple studies have employed SSAT knockout mice and the non-specific SSAT inhibitor Berenil in *in vivo* mouse models, observing no toxic effects to mice ^92–94^. On the other hand, SAT1 inhibition might have other potential therapeutic uses. Altered polyamine metabolism has been implicated in many human diseases and conditions, including cancer, osteoporosis, and ischemia/reperfusion leading to acute kidney injury ^92,94–97^ with SAT1 being previously suggested as a promising drug target ^90,98^.

### Small-scale structure activity relationship of OES2-0017 suggests the structural components necessary for activity

We performed a small-scale structure activity relationship (SAR) study of OES2-0017 by testing analogs thereof to identify the active moiety of OES2-0017 and detect preferential inhibitory activity against either SpeG or SAT1. The analyses included compounds with varied length of the saturated alkyl chain R_1_ and one of three R_2_ (a proton, an aminoethyl group, or an aminopropyl group) attached on either side of an amine group (Table 1 and Supplementary Fig. 31). First, we tested each of the compounds in whole-cell assays in combination with spermine to assess their growth inhibitory and polyamine synergy effects against *S. aureus* USA300. We observed a general reduction in the MIC of the compound as the length of R_1_ and R_2_ increased (Table 1). Spermine synergy appeared to be dependent on both the length of the alkyl chain and the addition of either the aminoethyl or aminopropyl group. The FICI value decreased (indicating a greater degree of synergy) as the length of the alkyl chain increased; for example, OES2-0047 with R_1_ as a proton exhibited no synergy (FICI = 0.88 ± 0.18, Synergy Score = 2.74 ± 0.39) whereas OES2-0052 with R_1_ (CH_2_)_5_CH_3_ showed synergy (FICI = 0.50 ± 0, Synergy Score = 6.55 ± 1.62). The synergistic interaction is further modestly improved when the primary amine is changed to a secondary one with an aminoethyl or aminopropyl; this trend (mostly detectable by FICI) is exemplified in OES2-0046 (R_2_ = H; FICI = 0.58 ± 0.14, Synergy Score = 7.91 ± 2.21), OES2-0085 (R_2_ = aminoethyl; FICI = 0.56 ± 0.06, Synergy Score = 3.89 ± 2.55), and OES2-0045 (R_2_ = aminopropyl; FICI= 0.38 ± 0, Synergy Score 5.64 ± 0.84; Table 1).

**Table 1.**
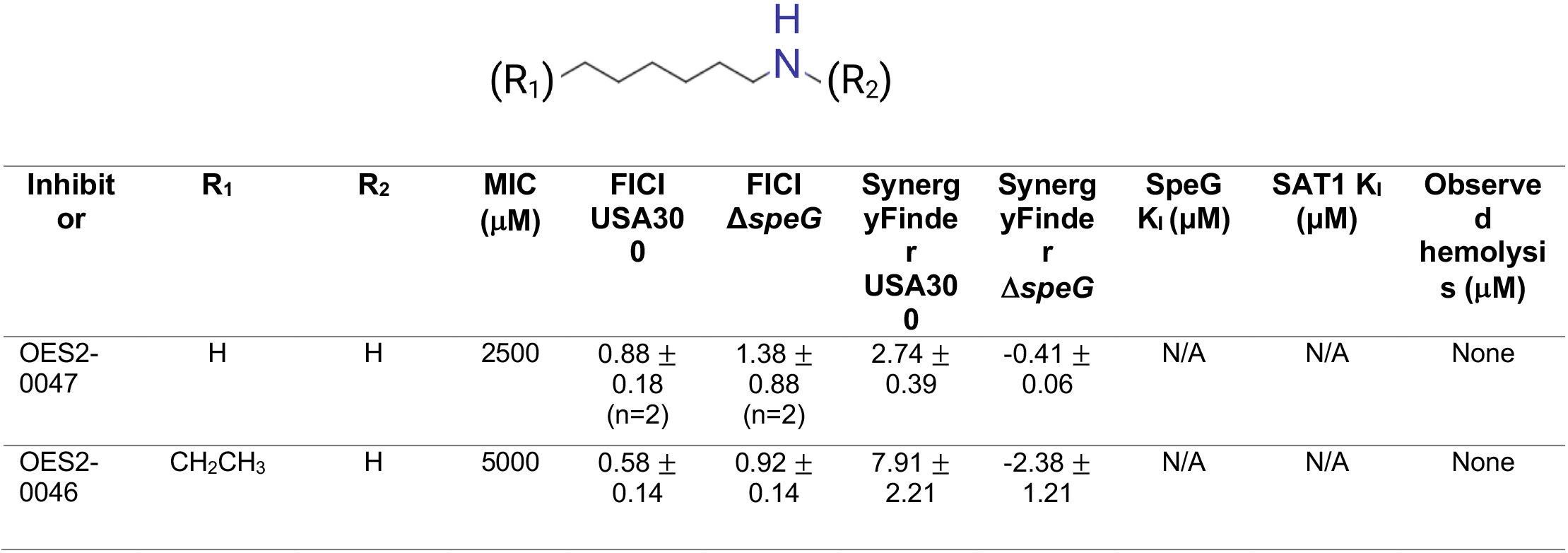
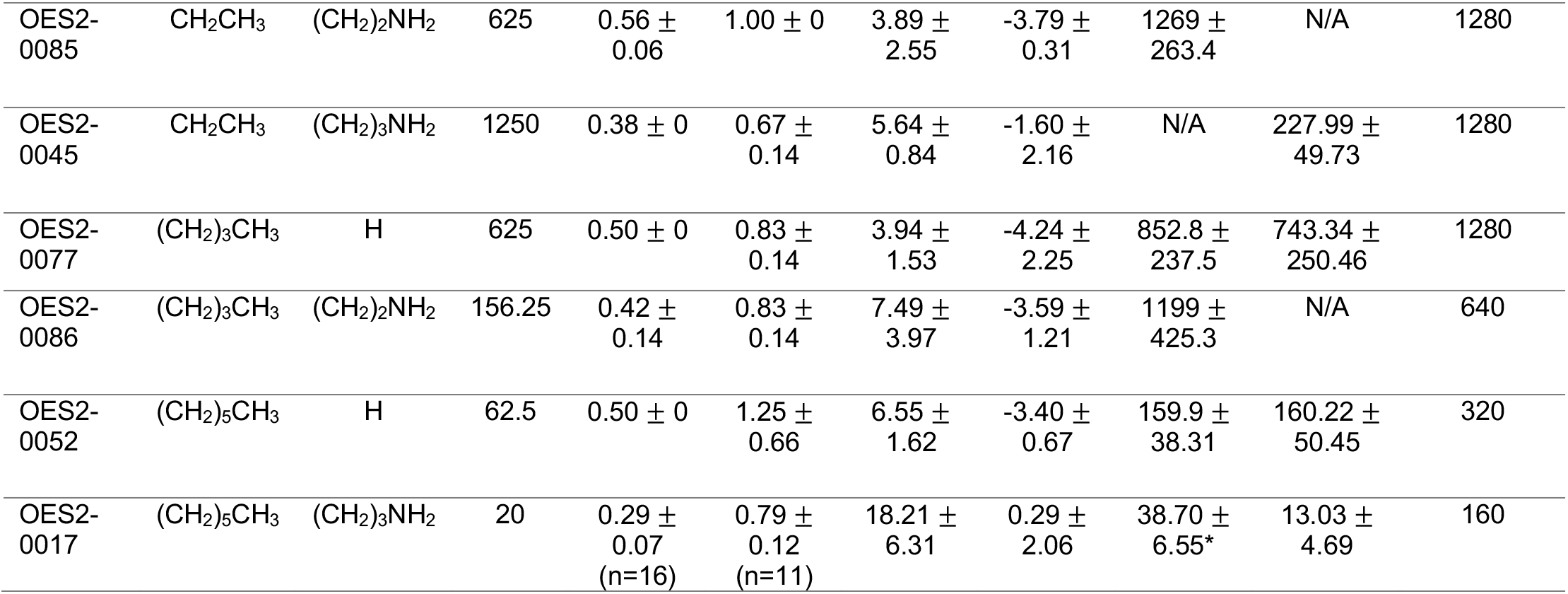
Small scale structure activity relationship study of OES2-0017. SAR study compounds were tested in combination with spermine against *S. aureus* USA300 and Δ*speG*. FICI values are calculated as the mean ± standard deviation of at least three replicates, unless otherwise specified. Inhibitory activity against both SpeG and SAT1 was evaluated in colorimetric enzymatic assays, and Ki values were estimated by the Cheng-Prusoff equation (equation 2) for SAT1 and by equation 3 for SpeG and are reported with relative error (materials and methods). Hemolytic activity of each compound was tested against sheep red blood cells. *Ki value estimated from supplementary Fig. 8.

| Inhibitor | R<sub>1</sub> | R<sub>2</sub> | MIC ($\mu\text{M}$) | FICI USA300 | FICI $\Delta\text{speG}$ | SynergyFinder USA300 | SynergyFinder $\Delta\text{speG}$ | SpeG $K_i$ ($\mu\text{M}$) | SAT1 $K_i$ ($\mu\text{M}$) | Observed hemolysis ($\mu\text{M}$) |
| --- | --- | --- | --- | --- | --- | --- | --- | --- | --- | --- |
| OES2-0047 | H | H | 2500 | $0.88 \pm 0.18$<br>(n=2) | $1.38 \pm 0.88$<br>(n=2) | $2.74 \pm 0.39$ | $-0.41 \pm 0.06$ | N/A | N/A | None |
| OES2-0046 | CH<sub>2</sub>CH<sub>3</sub> | H | 5000 | $0.58 \pm 0.14$ | $0.92 \pm 0.14$ | $7.91 \pm 2.21$ | $-2.38 \pm 1.21$ | N/A | N/A | None |
| OES2-0085 | CH<sub>2</sub>CH<sub>3</sub> | (CH<sub>2</sub>)<sub>2</sub>NH<sub>2</sub> | 625 | 0.56 ± 0.06 | 1.00 ± 0 | 3.89 ± 2.55 | -3.79 ± 0.31 | 1269 ± 263.4 | N/A | 1280 |
| OES2-0045 | CH<sub>2</sub>CH<sub>3</sub> | (CH<sub>2</sub>)<sub>3</sub>NH<sub>2</sub> | 1250 | 0.38 ± 0 | 0.67 ± 0.14 | 5.64 ± 0.84 | -1.60 ± 2.16 | N/A | 227.99 ± 49.73 | 1280 |
| OES2-0077 | (CH<sub>2</sub>)<sub>3</sub>CH<sub>3</sub> | H | 625 | 0.50 ± 0 | 0.83 ± 0.14 | 3.94 ± 1.53 | -4.24 ± 2.25 | 852.8 ± 237.5 | 743.34 ± 250.46 | 1280 |
| OES2-0086 | (CH<sub>2</sub>)<sub>3</sub>CH<sub>3</sub> | (CH<sub>2</sub>)<sub>2</sub>NH<sub>2</sub> | 156.25 | 0.42 ± 0.14 | 0.83 ± 0.14 | 7.49 ± 3.97 | -3.59 ± 1.21 | 1199 ± 425.3 | N/A | 640 |
| OES2-0052 | (CH<sub>2</sub>)<sub>5</sub>CH<sub>3</sub> | H | 62.5 | 0.50 ± 0 | 1.25 ± 0.66 | 6.55 ± 1.62 | -3.40 ± 0.67 | 159.9 ± 38.31 | 160.22 ± 50.45 | 320 |
| OES2-0017 | (CH<sub>2</sub>)<sub>5</sub>CH<sub>3</sub> | (CH<sub>2</sub>)<sub>3</sub>NH<sub>2</sub> | 20 | 0.29 ± 0.07<br>(n=16) | 0.79 ± 0.12<br>(n=11) | 18.21 ± 6.31 | 0.29 ± 2.06 | 38.70 ± 6.55* | 13.03 ± 4.69 | 160 |

The analogs that synergized with spermine lost the synergy against the Δ*speG* mutant (Table 1), suggesting their synergy is mediated through SpeG inhibition similar to OES2-0017. The analogs that did not exhibit synergy in whole-cell assays are hypothesized to have no SpeG inhibitory activity. Therefore, we tested each compound in enzymatic assays, against both SpeG and SAT1 and estimated K_i_ values using equation 2 for SAT1 and equation 3 for SpeG (materials and methods). The specific SSAT inhibition mechanism of each analog, if any, requires further study. For SAT1, we estimated Ki values based on the assumption that the OES2 compound series were acting as competitive inhibitors (discussed below). However, to estimate K_i_ values for the analogs against SpeG, we assumed that OES2-0017 and its analogs would act as non-competitive inhibitors, which may overestimate K_i_ values (see rationale above in the section discussing the OES2-0017-SpeG interaction). OES2-0017 remains the most potent inhibitor of SpeG, K_i_ = 38.70 ± 6.55 μM with OES2-0052 being the second most potent, K_i_ = 159.9 ± 38.31 μM (Table 1 and Supplementary Fig. 32). OES2-0047 and OES2-0046 do not synergize in whole-cell assays and do not inhibit SpeG *in vitro*; thus, they serve as controls to validate the correlation between *in vitro* and whole-cell assays and that our lead compound is targeting and inhibiting SpeG.

OES2-0017 inhibited SAT1 with a K_i_ of 13.03 ± 4.69 μM, lower than that of SpeG, but the K_i_ of OES2-0052 is almost equivalent in SAT1 and SpeG (160.22 ± 50.45 μM and 159.9 ± 38.31 μM, respectively; Table 1 and Supplementary Fig. 32). Similarly, OES2-0077 has almost equivalent K_i_ against SAT1 and SpeG (743.34 ± 250.46 and 852.8 ± 237.5 μM, respectively). Further, two of the tested compounds only inhibited SpeG (OES2-0085 at K_i_ = 1269 ± 263.4 μM and OES2-0086 at K_i_ = 1199 ± 425.3 μM) and not SAT1, in contrast, SAT1 activity increased at the highest tested concentrations of these analogs (Supplementary Fig. 32). Notably, OES2-0045 which synergized with spermine in whole-cell assays (FICI = 0.38 ± 0), suggesting SpeG inhibitory activity, showed no inhibition in the *in vitro* enzymatic assays; however, it inhibited SAT1 (K_i_ = 227.99 ± 49.73μM; Table 1 and Supplementary Fig. 32). These results showing differential activity of some analogs against both enzymes suggest a potential to improve specificity of an inhibitor towards SpeG.

To visualize the interaction of the analogs and SpeG, we performed docking of the analogs (OES2-0045/46/47/52/77/85/86) and spermine into the spermine binding (allosteric) site of the SpeG dimer from PDB 8fv1 (see Methods section; Supplementary Fig 33 and Supplementary table 3). Analogs established the same interaction network as OES2-0017 (Figure 3 G and Supplementary Fig 33). Compounds with the longest aliphatic chains (OES2-0045/52/77/85/86) established more non-polar interactions deep into the allosteric site (Supplementary Fig. 33). We performed MD simulations (see Methods) of the SpeG/ligand complex of OES2-0086 to compare the stability of the docked poses of compounds with an N_1_-N_4_ diammonium group (Supplementary Fig 33 and 34) to that of OES2-0017 and spermine with an N_1_-N_5_ diammonium group (Supplementary Fig 10). Similar to OES2-0017, OES2-0086 moves deeper into the non-polar cavity of monomer B, but in this case, the N_1_-N_4_ polar head establishes transient interactions between amino acids Glu37 (A) and Asp48 (B) carboxylate groups and exhibits high mobility of the aliphatic tail leading to non-stable hydrophobic interactions (Supplementary Fig. 34). The N_1_-N_5_ geometry, the same as spermine, has better coordination with the Glu/Asp pole than the N_1_-N_4_ geometry. The computational results show that N_1_-N_5_ diammonium compounds with long chains are able to bind in a similar way to spermine, and better binding modes are predicted for molecules with a N_1_-N_5_ diammonium group, in agreement with the observed experimental inhibitory activity (Table 1).

Computational studies were also undertaken for SAT1 to study the binding of OES2 series and spermine. In this case, the spermine binding site is located at the intermonomer A-B interface of the SAT1 dimer (PDB ID 2b58), which includes amino acids Glu28 (A), Tyr78 (A), Tyr90 (A), Asp92 (A), Glu93 (A), Asp82 (B), Trp84 (B) and Trp154 (B). Also, the ammonium groups establish ionic interactions reinforced by a network of H-bonds with the carboxylate sidechains of Asp82 (B), Asp92 (A) and Glu93 (A), and further stabilized by cation-π interactions between the aliphatic moieties and Trp84 (B) side chain, while the N_1_ ammonium group interacts with the catalytic residues Glu28 (A) and Trp154 (B) (Figure 9 and Supplementary Fig. 35). OES2 compounds were predicted to bind two sites in SAT1 that differ from the spermine binding site (Supplementary Fig. 36). Binding site 1 includes the acid residues Asp82 (B), Asp92 (A) and Glu93 (A) along with the Trp84 (B), as well as a non-polar cavity which is nearby, mainly formed by amino acids Tyr78 (A), Tyr90 (A), Leu128 (A), Thr80 (B), Leu88 (B), Tyr90 (B), Trp154 (B) and Leu156 (B). In this site, N_1_, N_1_-N_4_ or N_1_-N_5_ ammonium groups of all compounds established ionic/H-bonds networks with the carboxylate groups from Asp82 (B), Asp92 (A) and Glu93(A) reinforced with cation-π interactions of the one ammonium with Trp84 (B) side chain (Figure 9 and Supplementary Fig. 35). The aliphatic chains bind into the above-described non-polar cavity (Figure 9 and Supplementary Fig. 35). Compounds with the longest aliphatic chain established more non-polar and CH-π interactions, yielding the best docking results (Figure 9, Supplementary Fig. 35, and Supplementary Table 3).

**Figure 9.**
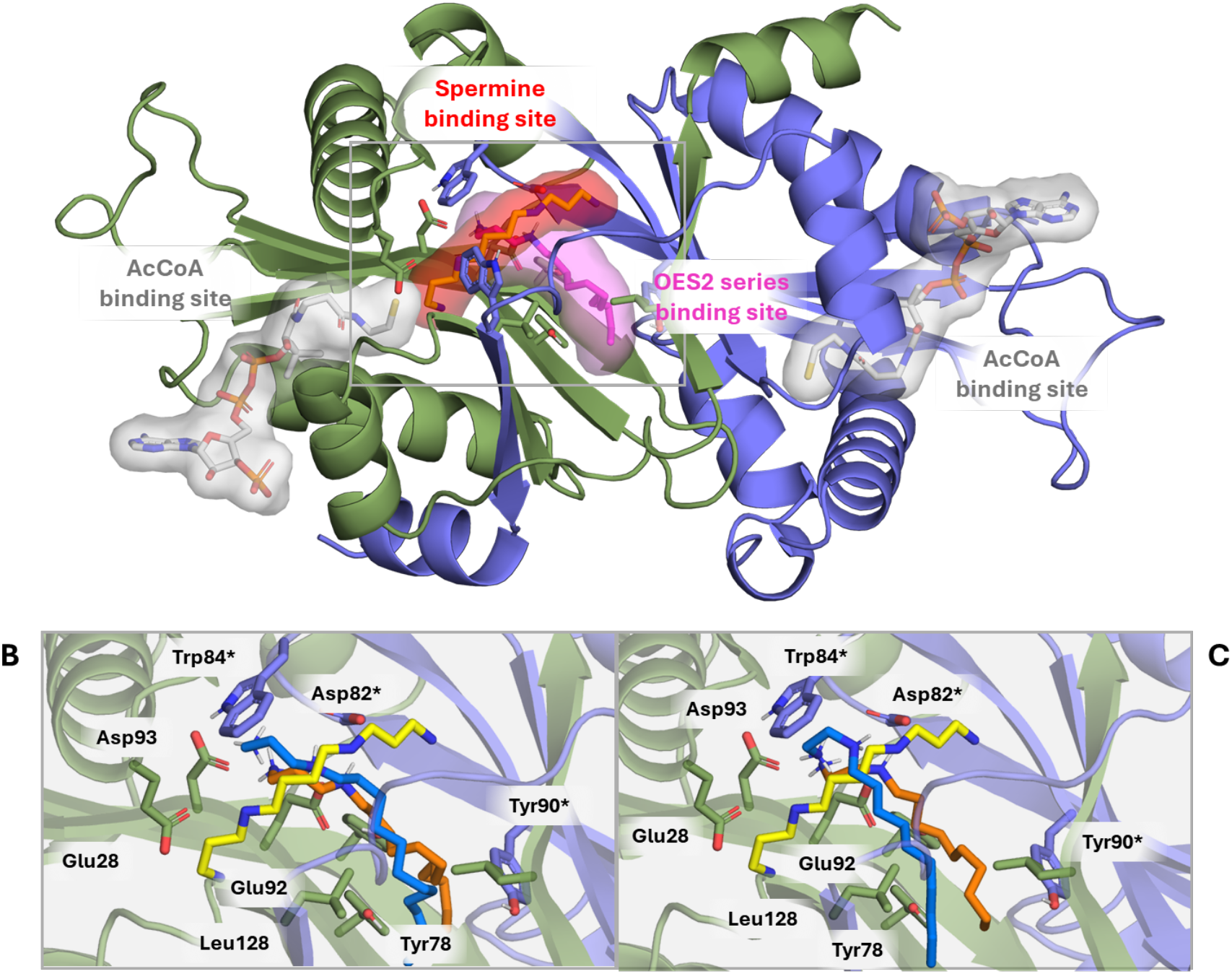
Insight into the binding mode of studied compounds to human SAT1 dimer (PDB ID 2b58). **A)** General view of the spermine binding site (red surface), the OES2 series binding site 1 (pale pink surface) and the AcCoA binding site (white surface). **B-C)** Detail of the interactions of the crystallographic bound pose of spermine (yellow, obtained by superimposition of PDB 3bj8 with PDB 2b58), the docked conformation (orange) and the conformation after 300 ns of MD simulation (blue) of compounds OES2-0017 (B) and OES2-0086 (C). Monomer A in green and monomer B in blue.

MD simulations were also undertaken to study the stability of the complexes of SAT1 with OES2-0017 and OES2-0086. For site 1, both complexes remained stable along the simulations and predicted ligand-protein interactions were stabilized: N_1_-N_5_ and N_1_-N_4_ diammonium moieties established ionic and H-bonding networks with Asp82 (B), Asp92 (A) and Glu93 (A) carboxylate groups, also reinforced by cation-π interactions with Trp84 (B) (Figure 9 and Supplementary Fig. 37). MM-GBSA energy calculations show that the N_1_-N_4_ diammonium moiety of OES2-0086 exerted a less favourable binding mode when establishing this network than the N_1_-N_5_ moiety of OES2-0017 (Supplementary Table 5). For site 2, the SAT1/ligand complexes also showed a stable binding mode (Supplementary Fig. 38 and Supplementary Table 5), suggesting it is possible for the polyamine analogs to bind this site. However, given the fact that site 2 is the AcCoA binding site and that all OES2 compounds were predicted to bind with weaker binding energies than in site 1, site 2 seems a less plausible binding site (Supplementary Fig. 36).

Together, the observed experimental structure-activity relationships (Table 1) were in agreement with the predicted *in silico* computational results, which suggest that compounds with longer aliphatic chains exerted the best performance, as well as the N_1_-N_5_ disubstituted pattern (spermine and OES2-0017), which exerted more efficient interactions with both, SpeG and SAT1 than the N_1_-N_4_ (OES2-0085 and OES2-0086) or the monoaminated derivatives (OES2-0052 and OES2-0077). SAT1 is unable to acetylate any of the OES2 analogs likely given that the predicted analog binding sites are different from that of spermine and that the compounds are not predicted to interact with the necessary catalytic residues (Figure 9 and Supplementary Fig. 35 and 36).

Given that the compounds also exhibit growth-inhibitory effects, we simultaneously evaluated their hemolytic activity. Hemolytic activity aligned with the MIC data trends, where the concentration required to lyse red blood cells was observed at lower concentrations for compounds with a longer alkyl chain (R_1_ = (CH_2_)_3_CH_3_ or (CH_2_)_5_CH_3_) and with an aminoethyl or aminopropyl group at R_2_ (Table 1 and Supplementary Fig. 39). Hemolysis for the SpeG inhibitory compounds OES2-0085 and OES2-0077 occurred at 1280 μM (∼2-fold higher than their MICs), while hemolytic activity of OES2-0086, and OES2-0052 was observed at ∼ 4-fold higher than their MICs (640 and 320 μM, respectively; Table 1 and Supplementary Fig. 39). We observed no hemolytic activity of two analogs with high MIC values OES2-0047 and OES2-0046 (Table 1 and Supplementary Fig. 39). At concentrations equivalent to their respective MIC values against *S. aureus* USA300, we observed no significant HepG2 cell damage of OES2-0047, OES2-0046, OES2-0052, and OES2-0017 compared to a DMSO-treated control (Supplementary Fig. 40). Given the reduced activity of OES2-0052 against SAT1 compared to OES2-0017 and its lack of observable toxicity against eukaryotic cells, we tested if this compound retains the potential adjuvant activity of OES2-0017. OES2-0052 abolishes the protective effect of spermine towards vancomycin, kanamycin, and rifampicin, and potentiates the activity of cefuroxime (Supplementary Fig. 41). This analysis provides a foundation for a larger scale targeted lead optimization to improve the potency of SpeG inhibition, while reducing the hemolytic and SAT1 inhibitory activity.

## Discussion

The cellular concentrations of polyamines in mammalian cells vary across tissues and physiological conditions ^18^. Polyamines are known to bind to macromolecules, such as DNA, RNA, phospholipids, and ATP, so levels of free intracellular polyamines are not definitive ^18^. In polyamine-producing bacteria, the contents of intracellular polyamines have been reported as high as 30 mM ^99^ and polyamine concentrations in the host increase ∼2-fold in response to infection ^13^. Detrimental effects of *speG* or alternative SSAT mutations on bacterial virulence ^13,28,29^ suggest that polyamine detoxification is a promising target for drug development and prompted us to identify inhibitors of bacterial polyamine detoxification to serve as antivirulence agents. Antivirulence agents are explored as emerging new antimicrobial solutions; however, the current repertoire of antivirulence strategies in preclinical and clinical development mostly focuses on disrupting “traditional” bona-fine virulence factors, including quorum sensing, secretion systems, bacterial toxins, stress response regulators, biofilm formation, adhesion, and conjugation ^100–106^. In contrast, our work seeks to disrupt virulence driven by the bacterial ability to respond to specific host-produced metabolites and other chemicals encountered at the infection site. Such chemicals, including polyamines, have been shown to influence bacterial colonization, proliferation and virulence thus making this approach an attractive yet underexplored antivirulence strategy ^10^. Herein, we carried out a high-throughput chemical screen and uncovered two lead compounds, OES2-0017 and OES1-1087, that show growth inhibition and synergy with spermine. These compounds exhibit broad-spectrum activity against Gram-positive and Gram-negative bacteria, including *K. pneumoniae, S.* Typhimurium and *S. epidermidis*, and the fungus *C. albicans.* We show that OES2-0017 inhibits the spermine/spermidine acetyltransferase SpeG and exhibits growth inhibition via membrane perturbation. OES1-1087 reduced *S. aureus* intracellular AcCoA concentration necessary for SpeG-mediated spermine detoxification and increased membrane fluidity, likely exacerbating membrane damage by spermine. OES2-0017 and OES1-1087 disruption of SpeG activity likely reduces the ability of bacteria to withstand host polyamines, conceivably underlying our observations that OES2-0017 treatment prevented intracellular survival of *S.* Typhimurium in murine macrophages and OES1-1087 reduced bacterial burden in a murine *S.* Typhimurium gastrointestinal infection.

Polyamine analogs have previously been demonstrated to have antiparasitic, anticancer ^46,98,107^, and antibacterial properties ^108–110^. Antibacterial polyamines described to date include mono-and bis-acyl polyamines, amine steroidal derivatives ^108^, and diacylpolyamines with aromatic head groups ^109^, which are structurally different from OES2-0017 and the other polyamine analogs tested in this study. One study tested linear polyamine analogs, similar to OES2-0017, and revealed bactericidal and β-lactam potentiating activity; however, the mechanism of growth inhibition and antibiotic potentiation were not discussed ^110^. In our study, the polyamine analog OES2-0017 similarly synergized with the tested β-lactam, cefuroxime, potentiated azithromycin activity, and abolished the antagonism between spermine and vancomycin, rifampicin, and kanamycin against *S. aureus*. Thus, we suggest the compound identified in our study has antibiotic adjuvant properties.

Specific bacterial SSAT inhibitors have not been identified previously. Notably, berenil (diminazene aceturate) was shown to inhibit SSAT and has been used *in vivo* in murine models for this purpose ^94^. *In vitro* assays against human SSAT suggested berenil is a potent competitive inhibitor of spermidine with a K_i_ of 2.0 μM ^111^; however, berenil also inhibits rat S-adenosyl-L-methionine decarboxylase and mouse polyamine oxidase ^111,112^. Activity against multiple enzymes may explain the toxic effects observed during intramuscular injection of camels with 10 mg/kg berenil ^113^; however no toxic effects were observed in mice dosed at 16 mg/kg weekly for six weeks ^94^ or with a single intraperitoneal or intramuscular dose of 3.5, 10, or 20 mg/kg ^113,114^. Aside from the SSAT activity, berenil was shown to have other activities, including immunomodulatory ^115^ and cardioprotective ^116^ activities, and is used in veterinary medicine mostly for parasitic infections albeit the exact underlying mode of such activity is not fully elucidated ^117^. Our study provides insights that may lead to the development of specific bacterial SSAT inhibitors. Our data suggest a concentration range within which OES2-0017 may remain efficacious against bacteria while avoiding potential adverse effects as it did not exhibit cytotoxicity towards HepG2 cells or hemolytic activity at concentrations equivalent to 1x and 2x the MIC against *S. aureus* USA300, respectively, while its SpeG-inhibitory activity was observed as low as 1/32^nd^ the MIC. Future studies will assess the potential toxicity of OES2-0017 and its analogs and their interaction with other polyamine-related enzymes that use natural polyamines as substrates.

*In vitro* enzymatic assays show that OES2-0017 is active against both SpeG and SAT1, suggesting potential activity against the human enzyme *in vivo*. SAT1 inhibitors have other potential therapeutic uses including disrupting tumor growth, protection from kidney injury following ischemia-reperfusion injury, and preventing osteoporosis ^90,92,94,98^. A small-scale SAR study identified compounds with differential activity towards the human SAT1 and *S. aureus* SpeG. Two analogs of OES2-0017, OES2-0085 and OES2-0086, showed inhibitory activity against SpeG but not SAT1. While the analogs OES2-0052 and OES2-0077, are almost equally potent inhibitors of SpeG and SAT1, suggesting improved specificity relative to OES2-0017. Although these analogs have lower inhibitory activity than OES2-0017, they and other analogs from our analysis can guide future SAR studies to optimize the inhibition of SpeG while limiting activity against SAT1.

We also identified OES1-1087 (isoproterenol) as a spermine synergist. Isoproterenol is a β1-β2-adrenergic agonist used in the treatment of heart block, heart failure, and cardiac arrest ^118^. Its identification in our chemical screen serves as an example of potential drug repurposing for the treatment of bacterial infections. Time and money spent on drug development are drastically reduced by repurposing as pharmacodynamic, pharmacokinetic, and toxicity profiles of the drugs are already known ^43^. Our results indicate that OES1-1087 reduces intracellular AcCoA content, likely impeding SpeG-mediated acetylation of spermine, and increases membrane fluidity, potentially enhancing the polyamine-mediated membrane perturbation. Enzymatic data and *in silico* docking studies suggest OES1-1087 may also interact with SpeG at the AcCoA binding site, although this interaction is relatively weak. OES1-1087 significantly reduced *S.* Typhimurium bacterial burden in the liver and spleen of a murine gastrointestinal infection model confirming *in vivo* antimicrobial activity. Importantly, it did not show detectable activity against the human SAT1 and no hemolytic activity or HepG2 cytotoxicity.

*S. aureus* was not previously known to have an SSAT except for strains that acquired SpeG, such as USA300. Notably, the USA300 Δ*speG* and other *S. aureus* strains that do not harbor *speG* (e.g., COL and NCTC8325) are only 4-fold and 8-fold more susceptible to spermidine and spermine, respectively, compared to wild-type USA300, suggesting a potential for another detoxification mechanism. Here, we identified PaiA_Sa_, a homolog of *B. subtilis* spermine/spermidine acetyltransferase PaiA, in *S. aureus,* providing its first experimental characterization as an additional SSAT in *S. aureus*. SpeG appears to be more dominant with a greater shift in polyamine susceptibility observed in a Δ*speG* mutant than *paiA*_Sa_::Tn; the contribution of PaiA_Sa_ to polyamine resistance in whole cells requires assessment in a *speG paiA_Sa_* double mutant in USA300 or a single *paiA_Sa_* mutant in a *speG^-^S. aureus* strain. We further showed that OES2-0017 can inhibit PaiA_Sa_ *in vitro*.

This work provides insights into understudied aspects of chemically mediated host-pathogen interactions. We discovered two novel inhibitors of spermine/spermidine *N*-acetyltransferase. The discovered inhibitors have potential applications as stand-alone antimicrobials, antivirulence agents, or antibiotic adjuvants. We provide *in vivo* and *in cellulo* proof of concept for these antimicrobials, which may further expand antivirulence strategies to include disrupting bacterial responses to chemicals at infection sites and chemically mediated host-pathogen interactions. These antimicrobials may serve as leads for further development, whereas isoproterenol (OES1-1087) could serve as a candidate for drug repurposing. Future work will aim to optimize target specificity and determine pharmacokinetic and pharmacodynamic properties of the active molecules, as well as to test the safety and efficacy of polyamine detoxification inhibitor-antibiotic combinations *in vivo*. Together, this work provides antimicrobial compounds with a novel mechanism to fight multidrug-resistant priority pathogens.

## Methods

### Bacterial strains and reagents

Supplementary Tables 6 and 7 list strains, plasmids and primers used in this work. Bacteria were grown in cation-adjusted Mueller Hinton medium (MHB) at 37°C, except for *Enterococcus faecalis* and *Enterococcus faecium*, which were grown in Tryptic Soy Broth and *Mycobacterium smegmatis,* which was grown in Luria Broth. **Antimicrobial susceptibility testing.** Minimum inhibitory concentration (MIC) and checkerboard assays were performed using the CLSI broth microdilution technique ^8^. The fractional inhibitory concentration indices (FICI) were calculated as FICI = (MIC_drug A_ in combination/MIC_drug A_ alone) + (MIC_drug B_ in combination)/MIC_Drug B_ alone). FICI values were interpreted as synergy when FICI ≤ 0.5, no interaction when 1 ≤ FICI ≤ 4, and antagonism when FICI > 4.0. Drug combination responses were also analyzed based on ZIP reference model with partial correction for negative values using SynergyFinder ^48,49^. Using this model, synergy scores between two compounds of less than -10 are likely to be antagonistic, between -10 and 10 are likely to be additive, and larger than 10 are likely to be synergistic.

### Chemical screen and polyamine analog library assembly

We assembled a library of polyamine analogs by selecting commercially available (Aldrich Market select) analogs with at least 60% structural similarity to substrates and products of polyamine biosynthesis and detoxification enzymes in bacteria (Supplementary tables 1 and 2). This library was screened at 20 μM in the presence and absence of spermine at 1/4th the wild-type MIC (5 mM). Library compounds (1 μL) were added to 96-well plates (Nunc) and filled to 100 μL of inoculated MHB. We also screened a library of previously approved drugs and natural product derivatives (Spectrum collection, MicroSource Inc.) at 20 μM against the wild-type USA300 strain in the presence of spermine at 1/4th the wild-type MIC (2.5 mM). Compounds (0.1 μL each) were administered to 384-well plates (Nunc) containing 50 μL of inoculated MHB with spermine using the Biomatrix BM6-BC (S&P Robotics inc.). For both screens, plates were incubated at 37°C, and bacterial growth was determined by reading OD_600_ after 21 and 16 hours, respectively, based on Z’ values calculated prior to screening using equation 1 ^119^. Hits from the screen were determined by a cut-off of at least 80% growth inhibition relative to a growth control and compounds listed as antibiotics in MicroSource Inc. source files were excluded (Supplementary Table 8).

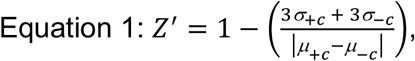

where σ is the standard deviation and μ the mean of the positive (+c) and negative controls (-c).

### Chemical screen and polyamine analog library dose-response assays

Compounds identified as primary hits were added to 96-well plates, filled with inoculated MHB with or without 1/4 MIC spermine (2.5 mM), and serially diluted in the inoculated media to produce dose-response curves. Each compound was tested in duplicate in the presence and absence of spermine. IC_50_ values were determined by fitting a nonlinear dose-response inhibition model in GraphPad Prism 10. Compounds displaying ≥ 2-fold decrease in IC_50_ in the presence of spermine or an IC_50_ in the spermine condition and no IC_50_ in absence of spermine in at least one replicate and that were commercially available and had no reported antimicrobial activity in literature were selected for further follow up.

### Chemogenomic Screening

#### *S. aureus* Nebraska Transposon Mutant Library (NTML) screen

Overnight cultures of the NTML ^50^ were prepared in 384-well plates using Biomatrix BM6-BC (S&P Robotics inc.) in 50 μL MHB containing 5 μg/mL erythromycin. The following day, MHB containing the drug screening condition of 1/4^th^ (5 mM), 1/8^th^ (2.5 mM), or 1/16^th^ (1.25 mM) the wild-type USA300 MIC of spermine, 1/4^th^ (312.5 μM) or 1/2 (625 μM), the wild-type USA300 MIC of OES1-1087 were inoculated using the Biomatrix BM6-BC. Plates were grown at 37°C, and OD_600_ was read after 24 h.

#### *B. subtilis* CRISPRi essential gene knockdown library screen

An overnight culture of the library ^120^, was prepared in a 384-well plate using Biomatrix BM6-BC (S&P Robotics inc.) in 50 μL MHB containing 6 μg/mL chloramphenicol. The following day, MHB containing OES2-0017 at the wild-type MIC and at 2-fold dilutions to 1/16^th^ the MIC (2.5-40 μM) and in the absence of OES2-0017 were inoculated using the Biomatrix BM6-BC. The experiment was performed with 0.05% xylose (allowing low level of *dcas9* expression) and without xylose (allowing basal *dcas9* expression). Plates were incubated at 37°C, and OD_600_ was read after 24 h.

#### General molecular techniques

Unmarked in-frame deletion of *speG* was performed using the temperature-sensitive allelic exchange plasmid pJB38 ^121^. Approximately 1-kb upstream and downstream of *speG* were PCR amplified and the upstream fragment digested with EcoRI and BamHI (New England Biolabs), and the downstream fragment digested with EcoRI and SalI (New England Biolabs). pJB38 was digested with BamHI and SalI and fragments were ligated with T4 DNA ligase (New England Biolabs). The resultant deletion plasmid pJB38-Δ*speG* was passaged through *E. coli* DC10B and selected for on 100 μg/mL ampicillin plates before electroporation into *S. aureus* USA300 JE2 and selected for on 10 μg/mL chloramphenicol plates ^122^. Deletion of *speG* was confirmed by PCR. The construction of *speG, pdhA,* and *lpdA* complementation vectors utilized pKK22 with a *S. aureus fba* promoter. *speG, pdhA* and *lpdA* were PCR amplified from *S. aureus* USA300 and the PCR products and plasmid were digested with AvrII and SacI (New England Biolabs). Digested fragments were ligated with T4 DNA ligase (New England Biolabs) and transformed into *E. coli* DH5α λ pir before passage into *S. aureus* RN4220 and finally into *S. aureus* USA300 Δ*speG, pdhA*::Tn, or *lpdA*::Tn, respectively. All transformation steps were selected for on 30 μg/mL trimethoprim.

#### Protein overexpression and purification

*speG* (encoding UniProt ID: A0A0H2XGJ0), SAUSA300_2083 (encoding UniProt ID: A0A0H2XDE9), SAUSA300_0441 (encoding UniProt ID: A0A0H2XIC5), and *paiA* (encoding UniProt ID: A0A0H2XGV3) were cloned from *S. aureus* USA300 into pET28a(+) and passaged through *E. coli* DH5α before transformation into *E. coli* BL21, all transformation steps were selected for on 75 µg/mL kanamycin. cDNA of *SAT1* (encoding UniProt ID: P21673) was codon-optimized, synthesized from, and cloned into pET28a(+) (Bio Basic Canada). Proteins were overexpressed under conditions previously described ^23^. Lysis was achieved using a Branson Sonifier 450 and the supernatant was isolated from the insoluble fraction by centrifugation at 10 000 xg for 30 minutes at 4°C. His-tag batch purification was performed using His-select nickel-affinity gel (Sigma-Aldrich). The gel was equilibrated with 50 mM sodium phosphate, pH 8.0 with 0.3 M sodium chloride before the addition of the cell lysate. Cell lysate was mixed with the His-select nickel-affinity gel overnight at 4°C and washed 3X with wash buffer (50 mM sodium phosphate, pH 8.0 with 0.3 M sodium chloride and 10 mM imidazole). Proteins were eluted with elution buffer (50 mM sodium phosphate, pH 8.0 with 0.3 M sodium chloride and 250 mM imidazole). Purified proteins were detected on a 12% acrylamide gel (Bio-Rad) by PageBlue staining solution (Fisher Scientific). Proteins were dialyzed in 100 mM Tris-HCl pH 7.5, 1 mM EDTA for colorimetric enzymatic assays or in 20 mM HEPES, pH 8.0 for HPLC using 10 kDa SnakeSkin^TM^ (Fisher Scientific) dialysis tubing then quantified using the Bradford method with protein assay dye reagent concentrate (Bio-Rad). Glycerol was added to a final concentration of 10% and aliquots of proteins were stored at -80°C. Protein aliquots were thawed on ice for use, with each aliquot thawed only once for immediate use. A relative protein purity is shown in Supplementary Fig. 42. The His tag was not cleaved for enzymatic assays as we performed a test with SpeG and observed no difference in the presence or absence of the tag on enzymatic activity (Supplementary Fig. 42).

### *In vitro* SSAT enzymatic assays

#### Steady state kinetics

Purified enzymes were used in spermine/spermidine *N*-acetyltransferase (SSAT) colorimetric enzymatic assays as described previously ^23,54^, with modifications. A 4X enzyme solution, 4X substrate solution and 2 mM 5, 5’-dithio-bis-(2-nitrobenzoic acid) (DTNB/Ellman’s reagent; Sigma Aldrich), were each prepared in 100 mM Tris-HCl, pH 7.5, 1 mM EDTA and added to a 96-well plate in the following order 25 μL enzyme solution, 50 μL Ellman’s reagent, and the reaction was initiated by adding 25 μL of substrate solution. Each reaction plate contained serial dilutions of Coenzyme A (5-150 nmol) as a standard. Upon initiating the reaction, plates were immediately transferred to a pre-warmed plate reader (Biotek), and reactions were allowed to proceed at 37°C with A_412 nm_ readings taken each minute for 45 min. The final enzyme concentration was 0.8 μg for SpeG, SAUSA300_2083, SAUSA300_0441, and SAT1 and 3.2 μg for PaiA_Sa_. The substrate solution contained a final concentration of 1 mM AcCoA (Sigma Aldrich) for SpeG, PaiA_Sa_, SAUSA300_2083, and SAUSA300_0441, and 0.25 mM AcCoA for SAT1 and variable concentrations of polyamines. SpeG was tested with spermine concentrations ranging from 0-4 mM, spermidine from 0-10 mM, and putrescine from 0-8 mM. PaiA_Sa_ was tested with spermine concentrations ranging from 0-8 mM, and spermidine and putrescine from 0-10 mM. SAUSA300_2083, and SAUSA300_0441 were tested with spermine concentrations ranging from 0-6 mM and spermidine and putrescine concentrations ranging from 0-8 mM. SAT1 was tested with spermine concentrations ranging from 0-1.5 mM. Each reaction condition is background subtracted by a corresponding heat inactivated control treated with the same concentration of polyamine. Kinetic parameters were estimated by fitting with the Michaelis-Menten equation and/or allosteric sigmoidal equation in Graphpad Prism 11.

#### Enzymatic inhibition assays

The colorimetric SSAT assay was performed where 25 μL of enzyme solution containing 0.8 μg enzyme, 100 mM Tris-HCl, pH 7.5, 1 mM EDTA and a dilution series of inhibitor was added to a 96-well plate. To account for any solvent effects, an 8X dilution series of each inhibitor was prepared in DMSO and 12.5 μL added to the final reaction volume of 100 μL, so that each condition received the same final concentration of DMSO. The enzyme-inhibitor solution was incubated at room temperature for 30 min to allow equilibration in the event inhibitors were slow-binding ^57^ before adding 50 μL Ellman’s reagent and 25 μL substrate solution to initiate the reaction. Some reactions were initiated without pre-incubation of the enzyme-inhibitor solution. OES2-0017 and analogs in the small-scale structure-activity relationship study were assayed with a substrate solution including a fixed concentration of spermine ≈ K_m_ (1500 μM for SpeG and 75 μM for SAT1) and 1 or 0.25 mM acetyl-CoA with SpeG and SAT1, respectively. OES1-1087 inhibition was assayed with a substrate solution including 1500 μM spermine and an AcCoA concentration at ≈ K_m_ (0.75 mM). Substrate solution to evaluate OES2-0017 inhibition of PaiA_Sa_ contained a final concentration of 1 mM AcCoA and spermine at ≈ K_m_ (2 mM). The reaction was monitored each minute at A_412 nm_ for 45 minutes. Absorbance of the reactions containing inhibitor were compared to a positive control containing only spermine, AcCoA, and solvent (DMSO). Data for IC_50_ plots were fit to a nonlinear regression inhibition model in GraphPad Prism 11 and K_i_ was estimated from equation 2 ^123^ for SAT1 and equation 3 for SpeG. The colorimetric SSAT assay was also performed in a 96-well plate where enzyme activity was monitored in the presence of the respective inhibitor across 8 concentrations of substrate and data were fit to a noncompetitive (SpeG) or competitive (PaiA_Sa_ and SAT1) inhibition model in Graphpad Prism 11.

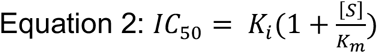

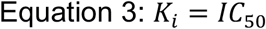

#### HPLC quantification of primary amines

Quantification of primary amines was achieved following pre-column derivatization with *o*-phthalaldehyde (OPA) and *N*-acetyl-L-cysteine (NAC) following the protocol described in ^56^, with some modifications. Enzymatic reactions were performed by mixing 50 μL of a 2X enzyme solution and 2X substrate solution together and incubating the reaction at 37 °C for the indicated amount of time. All reactions were performed in 20 mM HEPES pH, 8.0 for HPLC detection and concentrations reported below are the final concentration in a 100 μL reaction. All assays with SpeG and SAT1 contained 0.8 μg of enzyme, while PaiA_Sa_, SAUSA300_2083, and SAUSA300_0441 reactions contained 3.2 μg of enzyme. For assaying spermine acetylation, the substrate solution consisted of a final concentration of 1 mM AcCoA and 1 mM spermine for all tested enzymes. SpeG and SAT1 reactions were carried out at 37°C for 10 minutes, while PaiA_Sa_, SAUSA300_2083, and SAUSA300_0441 reactions were carried out at 37°C for 1 hour. To assess OES2-0017 inhibition of SpeG, the substrate solution contained a final concentration of 1 mM spermine, 1 mM AcCoA, and 300 μM OES2-0017. To assess OES2-0017 inhibition of SAT1, the substrate solution contained a final concentration of 75 µM spermine, 1 mM AcCoA, and 300 μM OES2-0017. Reactions were incubated at 37°C for 10 minutes. OES1-1087 inhibition of SpeG was tested with an enzyme solution consisting of 0.8 μg of SpeG and 3 mM OES1-1087 that was incubated for 30 min at room temperature before initiating the reaction with 1 mM spermine and 0.75 mM Acetyl-CoA and allowing the reaction to proceed at 37°C for 5 minutes. For assessing SAT1 inhibition by OES1-1087, the substrate solution contained a final concentration of 1 mM spermine, 0.75 mM AcCoA and 3 mM OES1-1087. The SAT1 reaction was incubated at 37°C for 10 minutes. For assessing OES2-0017 inhibition of PaiA_Sa_, the substrate solution contained a final concentration of 1 mM spermine, 1 mM AcCoA and 1.2 mM OES2-0017. The PaiA_Sa_ inhibition reaction was incubated at 37°C for 1 hour. All inhibition assays contained controls with the same concentration of DMSO used in the treatment condition and were performed with a heat inactivated enzyme control receiving both DMSO and inhibitor treatment. After incubation at 37°C, the reaction was stopped by the addition of 100 μL of ice-cold 1.5 M perchloric acid and vortexed at room temperature for 1 minute. 50 μL of ice-cold 2 M potassium carbonate was added and CO_2_ allowed to release before being vortexed for 1 minute at room temperature followed by centrifugation at 15 000 *x*g at 4°C for 10 minutes. The supernatant was extracted and diluted to a final dilution factor of 20 or 25 into HPLC-grade water containing a final concentration of 0.075% benzoic acid. Samples were derivatized with OPA/NAC, which was prepared fresh, using in-line pre-column derivatization as described in ^56^. Samples were separated on an Agilent 1200 series HPLC system equipped with an Agilent TC-C18 guard column (4.6 x 12.5 mm, 5 μm, Cat. # 520518-905) and Agilent TC-C18 column (4.6 x 150 mm, 5 μm Cat. # 588935-902). The HPLC was run in gradient mode (Table 2) with solvent A (0.1 M sodium acetate (Fisher); pH 7.2, 9% methanol, 0.5% tetrahydrofuran (Fisher)) and solvent B (100% HPLC-grade methanol) ^56^. The total run time for each sample including column regeneration was 30 minutes. Fluorescence of each polyamine derivative was detected with an excitation λ of 340 nm and emission λ of 450 nm. ACD labs software was used to analyze each chromatogram and peaks were quantified by running an external standard of each *N*^1^-acetylspermine (≥ 97%), spermine (≥ 96%), and OES2-0017 (98%; Ambeed) to construct a standard curve. All chemicals were purchased from Sigma unless otherwise specified.

**Table 2.** HPLC gradient program used for the separation of derivatized amines.

| Time (min) | Solvent A (%) | Solvent B (%) |
| --- | --- | --- |
| 0 | 70 | 30 |
| 9 | 35 | 65 |
| 15 | 0 | 100 |
| 20 | 0 | 100 |
| 27 | 70 | 30 |
| 30 | 70 | 30 |

### Computational studies

#### Protein preparation

Crystallographic structures of *Staphylococcus aureus* SpeG in absence or presence of spermine (PDB IDs 5ix3 and 8fv1, respectively) and the apo form of the human SSAT (SAT1; PDB ID 2b58) were obtained from the Protein Data Bank ^124^. These structures were selected as the best initial structure for each question addressed in this paper. The protein structures were submitted to a structural optimization with Maestro Package ^125^, which consist of removal of water molecules; assessment of the histidine protonation state at pH 7.4 with PROPKA module of Maestro package; addition of missing hydrogen atoms, residues or side chains; the addition of cap residues such as an acetyl group in the N’ terminal of each monomer and a N-methylamine group in the C’ terminal ends. Finally, a minimization in vacuum using OPLS4 force field to optimize the geometry of the side chains was performed ^126^.

#### Ligand preparation

Selected ligands were modelled in Chimera 1.18,^127^ the AM1-BCC charges were computed with ff14SB (Antechamber), ^128^ and a minimization consisting of 5000 steepest descent steps followed by 5000 conjugate gradient steps was performed.

#### Molecular docking calculations

Docking calculations with AutoDock4.2 ^129^ were submitted for three different binding sites in the prepared SpeG holo receptor (8fv1), and for two different binding sites of SAT1 (2b58) for all polyamine compounds (OES2-0017/45/46/47/52/77/85/86; Supplementary Table 3). For the docking calculations with SpeG (8fv1) in the allosteric site, a grid box was centred at the carbonyl oxygen of Tyr32 (monomer A) with a size of 60, 60, 60 and a grid spacing of 0.375 Å. To validate the docking method, spermine was docked following the same protocol finding that the top ranked predicted pose was the same as the crystallographic pose. For docking calculations in the acceptor site of SpeG in the presence of the crystallographic structure of spermine (PDB 8fv1) in the allosteric site, we employed a grid box centred between Glu79 (monomer A) and Phe28 (monomer A) with a size of 60, 60, 60, and a grid spacing of 0.375 Å. For docking calculations in the AcCoA binding site, we employed a grid box centred between Trp27, Ile84 and Tyr129 of monomer A, with a side box of 60, 60, 60 and a grid spacing of 0.375 Å. For SAT1 (2b58), the grid box was centred at the indole nitrogen of the side chain of Trp154 (monomer B) with a box size of 80, 80, 80 and grid spacing of 0.375 Å. For all studied systems, genetic algorithm parameters were used, and 30 different solutions were retrieved for each ligand. AutoDockTools ^129^ and Pymol ^130^ software were employed for the visual inspection of the obtained poses.

#### Molecular dynamics simulations

The protein/ligand complexes from the best predicted docked poses of spermine and compounds OES2-0017 and OES2-0086 were submitted to all-atom MD simulations, performed with Amber16 ^131^. Proteins were described using the force field ff14SB ^132^, with an octahedral simulation box whose edges are distant by at least 10 Å of any atoms. The TIP3P water molecules model was employed, and the charges of the system were neutralized with Na**^+^** and Cl**^-^**ions. The same equilibration and production protocol was used for all the simulations. The system was initially submitted to 1000 steps of steepest descent algorithm followed by 7000 steps of conjugate gradient algorithm, with a harmonic potential constraint of 100 kcal·mol^-1^·A^-2^ applied to the proteins and the ligand. In the subsequent steps, the constraints were systematically reduced (harmonic potential of 10, 5, 2.5 and 0 kcal·mol^-1^·A^-2^, for each step) for 600 steps of conjugate gradient algorithm each time. After this cycle, Langevin thermostat was used to heat the whole system from 0 K to 100 K in the canonical ensemble (NVT), with a harmonic potential restraint of 20 kcal·mol^-1^·A^-2^ applied to the proteins and the ligand. This restraint was also maintained during the heating from 100 K to 300 K in the Isothermal-isobaric ensemble (NPT). Finally, all harmonic restraints were removed, and the system was submitted to a simulation of 100 ps. For the production run, Langevin thermostat in the NPT ensemble, at a 2 fs time step, was employed and the particle mesh Ewald (PME) method was used to calculate long-range electrostatic interactions as implemented in AMBER16.

#### MD simulations analysis

Trajectory analysis was performed using the cpptraj module of AmberTools15 for the calculation of RMSD and RMSF. MM-GBSA calculations were also performed to evaluate the differential affinity of each compound. These parameters, along with the information extracted from the visual inspection (VMD) ^133^ were used to analyse the simulations.

#### Thermal shift assay

A protein thermal shift dye kit (Applied Biosystems) was used, and the dye and reactions were prepared following manufacturer’s instructions. Melt curves were obtained in a QuantStudio^TM^ 5 Real-Time PCR (Thermo Fisher Scientific) with the decoupled filter m3(586±10) from 25°C to 99°C with a ramp rate of 0.03°C/s and sample wells assigned ROX reporter. OES2-0017, OES1-1087, and spermine were prepared as a 20X solution in a 3-fold dilution series. Reactions were composed of 5 μL protein thermal shift buffer (Applied Biosystems), 11.5 μL of protein solution containing 1.6 μM SpeG in water, 1 μL of 20X compound, and 2.5 μL of 8X protein thermal shift dye to reach a total volume of 20 μL. Protein thermal shift dye was diluted fresh before each reaction. Melt curves were analyzed in Microsoft Excel and GraphPad Prism 11 using the workflow support protocol 1 described in ^134^.

#### Acetyl-CoA quantification

The assay was modified from ^135^, to use SAT1 to convert cellular AcCoA to CoA for fluorescent detection. Overnight cultures grown in 5 mL of LB with 5 μg/mL erythromycin for *pdhA*::Tn in a 37 °C shaking incubator, 300 rpm were diluted to an OD_600_ of 0.05 in 50 mL fresh LB media containing 1/4 X MIC OES1-1087 (312.5 µM) or OES2-0017 (5 µM) or an equivalent volume of DMSO. Cultures were grown to an OD_600_ of 0.4-0.5 and cells were harvested by centrifugation at 10,000 xg for 10 minutes. Cell pellets were resuspended in 1 mL 1X PBS with 10 μg/mL lysostaphin (Sigma Aldrich) and incubated at 37°C for 30 minutes. Samples were centrifuged and the supernatant was retained, transferred to a microfuge tube, and placed on ice. Ice cold perchloric acid (Sigma Aldrich) was added to supernatants to a final concentration of 0.75 M and gently mixed. Ice cold potassium carbonate was added to a final concentration of 0.66M to neutralize the solution. Tubes were left open for 1 minute to allow the release of CO_2_ before vortexing for 1 minute. Samples were then centrifuged and the supernatant retained. A standard curve of Acetyl-CoA (Sigma Aldrich) was prepared and 50 μL of each concentration (0, 15.625, 31.25, 62.5, 125, 250, 500, and 1000 nM) was added to a black 96-well plate in duplicate. Samples were added to the plate in duplicate to a final volume of 50 μL; if dilutions were required, samples were added to the plate and diluted with H_2_O to a final volume of 50 μL. A 2X reaction buffer solution was prepared consisting of 100 mM HEPES, pH 8.0, 1 mM EDTA, 20 nM SAT1 and 120 μM spermine. 50 μL of the 2X reaction buffer was added to 50 μL of samples/standards in the plate. The plate was incubated at 30°C for 30 minutes then 100 μL of 10 μM 7-Diethylamino-3-(4-maleimidophenyl)-4-methylcoumarin (CPM; Sigma Aldrich), in DMSO, was added to each condition and incubated at room temperature for 30 minutes in the dark. Fluorescence was read at λex 390 nm λem 479 nm in a Neo2 Synergy BioTek plate reader set to extended gain. To account for any cellular CoA that might impact the signal, all samples were background subtracted by cell supernatant reacted with 2X reaction buffer without the addition of SAT1.

#### Laurdan assay

The protocol was adapted from ^136^. Overnight cultures of *S. aureus* USA300 were grown in LB at 37 °C in a shaking incubator set at 300 rpm. Cultures were diluted 1/100 into 10 mL fresh LB and grown to an OD_600_ 0.3-0.5. A final concentration of 10 μM Laurdan (dissolved in dimethylformamide; DMF) was added to cultures and incubated at 37 °C in a stationary incubator for 10 minutes. Cells were harvested by centrifugation at 37°C 16,000 xg for 1 minute. Cells were washed 4X with pre-warmed Laurdan buffer (137 mM sodium chloride, 2.7 mM potassium chloride, 10 mM potassium phosphate monobasic, 0.2% glucose, 1% DMF) and resuspended to an OD_600_ 0.8 in Laurdan Buffer. 100 μL of the stained cells were added to a 96-well black plate containing 100 μL of pre-warmed Laurdan buffer with 2X the desired compound concentration. The plate was immediately placed in a pre-warmed plate reader and incubated in the dark at 37 °C for 30-minutes followed by measuring fluorescence at λex 350 nm λem 460 and 500 nm. Data was analyzed by calculating the Laurdan generalized polarization (GP) using equation 4.

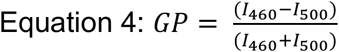

#### Identification of homologous acetyltransferases

*B. subtilis* BltD and PaiA protein sequences were used as a query in a Position-Specific Iterative Basic Local Alignment Search Tool (PSI-BLAST) search with results limited to records of *S. aureus* subsp. aureus USA300_FPR3757. Results were filtered based on alignment scores (E value).

#### Membrane permeabilization assay

Fluorometric Disc3(5) membrane permeabilization assays were performed as described in ^65^.

#### Phylogenetic analysis

Protein BLAST analysis was used to identify proteins with homology to *S. aureus* SpeG. Sequences were downloaded in FASTA format and a multiple sequence alignment was generated using MUSCLE ^137^. This analysis involved 94 amino acid sequences (Supplementary Table 4). A phylogenetic tree was constructed using a neighbor joining method ^138^ using MEGA11^139^ and visualized using iTOL ^140^. All ambiguous positions were removed for each sequence pair (pairwise deletion option). There were a total of 384 positions in the final dataset; positions represent the amino acid sites compared across all sequences. The evolutionary distances were computed using the p-distance method ^141^ and are in the units of the number of amino acid differences per site. The rate variation among sites was modeled with a gamma distribution (shape parameter = 1). The tree is the best estimate of the true evolutionary relationships among the analyzed sequences, given the available data and the neighbor-joining algorithm’s assumptions ^139^.

#### Macrophage infections

RAW-264.7 cells were seeded (1x10^6^ cells/mL) in RPMI (Sigma Aldrich) supplemented with 10% fetal bovine serum (Fisher Scientific), 1 mM HEPES (Thermofisher), and 100X MEM non-essential amino acids solution (Thermofisher) in 24-well plates 2 days prior to infection and incubated at 37°C and 5% CO_2_. *S.* Typhimurium 14028s wild-type and *speG*::Cm were grown in appropriate selection for 16 h at 37°C. Bacteria were subcultured in LB with no antibiotics until late log phase. The OD_600_ was normalized and cells washed with fresh media, 10 μL of washed cells were added to 1 mL of supplemented RPMI and 50 μL of the diluted bacteria was added to each well. Plates were centrifuged at 1000 rpm for 1 minute and incubated at 37°C, 5% CO_2_ for 10 minutes to complete infection. Media were replaced (1 mL/well) at ten minutes post-infection, with RPMI supplemented with 100 μg/mL gentamicin for 30 minutes to kill extracellular bacteria. After 30 minutes, fresh RPMI media supplemented with 10 μg/mL gentamicin with and without 10 μM OES2-0017 were added. Plates were then incubated for 4 hours. The pre-infection group was treated with 10μM OES2-0017 30 minutes prior to infection with *S.* Typhimurium. Cells were lysed by washing twice with 1 mL 1X PBS and then adding 1 mL 1% PBS-Triton for 5 minutes per well. Lysed cells were diluted and plated on LB agar and incubated at 37°C for 16 hours. Cells were lysed and plated at time 0, 4 hours post-infection, and 15 hours post-infection.

### Murine infection model

#### Mice

All work with mice was approved by the Canadian Council on Animal Care Committee under the University of Alberta Animal Use Protocol #3860. Female C57BL/6J mice (Jackson Laboratory) were bred in-house under specific-pathogen-free biocontainment level 2 conditions with 12 h light and dark cycles from 7 am to 7 pm, at 21-25°C and 45-55% humidity. Mice were provided a 4% fat (w/w) diet *ad libitum*.

#### Murine infections

C57BL/6J mice were orally gavaged with 20 mg/mL streptomycin 20 hours prior to oral infection with 10^8^ cfu/mL of *S. enterica* serovar Typhimurium strain 14028s. Mice were injected with 10 mg/kg OES1-1087 or saline (0.9% NaCl) i.p. 4-hours post infection and then each day post infection. Body weights were monitored daily, and animals were sacrificed 4 days post-infection. Spleen and liver tissues were harvested and plated on Luria Broth (LB) agar. Colony PCR with primers specific for *ssrB* was performed to validate the identity of recovered bacteria. Two independent trials were conducted with each trial consisting of 5 mice in the OES1-1087 treatment condition and 5 mice in the vehicle control condition.

#### Hemolysis Assay

The hemolysis assay was performed using defibrinated sheep blood (Thermo Fisher Scientific and Cedarlane) as previously described ^142^.

#### HepG2 cell culture and LDH assay

HepG2 hepatocarcinoma cells were cultured at 37°C in the presence of 5% CO_2_ in 1x MEM medium (Gibco) supplemented with the following additives: FBS (10% v/v), 1x MEM non-essential amino acid mix (Gibco), Penicillin-Streptomycin (Wisent) at 100 units/mL and 100 µg/mL respectively, Amphotericin B at 250 ng/mL (Gibco), and Sodium Pyruvate at 1mM (Wisent). HepG2 cells were passaged using trypsin and maintained in T25 tissue culture flasks. To measure compound toxicity, 3 T25 flasks of HepG2 cells were treated with trypsin and resuspended in the complete tissue culture medium described above. Cells from each flask were seeded into a 96 well clear tissue culture plate (Thermo Scientific Bio-Lite™) at ∼2.5x10^3^ cells per well and incubated for 48h. After this incubation period, the compounds of interest were diluted in complete tissue culture medium to their established MIC against *S. aureus* USA300 and then immediately added to the 96 well dish containing HepG2 cells in a 100 µL volume. As a vehicle control, DMSO alone was also used to treat cells. After 4h incubation, the cytotoxicity was measured using the LDH Cytotoxicity Assay Kit (Sigma, #MAK529) according to the manufacturer’s instruction with the following modifications: sterile 1x PBS was used in place of water for the no treatment control, the positive control agent Tergitol that causes cell death was added to select HepG2 containing wells the same time as the test compounds were added, and finally 140 µL of the reagent containing the tetrazolium salt was used. After 10 min incubation with the colorimetric reagent, the optical density was measured at 500 nm using a Biotek Epoch2 plate reader.

## Data Availability

Whole genome sequence data that support the findings of this study have been deposited in GenBank with accession numbers SAMN45063681 (ACJ4IB) for *sbnB*::Tn and SAMN45063680 (ACJ4HW) for its USA300 JE2 wild type control. The authors declare that the data supporting the findings of this study are available within the paper and its supplementary information files. Source data for all main and supplementary figures are provided with this paper.

## Acknowledgements

This work was supported by Natural Sciences and Engineering Research Council of Canada discovery grant (RGPIN-2022-04239 and DGECR-2022-00206), a Saskatchewan Health Research Foundation establishment grant (6115), a Canadian Institutes of Health Research grant (202403PPE-522800), and start-up funds from the Faculty of Science at the University of Regina to OME. OME holds a Canada Research Chair in Chemogenomics and Antimicrobial Research (CRC-2024-00304). PBM and JW were supported by a Canadian Institutes of Health Research Canada Graduate Scholarship—Master’s (CGS-M) award. PBM was also supported by the Verna Martin Memorial Scholarship and a Canadian Institutes of Health Research Canada Graduate Scholarship—Doctoral (CGS-D) award. WE holds a Canada Research Chair in Microbiome Research (CRC-00249). WE is funded by Crohn’s & Colitis Canada, Natural Sciences and Engineering Research Council of Canada, Major Innovation Fund, Antimicrobial Resistance -One Health Consortium, and Striving for Pandemic Preparedness - The Alberta Research Consortium (SPP-ARC). Work in the DEH laboratory was funded by an operating grant (PJT-183848) from the Canadian Institutes of Health Research. SMS and OMC were funded by the Spanish Ministry of Science, Innovation and Universities MICIU/AEI/10.13039/501100011033 (PID2023-152271NB-I00).

## Author contributions

O.M.E. conceived and coordinated the research. O.M.E., D.E.H., W.E., and S.M.S. designed and supervised the research and acquired the funding; P.B.M. performed all experiments and analyzed the data, with the following exceptions: R.S.F. performed and analyzed the cytotoxicity experiments. J.W. performed the macrophage and murine infections and J.W. and W.E. analyzed the data. O.M.C. performed the *in silico* computational experiments and O.M.C. and S.M.S. analyzed the data. A.M.S. preformed the phylogenetic analysis. O.M.E. and P.B.M. wrote the paper. The rest of the authors wrote the first draft of their respective sections. All authors commented on the manuscript. R.S.F. and J.W. contributed equally. D.E.H. and W.E. contributed equally.

