## Supplementary Tables and Figures for "Discovery of broad-spectrum bacterial polyamine detoxification inhibitors as potential antivirulence agents and antibiotic adjuvants"

Running title: Broad-spectrum polyamine detoxification inhibitors

**Supplementary Table 1. Query compounds for polyamine analog library construction.**

| Query Compound | Rationale | Reference |
|---|---|---|
| **Putrescine** | Polyamine | |
| **Spermidine** | Polyamine | |
| **Spermine** | Polyamine | |
| **Cadaverine** | Polyamine | |
| **Ornithine** | Precursor in biosynthetic pathway of polyamines | |
| **Arginine** | Precursor in biosynthetic pathway of polyamines | |
| **Agmatine** | Precursor in biosynthetic pathway of polyamines | |
| ***N*-carbamoylputrescine** | Precursor in biosynthetic pathway of polyamines | |
| ***N*-(1-amino-1-carboxy-2-ethyl)-glutamic acid** | Intermediate in SbnAB catalyzed reaction for the synthesis of L-2,3-diaminopropionic acid | 1 |
| **L-2-3-diaminopropionic acid** | Product of SbnB catalyzed reaction in *S. aureus* (initial chemogenomics screen suggested SbnB to be involved in polyamine response; however, we later ruled that out as *sbnB*::Tn spermine susceptibility was due to loss of *speG*) | 1 |
| **α-ketoglutarate** | Product of SbnB catalyzed reaction in *S. aureus* | 1 |
| ***N*8-acetylspermidine** | Product of SpeG catalyzed reaction | 2 |
| **N1,N4-bis(2,3-butadienyl)-1,4-butanediamine** | Polyamine oxidase inhibitor | 3 |
| **1,8-diaminooctane** | Polyamine oxidase inhibitor | 4 |
| **bis(ethyl)norspermine** | Polyamine metabolism inhibitor | 5 |
| **Diethylhomospermine** | Polyamine metabolism inhibitor | 5 |

**Supplementary Table 2. Polyamine analog library**

| Compound Name | SMILES | Solvent |
|---|---|---|
| **1-[3-(DIMETHYLAMINO)PROPYL]-3-ETHYLUREA** | N(CCCNC(=O)NCC)(C)C | DMSO |
| **N,N-Dimethyldipropylenetriamine** | N(CCCNCCCN)(C)C | DMSO |
| **4-(Diethylamino)butylamine** | N(CCCCN)(CC)CC | DMSO |
| **2-Hydroxybut-2-enoic acid** | OC(=O)C(=O)CC | DMSO |
| **2,6,10-Trimethyl-2,6,10-triazaundecane** | N(CCCN(C)C)(CCCN(C)C)C | DMSO |
| **3-(Butylamino)propionitrile** | N(CCCC)CCC#N | DMSO |
| **N-(4-Aminobutyl)acetamide** | N(CCCCN)C(=O)C | DMSO |
| **Dioctylamine** | N(CCCCCCCC)CCCCCCCC | DMSO |
| **L-Alanyl-l-glutamine** | N([C@@H](CCC(=O)N)C(=O)O)C(=O)[C@@H](N)C | $H_2O$ |
| **Cycloheptylamine** | NC1CCCCCC1 | DMSO |
| **3,3'-Iminodipropionitrile** | N(CCC#N)CCC#N | DMSO |
| **1-Allylpiperidine** | N1(CCCCC1)CC=C | DMSO |
| **(S)-5-Acetamido-2-amino-5-oxopentanoic acid** | N[C@@H](CCC(=O)NC(=O)C)C(=O)O | DMSO |
| **Ac-Arg-OH 2H2O** | N([C@@H](CCCNC(=N)N)C(=O)O)C(=O)C.O.O | $H_2O$ |
| **N-Ethyl-1,3-propanediamine** | N(CCCN)CC | DMSO |
| **1,3-Bis[3-(dimethylamino)propyl]urea** | N(CCCNC(=O)NCCCN(C)C)(C)C | DMSO |
| **N1-Dodecylpropane-1,3-diamine** | N(CCCN)CCCCCCCCCCCC | DMSO |
| **N,N-Dibutyl-1,3-propanediamine** | N(CCCN)(CCCC)CCCC | DMSO |
| **2-Amino-5-diethylaminopentane** | N(CCCC(N)C)(CC)CC | DMSO |
| **Dipropylenetriamine** | N(CCCN)CCCN | DMSO |
| **N,N'''-1,6-Hexanediylbis(N'-cyanoguanidine)** | N(CCCCCCNC(=N)NC#N)C(=N)NC#N | DMSO |
| **2-(Butylamino)ethylamine** | N(CCN)CCCC | DMSO |
| **Bis(hexamethylene)triamine** | N(CCCCCCN)CCCCCCN | DMSO |
| **Isoamylamine** | NCCC(C)C | DMSO |
| **butylurea** | N(CCCC)C(=O)N | DMSO |
| **dibutylamine** | N(CCCC)CCCC | DMSO |
| **N-(piperidin-3-ylmethyl)acetamide** | N1CC(CCC1)CNC(=O)C | DMSO |
| **N-butylimidodicarbonimidic diamide** | N(CCCC)C(=N)NC(=N)N | DMSO |
| **N''-[(E)-butyl]imidodicarbonimidic diamide** | N(\C(=N\CCCC)\N)C(=N)N | DMSO |
| **L-Alanine** | N[C@@H](C)C(=O)O | $H_2O$ |
| **Butyl(Ethyl)Amine** | N(CCCC)CC | DMSO |
| **pentan-1-amine** | NCCCCC | DMSO |
| **4-oxoheptanedioic acid** | OC(=O)CCC(=O)CCC(=O)O | $H_2O$ |

| | | |
|---|---|---|
| **4-methyl-2-oxopentanoic acid** | OC(=O)C(=O)CC(C)C | DMSO |
| **2-oxobutanedioic acid** | OC(=O)CC(=O)C(=O)O | DMSO |
| **hexane-1,6-diamine** | NCCCCCCN | DMSO |
| **(4-aminobutyl)dimethylamine** | N(CCCCN)(C)C | DMSO |
| **4-(pyrrolidin-1-yl)butan-1-amine** | N1(CCCC1)CCCCN | DMSO |
| **butan-1-amine** | NCCCC | DMSO |
| **(2S)-2-amino-4-carbamoylbutanoic acid** | N[C@@H](CCC(=O)N)C(=O)O | $H_2O$ |
| **3-aminopropanoic acid** | NCCC(=O)O | $H_2O$ |
| **(2S)-5-carbamimidamido-2-acetamidopentanoic acid** | N([C@@H](CCCNC(=N)N)C(=O)O)C(=O)C | DMSO |
| **(3-{[3-(dimethylamino)propyl]amino}propyl) dimethylamine** | N(CCCNCCCN(C)C)(C)C | DMSO |
| **propane-1,3-diamine** | NCCCN | DMSO |
| **(3-aminopropyl)(octyl)amine** | N(CCCN)CCCCCCCC | DMSO |
| **octan-1-amine** | NCCCCCCCC | DMSO |
| **hexan-1-amine** | NCCCCCC | DMSO |
| **bis(3-aminopropyl)(methyl)amine** | N(CCCN)(CCCN)C | DMSO |
| **(2-methylpropyl)urea** | N(CC(C)C)C(=O)N | DMSO |
| **pentane-1,3-diamine** | NC(CCN)CC | DMSO |
| **4-carbamimidamidobutanoic acid** | N(CCCC(=O)O)C(=N)N | $H_2O$ |
| **Dodecylamine** | NCCCCCCCCCCCC | DMSO |
| **2-Methylbutan-1-amine** | NCC(CC)C | DMSO |
| **Dimethyl 2-Oxoglutarate** | O(C)C(=O)CCC(=O)C(=O)OC | DMSO |
| **L-Theanine** | N[C@@H](CCC(=O)NCC)C(=O)O | $H_2O$ |
| **(S)-2-Piperazinecarboxylic Acid Dihydrochloride** | Cl.Cl.N1[C@@H](CNCC1)C(=O)O | $H_2O$ |
| **N-[6-(acetylamino)hexyl]acetamide** | N(CCCCCCNC(=O)C)C(=O)C | $H_2O$ |
| **N~1~,N~1~-bis[3-(dimethylamino)propyl]-N~3~,N~3~-dimethyl-1,3-propanediamine** | N(CCCN(C)C)(CCCN(C)C)CCCN(C)C | DMSO |
| **2-piperazinecarboxylic acid** | N1C(CNCC1)C(=O)O | $H_2O$ |
| **(R)-2-Amino-5-guanidinopentanoic acid** | N[C@H](CCCNC(=N)N)C(=O)O | DMSO |
| **3-(1-pyrrolidinyl)propylamine** | N1(CCCC1)CCCN | DMSO |
| **N~1~,N~4~-dimethyl-1,4-butanediamine** | N(CCCCNC)C | DMSO |
| **N~1~,N~3~-dimethyl-N~1~-[3-(methylamino)propyl]-1,3-propanediamine** | N(CCCNC)(CCCNC)C | DMSO |
| **(S)-2-Amino-6-guanidinohexanoic acid** | N[C@@H](CCCCNC(=N)N)C(=O)O | $H_2O$ |

| | | |
|---|---|---|
| **N~1~,N~3~-bis(2-aminoethyl)-1,3-propanediamine** | N(CCN)CCCNCCN | DMSO |
| **N1,N1-Bis(3-aminopropyl)propane-1,3-diamine** | N(CCCN)(CCCN)CCCN | DMSO |
| **tetrahydro-2(1H)-pyrimidinone** | N1CCCNC1=O | $H_2O$ |
| **N~1~,N~1~,N~3~-trimethyl-1,3-propanediamine** | N(CCCNC)(C)C | DMSO |
| **1,4-diazepane** | N1CCNCCC1 | DMSO |
| **N1,N1-Dimethyl-N3-propylpropane-1,3-diamine** | N(CCCNCCC)(C)C | DMSO |
| **N~1~,N~6~-dimethyl-1,6-hexanediamine** | N(CCCCCCNC)C | DMSO |
| **N,N'-Dimethyl-1,3-propanediamine** | N(CCCNC)C | DMSO |
| **(2S)-6-amino-2-[bis(carboxymethyl)amino]hexanoic acid** | N([C@@H](CCCCN)C(=O)O)(CC(=O)O)CC(=O)O | $H_2O$ |
| **[4-(aminomethyl)cyclohexyl]methylamine** | NCC1CCC(CC1)CN | DMSO |
| **Gly-Gln** | N([C@@H](CCC(=O)N)C(=O)O)C(=O)CN | $H_2O$ |
| **1,7-heptanediamine** | NCCCCCCCN | DMSO |
| **1-decanamine** | NCCCCCCCCCC | DMSO |
| **2-Amino-5-ureidopentanoic acid** | N(CCCC(N)C(=O)O)C(=O)N | DMSO |
| **1,4-cyclohexanediamine** | NC1CCC(CC1)N | DMSO |
| **N~1~,N~1~,N~4~,N~4~-tetramethyl-1,4-butanediamine** | N(CCCCN(C)C)(C)C | DMSO |
| **3-(1-piperidinyl)propylamine** | N1(CCCCC1)CCCN | DMSO |
| **Nomega-Nitro-L-arginine** | [N+](=O)(NC(=N)NCCC[C@H](N)C(=O)O)[O-] | $H_2O$ |
| **(2S)-2-aminohexanedioic acid** | N[C@@H](CCCC(=O)O)C(=O)O | $H_2O$ |

Supplementary Table 3. Calculated binding energies (kcal·mol$^{-1}$) of the top ranked docked pose of each ligand into the different binding sites of SpeG dimer and the spermine binding site of SAT1, calculated with AutoDock.

| | SpeG (8fv1) | | | SAT1 (2b58) |
|---|---|---|---|---|
| **Ligands** | **Allosteric site** | **Acceptor site** | **AcCoA binding site** | **Spermine binding site** |
| Spermine | -14.91 | -- | -- | -9.91 |
| OES2-0017 | -11.23 | -9.57 | -- | -5.97 |
| OES2-0045 | -7.14 | -- | -- | -5.57 |
| OES2-0046 | -4.56 | -- | -- | -4.33 |
| OES2-0047 | -4.05 | -- | -- | -4.03 |
| OES2-0052 | -7.14 | -- | -- | -4.90 |
| OES2-0077 | -6.19 | -- | -- | -4.62 |
| OES2-0085 | -8.44 | -- | -- | -5.73 |
| OES2-0086 | -7.7 | -- | -- | -6.02 |
| *R*-OES1-1087 | -- | -- | -4.63 | -- |
| *S*-OES1-1087 | -- | -- | -4.35 | -- |

Supplementary Table 4. Bacterial species and their accession numbers included in the phylogenetic tree presented in Supplementary figure 25.

| No. | Strain | Accession No. |
|---|---|---|
| **1** | *E. coli* strain K12 | NP_416101.1 |
| **2** | *Salmonella enterica* serovar Paratyphi A AKU12601 | CAR59427.1 |
| **3** | *Enterobacter hormaechei* RHBSTW-00564 | QLW45786.1 |
| **4** | *Listeria monocytogenes* 198 | KSZ46255.1 |
| **5** | *Vibrio cholerae* O1 KW3 | ALJ66217.1 |
| **6** | *Serratia marcescens* ID148138 | POX09291.1 |
| **7** | *Klebsiella quasipneumoniae* MMCC7 | UAD16483.1 |
| **8** | *Klebsiella variicola* NUKP-18 | GKI90423.1 |
| **9** | *Yersinia pestis* 9 | ERP82017.1 |
| **10** | *Staphylococcus aureus* USA300-ISMMS1 | AHJ05871.1 |
| **11** | *Staphylococcus aureus* USA300-0114 | CAC6174972.1 |
| **12** | *Staphylococcus aureus* USA300_FPR3757 | ABD21585 |
| **13** | *Staphylococcus aureus* NRS694 | CAC6299644.1 |
| **14** | *Citrobacter freundii* RHBSTW-00334 | QLY37052.1 |
| **15** | *Cronobacter sakazakii* G4023 | UEQ66194.1 |
| **16** | *Enterobacter cloacae* RHBSTW-00399 | QLW21244.1 |
| **17** | *Burkholderia multivorans* 2020Y71628512IV | UQP68389.1 |
| **18** | *Klebsiella aerogenes* RHBSTW-00938 | QMR42362.1 |
| **19** | *Vibrio parahaemolyticus* BB22OP | AGB11804.1 |
| **20** | *Yersinia enterocolitica* YE-P4 | EOR64463.1 |
| **21** | *Klebsiella oxytoca* RHBSTW-00432 | QLU24515.1 |
| **22** | *Klebsiella michiganensis* RHBSTW-00409 | QLO24962.1 |
| **23** | *Proteus mirabilis* DP2019 | UZK71462.1 |
| **24** | *Bacillus thuringiensis* UFT038 | UOB63615.1 |
| **25** | *Burkholderia pseudomallei* MSHR1046 | UZU29773.1 |
| **26** | *Enterobacter kobei* STW0522-60 | BBV81867.1 |
| **27** | *Enterobacter asburiae* JBIWA007 | UAN34532.1 |
| **28** | *Yersinia ruckeri* 17Y0159 | UIM99950.1 |
| **29** | *Escherichia albertii* 147_1_TBG_A | UUL20973.1 |
| **30** | *Aeromonas hydrophila* AG-2013-AG1 | WEF00472.1 |
| **31** | *Bacillus mycoides* JAS23/1 | QWI13605.1 |
| **32** | *Raoultella ornithinolytica* NCTC9164 | WP_004862437.1 |

| | | |
|---|---|---|
| **33** | *Escherichia fergusonii* RHB03-C23 | QLM34977.1 |
| **34** | *Morganella morganii* K266 | UVZ52482.1 |
| **35** | *Yersinia pseudotuberculosis* FDAARGOS_584 | AYW87375.1 |
| **36** | *Burkholderia cenocepacia* C6433 | USB83525.1 |
| **37** | *Burkholderia cenocepacia* K56-2Valvano | EPZ91470.1 |
| **38** | *Burkholderia cenocepacia* J2315 | CAR50744.1 |
| **39** | *Shigella sonnei* ECFood+09 | PBP11012.1 |
| **40** | *Pantoea ananatis* OC5a | QTC47233.1 |
| **41** | *Bacillus albus* PG 17 | RXJ29246.1 |
| **42** | *Citrobacter k*oseri TUM13189 | BDG84699.1 |
| **43** | *Vibrio alginolyticus* AUSMDU00064140 | WAE59614.1 |
| **44** | *Shigella flexneri* SWHIN_101 | ULK85562.1 |
| **45** | *Providencia rettgeri* 18004577 | URR22833.1 |
| **46** | SAT1_HUMAN | NP_002961.1 |
| **47** | *Shewanella algae* A291 | QTE95040.1 |
| **48** | *Staphylococcus epidermidis* NCTC13360 | SUM28383.1 |
| **49** | *Hafnia alvei* H4 | KAA0260357.1 |
| **50** | *Lactococcus lactis* LMG9447 | KSU20519.1 |
| **51** | *Mus musculus*-SAT1_MOUSE | P48026.1 |
| **52** | *Achromobacter xylosoxidans* PartM-Axylosoxidans-RM8376 | UON37785.1 |
| **53** | *Shigella boydii* 183 | QGU65304.1 |
| **54** | *Providencia stuartii* CAVP490 | WER28085.1 |
| **55** | *Shigella dysenteriae* SWHEFF_49 | ULK13713.1 |
| **56** | *Edwardsiella tarda* SC002 | WGE28148.1 |
| **57** | *Listeria innocua* ATCC 33090 | WCT28232.1 |
| **58** | Diamine acetyltransferase 1 *Salmo salar* | ACN12572.1 |
| **59** | *Methylobacterium oxalidis* NBRC 107715 | GLS63018.1 |
| **60** | *Proteus vulgaris* FDAARGOS_1507 | UBH60500.1 |
| **61** | *Bordetella hinzii* 2B3 | QWF42849.1 |
| **62** | *Nocardia farcinica* W6977 | AXK84801.1 |
| **63** | *Staphylococcus warneri* NCTC11044 | VED76743.1 |
| **64** | *Staphylococcus haemolyticus* NBRC 109768 | GEQ09032.1 |
| **65** | spermidine/spermine N1-acetyltransferase 1 [*Xenopus laevis*] | DAA34838.1 |
| **66** | *Staphylococcus xylosus* NBRC 109770 | GEQ09573.1 |

| | | |
|---|---|---|
| **67** | *Staphylococcus gallinarum* SNUC 1046 | RIL33103.1 |
| **68** | *Staphylococcus hominis* As1 | KMU57174.1 |
| **69** | *Staphylococcus lugdunensis* NCTC12217 | SQI97098.1 |
| **70** | *Staphylococcus pragensis* CCM 8529 | GGG93374.1 |
| **71** | *Staphylococcus capitis* C87 | EFS18284.1 |
| **72** | *Cyanobium* sp. ULC084bin3 | PZU96833.1 |
| **73** | *Cricetulus griseus* Chinese hamster | AAF86286.1 |
| **74** | *Nocardia asteroids* FDAARGOS_1620 | UGT61696.1 |
| **75** | SAT1_*Mus musculus*_Mouse | NP_033147.1 |
| **76** | SAT1_ *Sus scrofa*_Pig | NP_999523.1 |
| **77** | SAT1_*Gallus gallus*_CHICK | NP_989517.1 |
| **78** | *Staphylococcus cohnii* NBRC 109713 | GEP86880.1 |
| **79** | *Staphylococcus aureus* NRS702 | CAC6336879.1 |
| **80** | *Saccharomyces cerevisiae* S288C | DAA11917.1 |
| **81** | *Candida railenensis* CLIB 1423 | CAH2354588.1 |
| **82** | *Candida albicans* SC5314 | AOW27049.1 |
| **83** | *Aspergillus awamori* IFM 58123 | GCB18706.1 |
| **84** | *Aspergillus nidulans* FGSC A4 | XP_050467072.1 |
| **85** | SAT1_ *Bos taurus*_BOVIN | NP_001029505.1 |
| **86** | *Acinetobacter baumannii* 35 Dpa | WHT50386.1 |
| **87** | *Methanobrevibacter curvatus* DSM 11111 | KZX12885.1 |
| **88** | *Candidatus Lokiarchaeota* BC3 archaeon | TXT64754.1 |
| **89** | *Thermoplasma acidophilum* CF.Pisc.bin.16 | MCY0851537.1 |
| **90** | *Klebsiella pneumoniae* MKP103 | WP_002903719.1 |
| **91** | *Salmonella enterica* subsp. *enterica* serovar Typhimurium str. 14028S chromosome | ACY88287.1 |
| **92** | spermidine acetyltransferase [*Pseudomonas aeruginosa* PAO1]PA4114 | NP_252803.1 |
| **93** | hypothetical protein PA1377 [*Pseudomonas aeruginosa* PAO1] | NP_250068.1 |
| **94** | hypothetical protein PA1472 [*Pseudomonas aeruginosa* PAO1] | NP_250163.1 |

Supplementary Table 5. Energy terms (kcal·mol$^{-1}$) calculated as the mean of the MM-GBSA computed for each replica.

| | OES2-0017 | | OES2-0086 | |
|---|---|---|---|---|
| | **Mean** | **Std. Dev.** | **Mean** | **Std. Dev.** |
| **SpeG** | -46.53 | 7.49 | -40.8 | 8.97 |
| **SAT1** | -70.145 | 8.77 | -56.86 | 8.36 |

Supplementary Table 6. Strains and plasmids used in this study:

| Bacterial Strain | Description | Reference |
|---|---|---|
| ***Staphylococcus aureus*** | | |
| **USA300** | Community associated MRSA USA300 LAC isolated from a skin and soft tissue infection, cured of plasmids, known as JE2 | Lab stock* |
| **RN4220** | Restriction endonuclease deficient lab strain | Lab stock |
| **Newman** | MSSA lab strain | Lab stock |
| **COL** | MRSA lab strain | Lab stock |
| **NCTC8325** | MSSA lab strain | Lab stock |
| **CMRSA-10** | MRSA USA300 strain | Lab stock |
| **Nebraska Transposon Mutant Library** | Nebraska Transposon Mutant Library Screening Array; 1920 *S. aureus* subsp. aureus USA300 JE2, transposon (Tn) mutants arrayed in five 384-well microtiter plates. ErmR | [10]* |
| **USA300 Δ*speG*** | USA300 with *speG* deletion created with pJB38 mutagenesis | This study |
| **USA300 Δ*sbnB*** | USA300 with *sbnB* deletion created with pJB38 mutagenesis | This study |
| **USA300 + CV** | USA300 with pKK22 control vector | Lab stock |
| **Δ*speG* + CV** | USA300 Δ*speG* mutant with pKK22 control vector | This study |
| **Δ*speG* + p*speG*** | USA300 Δ*speG* mutant with *speG* cloned into pKK22 under the expressioin of Pfba | This study |
| ***Escherichia coli*** | | |
| **DH5α** | Lab strain engineered for transformation efficiency | Lab stock |
| **DH5α λ pir** | Lab strain engineered to support the replication of plasmids with an R6K origin of replication. | Lab stock |

| | | |
|---|---|---|
| **DC10B** | Lab strain optimized for transforming DNA into *S. aureus* | Lab stock* |
| **BL21** | Lab strain with T7 RNA polymerase for expressing recombinant proteins | Lab stock |
| **K-12 BW25113** | | Lab stock |
| **K-12 BW25113 Δ*speG*** | *speG* mutants of *E. coli* K-12 from nonessential single-gene deletion library | [11] |
| ***Klebsiella pneumoniae*** | | |
| **MKP103** | | Lab stock |
| **MKP103 *speG*::Tn30** | *speG* transposon mutant from sequence-defined transposon mutant library of *K. pneumoniae* strain KPNIH1 | [12] |
| ***Salmonella enterica* serovar Typhimurium 14028s** | Laboratory strain isolated from chicken heart and liver | Lab stock |
| ***S.* Typhimurium *speG*::Cm** | *speG* gene disruption from sequence-defined single-gene deletion mutant library | [13] |
| ***Bacillus subtilis* 168** | | |
| **CRISPRi collection** | *B. subtilis* CRISPRi essential gene knockdown strain collection | [14] |
| ***ΔpaiA*** | *B. subtilis* 168 containing a deletion in *paiA,* KanR | [15] |
| ***ΔbltD*** | *B. subtilis* 168 containing a deletion in *bltD,* KanR | [15] |
| ***Enterococcus*** | | |
| ***E. faecalis* TX1322** | Clinical isolate, isolated from feces | Lab stock* |
| ***E. faecium* ERV165** | Clinical isolate, isolated from feces | Lab stock* |
| ***Staphylococcus epidermidis*** | | |
| **NRS6** | Clinical isolate | Lab stock* |
| **NIH04008** | Clinical isolate | Lab stock* |

| | | |
|---|---|---|
| ***Burkholderia cenocepacia* K56-2** | Cystic fibrosis clinical isolate, isolated from sputum | Lab stock |
| ***Mycobacterium smegmatis* ATCC607** | Human isolate | Lab stock |
| ***Candida albicans* SC5314** | Lab strain | Lab stock |
| ***Acinetobacter baumannii* ATCC19606** | Clinical isolate from human urine | Lab stock |

| **Plasmid** | **Description** | **Reference** |
|---|---|---|
| **pJB38** | *S. aureus* temperature-sensitive allelic exchange plasmid | [16*] |
| **pJB38-Δ*speG*** | pPM3; Vector for allelic replacement of *speG* | This study |
| **pJB38-Δ*sbnB*** | pPM2; Vector for allelic replacement of *sbnB* | This study |
| **pKK22 Pfba** | pKK22 complementation vector with *S. aureus fba* promoter with trimethoprim resistance. | Lab stock |
| **pKK22 Pfba-*speG*** | pKK22 complementation vector with *S. aureus speG* cloned downstream of the *S. aureus fba* promoter. | This study |
| **pKK22 Pfba-*pdhA*** | pKK22 complementation vector with *S. aureus pdhA* cloned downstream of the *S. aureus fba* promoter. | This study |
| **pKK22 Pfba-*lpdA*** | pKK22 complementation vector with *S. aureus lpdA* cloned downstream of the *S. aureus fba* promoter. | This study |
| **pET28a(+)** | IPTG-inducible expression vector with a 6X His tag and kanamycin resistance. | Lab stock |
| **pET28a(+)-*speG*** | pPM6; Vector for expression and purification of SpeG | This study |
| **pET28a(+)-*SAUSA300_2083*** | pPM11; Vector for expression and purification of SAUSA300_2083 | This study |
| **pET28a(+)-*paiA*** | pPM12; Vector for expression and purification of PaiA | This study |

| | | |
|---|---|---|
| **pET28a(+)-SAT1** | pPM15; Vector for the expression and purification of SAT1 | This study |
| **pET28a(+)-*SAUSA300_0441*** | pPM16; Vector for the expression and purification of SAUSA300_0441 | This study |

*Obtained from BEI Resources, NIAID, NIH.

**Supplementary Table 7.** Primers used in this study

| Primer Name | Sequence |
|---|---|
| **SpeG_del_US_F_EcoRI** | tataGAATTCcctgatatccccaagtgatga |
| **SpeG_del_US_R_BamHI** | tataGGATCCtcataaggctcttcaaacca |
| **SpeG_del_DS_F_BamHI** | tataGGATCCcgagtcacaaggatttaaaacaga |
| **SpeG_del_DS_R_SalI** | tataGTCGACaactcggcagttcaaatgatg |
| **SpeG_del_Check_F** | Gaatttgggctgtattcacca |
| **SpeG_del_Check_R** | Ccatgggacaactcgcttta |
| **sbnB_del_US_F_EcoRI** | tatagaattcaggaagcgcttttgattgaa |
| **sbnB_del_US_R_BamHI** | tataggatccatgattacctcccgcttgttc |
| **sbnB_del_DS_F_BamHI** | tataggatccggacgtgaagacgatgatga |
| **sbnB_del_DS_R_SalI** | tatagtcgactatacggtgcgccaatatca |
| **sbnB_del_check_F** | gaatcattatcaattaggtaacacaca |
| **sbnB_del_check_R** | atgactgcattgcaaatggta |
| **pJB38_check_F** | TGCCACCTGACGTCTAAGAA |
| **pJB38_check_R** | TTTGCGCTTAAAACCAGTCA |
| **speG_comp_F_AvrII** | tata**CCTAGG**atgaaactaagagcattagagtatagtgatt |
| **speG_comp_R_SacI** | gcgc**GAGCTC**ctacaaaatatattcagatttcaataatg |
| **pdhA_comp_F_AvrII** | GTGTCCTAGGATGGCTCCTAAGTTACAAG |
| **pdhA_comp_R_SacI** | TATAGAGCTCTTACTTCGACTCCTTCTCTTTGTAAA |
| **lpdA_comp_F_AvrII** | TATACCTAGGATGGTAGTTGGAGATTTCCCAATTG |
| **lpdA_comp_R_SacI** | GTGTGAGCTCTTACATTGTATGGATTGGGT |
| **pKK22_Check_F** | CGCATACATTCTTACATTTAGTGC |
| **pKK22_Check_R** | CCATCAATTTTAACAATCACCGC |
| **T7 promoter** | TAATACGACTCACTATAGGG |
| **T7 terminator** | GCTAGTTATTGCTCAGCGG |
| **speG_ pET28a_F_NdeI** | gtctCATATGatgaaactaagagcattagagtatagtgatt |
| **speG_pET28a_R_EcoRI** | gctataGAATTCctacaaaatatattcagatttcaataatg |
| **2083_pET28a_F_NdeI** | gcgcCATATGgctaaagaactttgttttga |
| **2083_pET28a_R_EcoRI** | gcgcGAATTCttagtaatttaaatatttttcgaaaacgac |
| **paiA_pET28a_F _NdeI** | gcgcCATATGgctgggataatcaaaga |

| | |
|---|---|
| **paiA_pET28a_R_EcoRI** | gcgcGAATTCttataactctttttccataattaaatctgtatc |
| **0441_pET28a_F_NdeI** | gatacatATGACAAATAATGACACCATCATGTTAC |
| **0441_pET28a_R_EcoRI** | tatagaattcTTATTTTATAGTTAACGCCATAATATGAACAGG |
| **ssrB_F** | GACTACAAAGACCATGACGGTGATTATAAAGATCATGACATCGATTACAAGGATG |
| **ssrB_R** | ATGATGCGGTCTCGAGTCATACTCTATTAACCTCATTCTTCGG |

**Supplementary Table 8.** High-throughput screening of chemical libraries against *S. aureus* USA300 to identify inhibitors of polyamine detoxification.

| Category | Spectrum Collection Screen | Polyamine analog screen |
|---|---|---|
| **Assay** | Type of Assay: Whole cell *S. aureus* USA300 | Type of Assay: Whole cell *S. aureus* USA300 |
| | Target: Polyamine detoxification mechanisms | Target: Polyamine detoxification mechanisms |
| | Primary Measurement: Detection of growth by OD600 | Primary Measurement: Detection of growth by OD600 |
| | Key reagents: Mueller Hinton Broth containing 2.5 mM spermine (Sigma) | Key reagents: Mueller Hinton Broth containing 5 mM spermine (Sigma) |
| | Assay protocol: see methods section Chemical screen and polyamine analog assembly | Assay protocol: see methods section Chemical screen and polyamine analog assembly |
| **Library** | Library composition: 2560 compounds comprised of previously approved drugs and natural product derivatives | Library composition: 83 compounds comprised of polyamine analogs with at least 60% structural similarity to substrates and products of polyamine biosynthesis and detoxification enzymes in bacteria. See supplementary tables 1 and 2 |
| | Source: Spectrum collection, MicroSource Discovery Systems, Inc. | Source: Sigma Aldrich Market Select |
| **Screen** | Format: 384-well non-treated, rounded square-well, clear, flat bottom microplates (Nunc) | Format: 96-well non-treated, round, clear, flat bottom microplates (Nunc) |
| | Concentration(s) tested: 20 $\mu$M 0.2% final concentration DMSO | Concentration(s) tested: 20 $\mu$M 1% final concentration DMSO |
| | Plate controls: 0.2% DMSO | Plate controls: 1% DMSO |
| | Reagent/ compound dispensing system: Biomatrix BM6-BC (S&P Robotics inc.) | Reagent/ compound dispensing system: Eppendorf multichannel pipette |
| | Detection instrument and software: Biotek Synergy Neo2 plate reader, Gen5 software. | Detection instrument and software: Biotek Synergy Neo2 plate reader, Gen5 software. |
| | Assay validation/QC: Z' score [1] | Assay validation/QC: Z' score [1] |
| | Normalization: percent growth relative to untreated controls | Normalization: percent growth relative to untreated controls |
| **Post-HTS analysis** | Hit criteria: at least 80% growth inhibition relative to growth control, and exclusion of known antimicrobial compounds. | Hit criteria: at least 80% growth inhibition relative to growth control. |
| | Hit rate: 4.92% (126/2560 compounds) | Hit rate: 3.61% (3/83 compounds) |
| | Additional assay(s): dose-response assays | Additional assay(s): dose-response assays |
| | Confirmation of hit purity and structure: compounds resupplied from commercial provider and $^{13}C$ and $^{1}H$ NMR performed on the hit used in mode of action determination assays. | Confirmation of hit purity and structure: compounds resupplied from commercial provider and $^{13}C$ and $^{1}H$ NMR performed on the hit used in mode of action determination assays. |

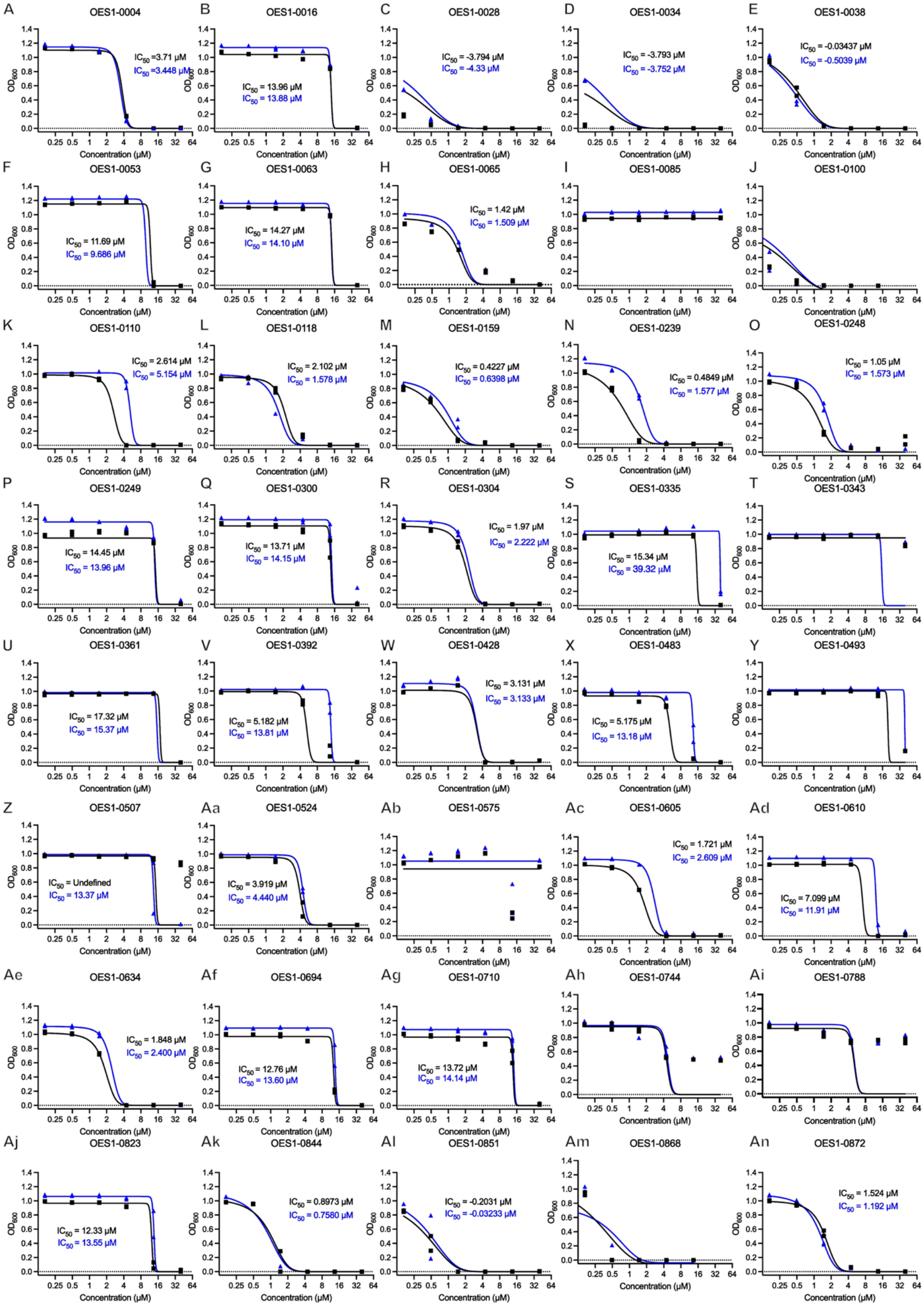

A OES1-0004
IC50 =3.71 µM
IC50 =3.448 µM
B OES1-0016
IC50 = 13.96 µM
IC50 = 13.88 µM
C OES1-0028
IC50 = -3.794 µM
IC50 = -4.33 µM
D OES1-0034
IC50 = -3.793 µM
IC50 = -3.752 µM
E OES1-0038
IC50 = -0.03437 µM
IC50 = -0.5039 µM
F OES1-0053
IC50 = 11.69 µM
IC50 = 9.686 µM
G OES1-0063
IC50 = 14.27 µM
IC50 = 14.10 µM
H OES1-0065
IC50 = 1.42 µM
IC50 = 1.509 µM
I OES1-0085
J OES1-0100
K OES1-0110
IC50 = 2.614 µM
IC50 = 5.154 µM
L OES1-0118
IC50 = 2.102 µM
IC50 = 1.578 µM
M OES1-0159
IC50 = 0.4227 µM
IC50 = 0.6398 µM
N OES1-0239
IC50 = 0.4849 µM
IC50 = 1.577 µM
O OES1-0248
IC50 = 1.05 µM
IC50 = 1.573 µM
P OES1-0249
IC50 = 14.45 µM
IC50 = 13.96 µM
Q OES1-0300
IC50 = 13.71 µM
IC50 = 14.15 µM
R OES1-0304
IC50 = 1.97 µM
IC50 = 2.222 µM
S OES1-0335
IC50 = 15.34 µM
IC50 = 39.32 µM
T OES1-0343
U OES1-0361
IC50 = 17.32 µM
IC50 = 15.37 µM
V OES1-0392
IC50 = 5.182 µM
IC50 = 13.81 µM
W OES1-0428
IC50 = 3.131 µM
IC50 = 3.133 µM
X OES1-0483
IC50 = 5.175 µM
IC50 = 13.18 µM
Y OES1-0493
Z OES1-0507
IC50 = Undefined
IC50 = 13.37 µM
Aa OES1-0524
IC50 = 3.919 µM
IC50 = 4.440 µM
Ab OES1-0575
Ac OES1-0605
IC50 = 1.721 µM
IC50 = 2.609 µM
Ad OES1-0610
IC50 = 7.099 µM
IC50 = 11.91 µM
Ae OES1-0634
IC50 = 1.848 µM
IC50 = 2.400 µM
Af OES1-0694
IC50 = 12.76 µM
IC50 = 13.60 µM
Ag OES1-0710
IC50 = 13.72 µM
IC50 = 14.14 µM
Ah OES1-0744
Ai OES1-0788
Aj OES1-0823
IC50 = 12.33 µM
IC50 = 13.55 µM
Ak OES1-0844
IC50 = 0.8973 µM
IC50 = 0.7580 µM
Al OES1-0851
IC50 = -0.2031 µM
IC50 = -0.03233 µM
Am OES1-0868
An OES1-0872
IC50 = 1.524 µM
IC50 = 1.192 µM
OD600
Concentration (µM)

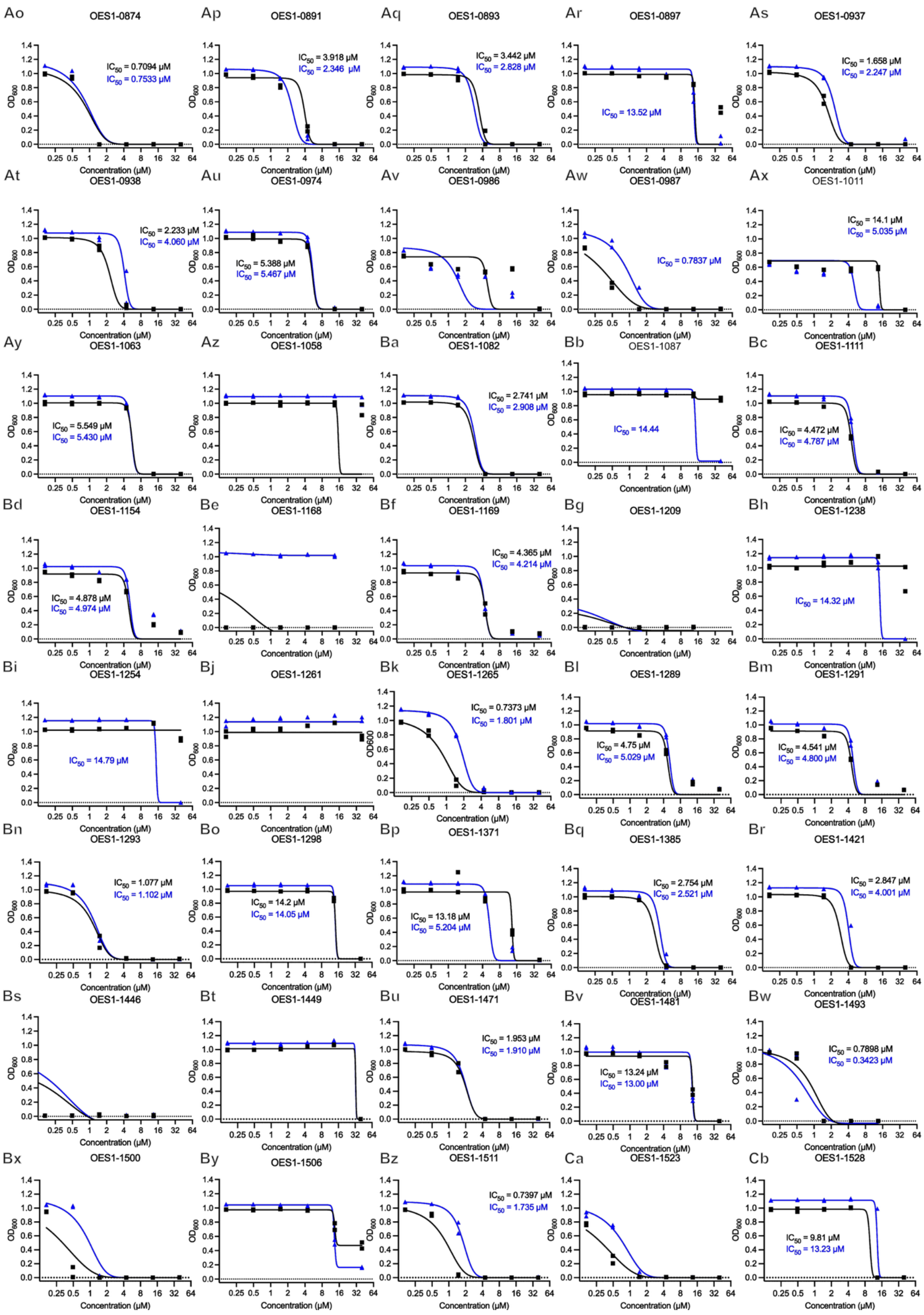
Ao
OES1-0874
IC50 = 0.7094 μM
IC50 = 0.7533 μM
Ap
OES1-0891
IC50 = 3.918 μM
IC50 = 2.346 μM
Aq
OES1-0893
IC50 = 3.442 μM
IC50 = 2.828 μM
Ar
OES1-0897
IC50 = 13.52 μM
As
OES1-0937
IC50 = 1.658 μM
IC50 = 2.247 μM
At
OES1-0938
IC50 = 2.233 μM
IC50 = 4.060 μM
Au
OES1-0974
IC50 = 5.388 μM
IC50 = 5.467 μM
Av
OES1-0986
Aw
OES1-0987
IC50 = 0.7837 μM
Ax
OES1-1011
IC50 = 14.1 μM
IC50 = 5.035 μM
Ay
OES1-1063
IC50 = 5.549 μM
IC50 = 5.430 μM
Az
OES1-1058
Ba
OES1-1082
IC50 = 2.741 μM
IC50 = 2.908 μM
Bb
OES1-1087
IC50 = 14.44
Bc
OES1-1111
IC50 = 4.472 μM
IC50 = 4.787 μM
Bd
OES1-1154
IC50 = 4.878 μM
IC50 = 4.974 μM
Be
OES1-1168
Bf
OES1-1169
IC50 = 4.365 μM
IC50 = 4.214 μM
Bg
OES1-1209
Bh
OES1-1238
IC50 = 14.32 μM
Bi
OES1-1254
IC50 = 14.79 μM
Bj
OES1-1261
Bk
OES1-1265
IC50 = 0.7373 μM
IC50 = 1.801 μM
Bl
OES1-1289
IC50 = 4.75 μM
IC50 = 5.029 μM
Bm
OES1-1291
IC50 = 4.541 μM
IC50 = 4.800 μM
Bn
OES1-1293
IC50 = 1.077 μM
IC50 = 1.102 μM
Bo
OES1-1298
IC50 = 14.2 μM
IC50 = 14.05 μM
Bp
OES1-1371
IC50 = 13.18 μM
IC50 = 5.204 μM
Bq
OES1-1385
IC50 = 2.754 μM
IC50 = 2.521 μM
Br
OES1-1421
IC50 = 2.847 μM
IC50 = 4.001 μM
Bs
OES1-1446
Bt
OES1-1449
Bu
OES1-1471
IC50 = 1.953 μM
IC50 = 1.910 μM
Bv
OES1-1481
IC50 = 13.24 μM
IC50 = 13.00 μM
Bw
OES1-1493
IC50 = 0.7898 μM
IC50 = 0.3423 μM
Bx
OES1-1500
By
OES1-1506
Bz
OES1-1511
IC50 = 0.7397 μM
IC50 = 1.735 μM
Ca
OES1-1523
Cb
OES1-1528
IC50 = 9.81 μM
IC50 = 13.23 μM
OD600
Concentration (μM)

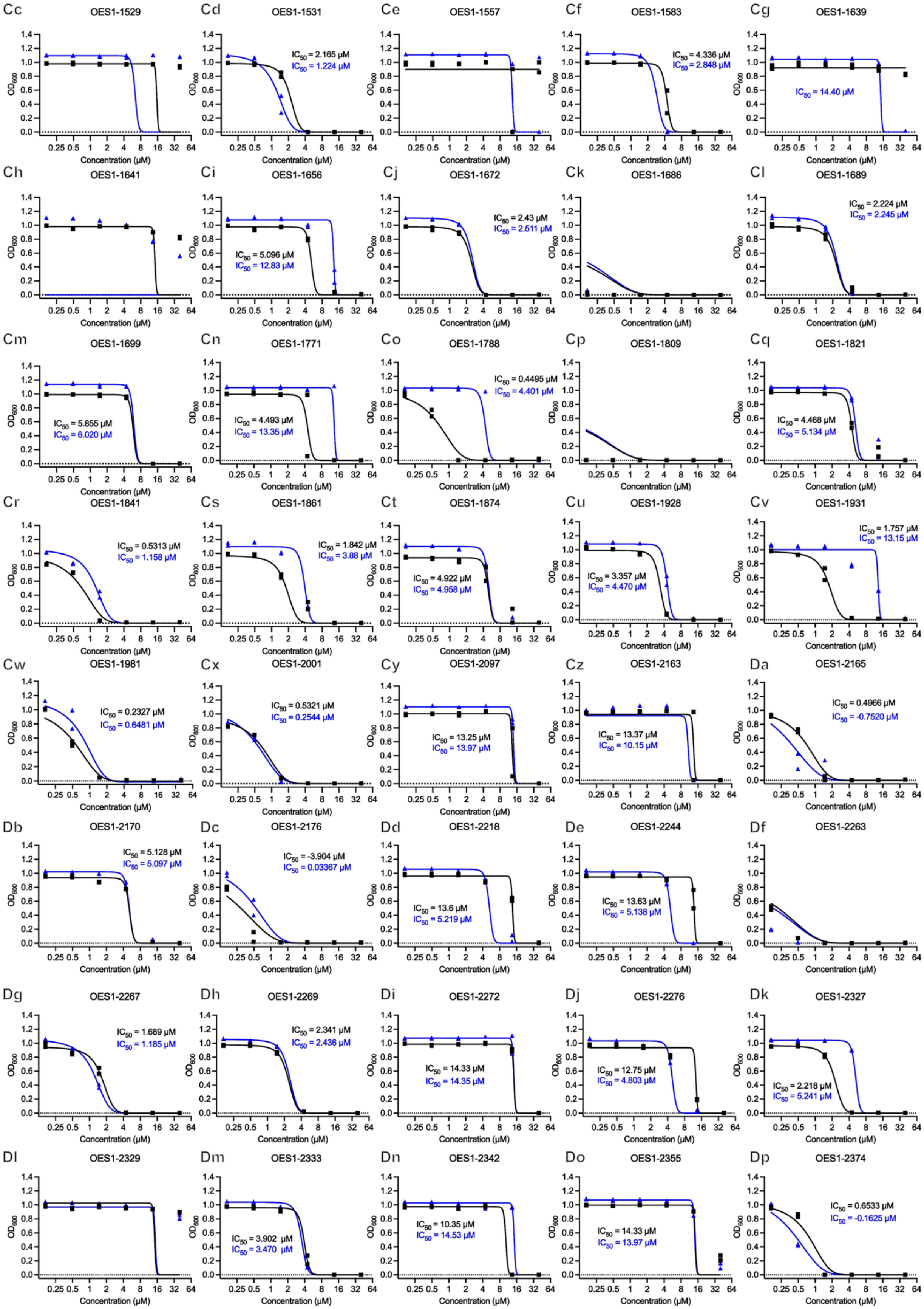
Cc OES1-1529
Cd OES1-1531
IC50 = 2.165 µM
IC50 = 1.224 µM
Ce OES1-1557
Cf OES1-1583
IC50 = 4.336 µM
IC50 = 2.848 µM
Cg OES1-1639
IC50 = 14.40 µM
Ch OES1-1641
Ci OES1-1656
IC50 = 5.096 µM
IC50 = 12.83 µM
Cj OES1-1672
IC50 = 2.43 µM
IC50 = 2.511 µM
Ck OES1-1686
Cl OES1-1689
IC50 = 2.224 µM
IC50 = 2.245 µM
Cm OES1-1699
IC50 = 5.855 µM
IC50 = 6.020 µM
Cn OES1-1771
IC50 = 4.493 µM
IC50 = 13.35 µM
Co OES1-1788
IC50 = 0.4495 µM
IC50 = 4.401 µM
Cp OES1-1809
Cq OES1-1821
IC50 = 4.468 µM
IC50 = 5.134 µM
Cr OES1-1841
IC50 = 0.5313 µM
IC50 = 1.158 µM
Cs OES1-1861
IC50 = 1.842 µM
IC50 = 3.88 µM
Ct OES1-1874
IC50 = 4.922 µM
IC50 = 4.958 µM
Cu OES1-1928
IC50 = 3.357 µM
IC50 = 4.470 µM
Cv OES1-1931
IC50 = 1.757 µM
IC50 = 13.15 µM
Cw OES1-1981
IC50 = 0.2327 µM
IC50 = 0.6481 µM
Cx OES1-2001
IC50 = 0.5321 µM
IC50 = 0.2544 µM
Cy OES1-2097
IC50 = 13.25 µM
IC50 = 13.97 µM
Cz OES1-2163
IC50 = 13.37 µM
IC50 = 10.15 µM
Da OES1-2165
IC50 = 0.4966 µM
IC50 = -0.7520 µM
Db OES1-2170
IC50 = 5.128 µM
IC50 = 5.097 µM
Dc OES1-2176
IC50 = -3.904 µM
IC50 = 0.03367 µM
Dd OES1-2218
IC50 = 13.6 µM
IC50 = 5.219 µM
De OES1-2244
IC50 = 13.63 µM
IC50 = 5.138 µM
Df OES1-2263
Dg OES1-2267
IC50 = 1.689 µM
IC50 = 1.185 µM
Dh OES1-2269
IC50 = 2.341 µM
IC50 = 2.436 µM
Di OES1-2272
IC50 = 14.33 µM
IC50 = 14.35 µM
Dj OES1-2276
IC50 = 12.75 µM
IC50 = 4.803 µM
Dk OES1-2327
IC50 = 2.218 µM
IC50 = 5.241 µM
Dl OES1-2329
Dm OES1-2333
IC50 = 3.902 µM
IC50 = 3.470 µM
Dn OES1-2342
IC50 = 10.35 µM
IC50 = 14.53 µM
Do OES1-2355
IC50 = 14.33 µM
IC50 = 13.97 µM
Dp OES1-2374
IC50 = 0.6533 µM
IC50 = -0.1625 µM
OD600
Concentration (µM)

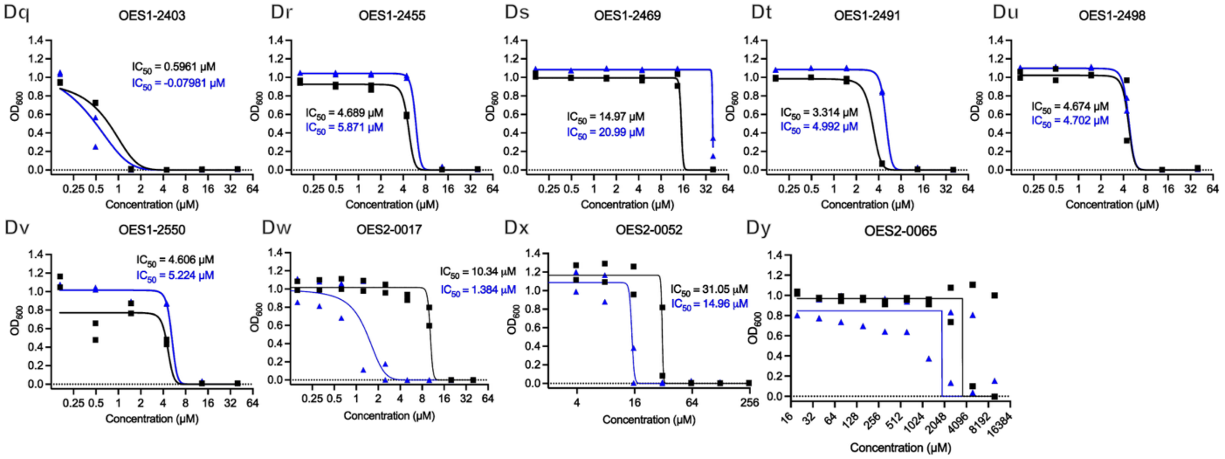


Supplementary Figure 1. **Dose-response assays of hits from chemical screen identifies 8 compounds with reduced MIC in the presence of spermine. A-Dy)** Dose response assays of compounds from the chemical screen against USA300 in the presence of a sub-inhibitory concentration of spermine (2.5 mM, blue triangles) and in the absence of spermine (black circles). Compounds showing ≥ 2-fold decrease in $IC_{50}$ in the presence of spermine compared to in its absence or an $IC_{50}$ in the spermine condition and no defined $IC_{50}$ in the absence of spermine in at least one replicate, with no antimicrobial or antiseptic activity reported in literature, and that are commercially available were considered hits. The 8 hits are OES1-0507, OES1-1087, OES1-1238, OES-1254, OES1-1639, OES2-0017, OES2-0052, and OES2-0065. Data represent one biological replicate and two technical replicates with individual data points represented on the graph and were fit with a nonlinear dose-response inhibition model in GraphPad Prism 10. Where applicable, $IC_{50}$ values are reported for the condition without spermine (black) and with spermine (blue).

A

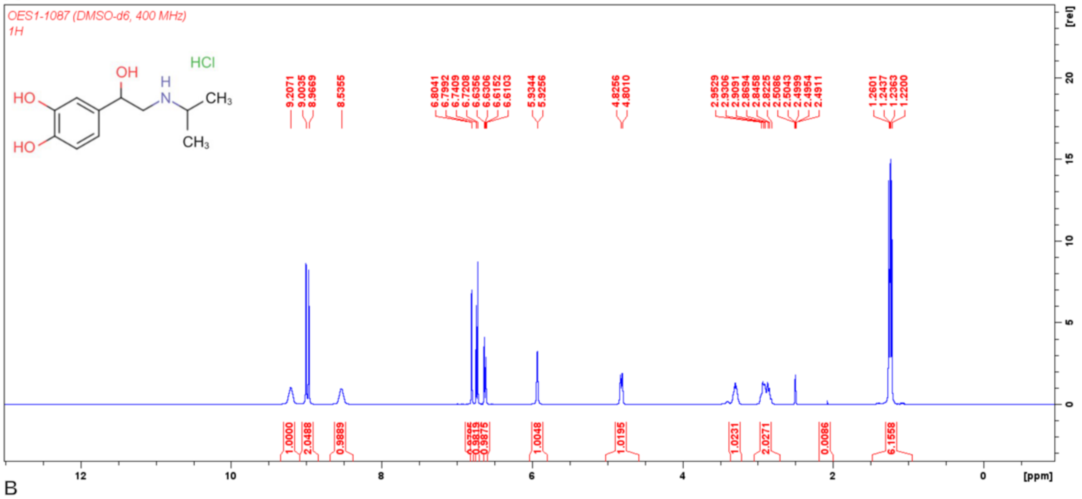
OES1-1087 (DMSO-d6, 400 MHz)
1H
HCl
[ppm]
[rel]

B

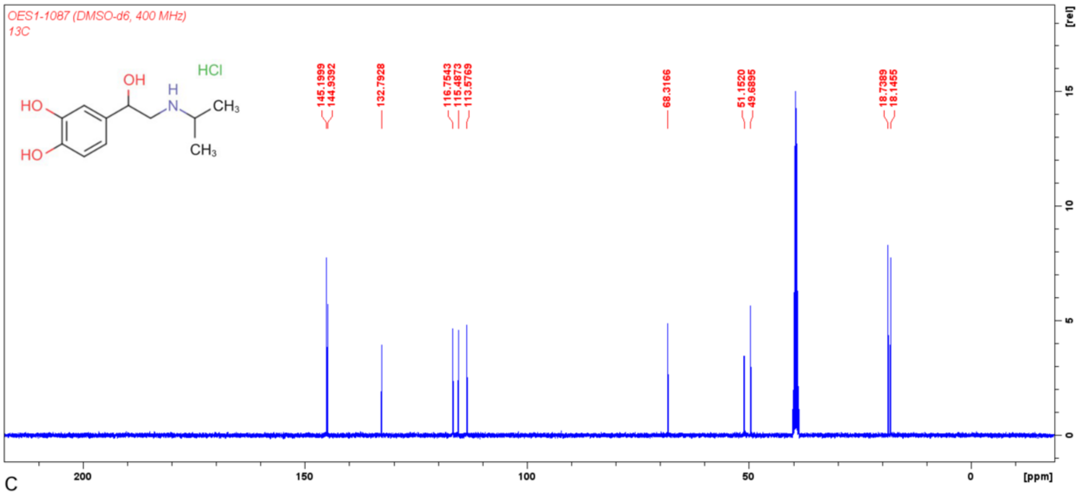
OES1-1087 (DMSO-d6, 400 MHz)
13C
HCl
[ppm]
[rel]

C

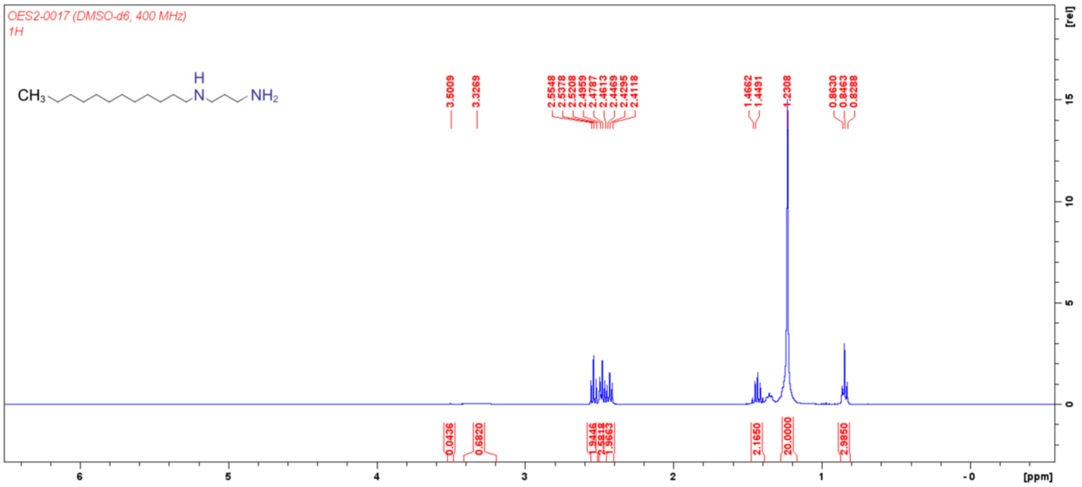
OES2-0017 (DMSO-d6, 400 MHz)
1H
[ppm]
[rel]

D

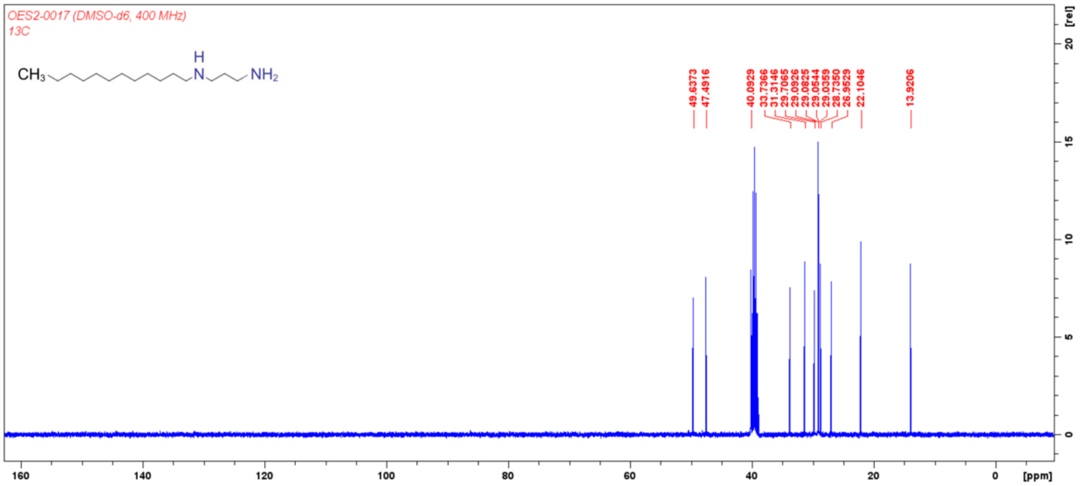


Supplementary Figure 2. **Structural confirmation of lead compounds. A)** $^{1}H$ NMR of OES1-1087 (400 MHz, DMSO-$d_6$): δ 9.21 (s, 1H), δ 8.98 (d, *J* = 14.7 Hz, 2H), 8.54 (s, 1H), 6.80 (d, *J* = 2.0 Hz, 1H), 6.73 (d, *J* = 8.0 Hz, 1H), 6.62 (dd, *J* = 8.1, 2.0 Hz, 1H), 5.93 (d, *J* = 3.5 Hz, 1H), 4.81 (d, *J* = 9.8 Hz, 1H), 2.95-2.82 (m, 2H), 1.25 (dd, *J* = 9.5, 6.5 Hz, 6H). **B)** $^{13}C$ NMR of OES1-1087 (400 MHz, DMSO-$d_6$), δ 145.20, 144.94, 132.80, 116.75, 115.49, 113.58, 68.31, 51.15, 49.69, 18.74, 18.15. **C)** $^{1}H$ NMR of OES2-0017 (400 MHz, DMSO-$d_6$) δ 3.50 (s, 1H), 3.32 (s, 2H), 2.54 (t, *J* = 6.8 Hz, 2H), 2.48 (t, *J* = 6.9 Hz, 2H), 2.43 (t, *J* = 6.9 Hz, 2H), 1.43 (quint, *J* = 6.9 Hz, 2H), 1.23 (s, 20H), 0.85 (t, *J* = 6.8 Hz, 3H). **D)** $^{13}C$ NMR of OES2-0017 (400 MHz, DMSO-$d_6$), δ 49.64, 47.49, 40.09, 33.74, 31.31, 29.71, 29.09, 29.08, 29.05, 29.04, 28.74, 26.95, 22.10, 13.92.

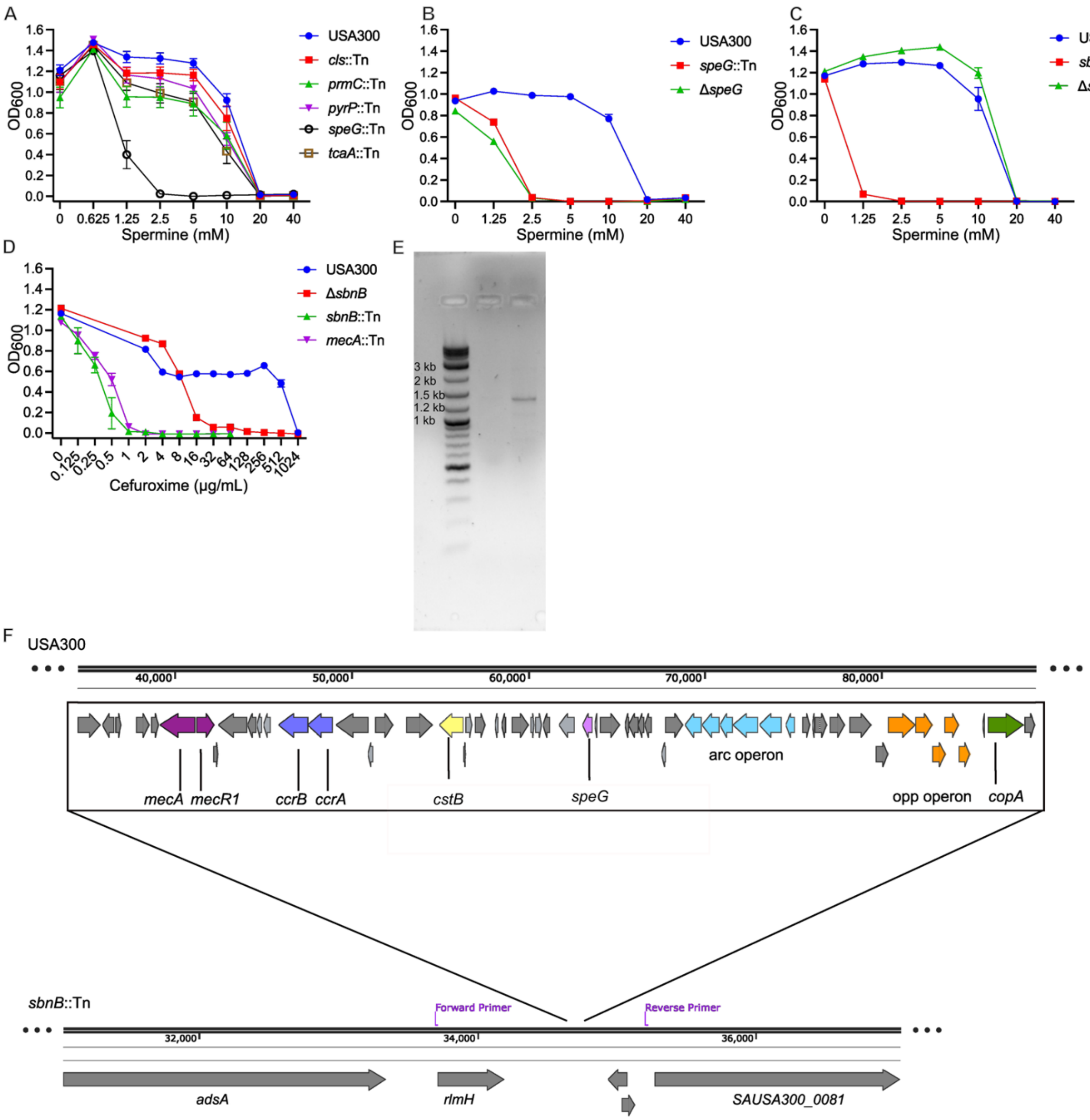


Supplementary Figure 3. **Chemogenomic screen of the NTML with spermine reveals five determinants involved in the spermine resistance of USA300.** Spermine MICs of **A)** the five transposon mutants identified in a chemogenomic screen (n=6) **B)** Δ*speG* and *speG*::Tn mutants (n=6), and of **C)** Δ*sbnB* (n=7) and sbnB::Tn (n=3) mutants**. D)** Cefuroxime MIC of *S. aureus* USA300, Δ*sbnB, sbnB*::Tn, and *mecA*::Tn (n=2). **E)** PCR of *S. aureus* USA300 (lane 2) and *sbnB*::Tn (lane 3) amplifying the region of *sbnB*::Tn that is truncated (~1500 bp) and remains intact in USA300 with the primer pair shown in (**F**). A ~46.8 kb truncation in the *sbnB*::Tn mutant includes *mecA* and *speG* rendering the strain susceptible to cefuroxime and spermine, respectively. **F)** Genetic organization depicting USA300 and the *sbnB*::Tn strain that harbours a ~46.8 kb truncation including SCC*mec* IVa cassette (including *mecA, mecR1, ccrB, ccrA,* and *cstB)* and the arginine catabolic mobile element (including *speG,* the arc and opp operons, and *copA*).

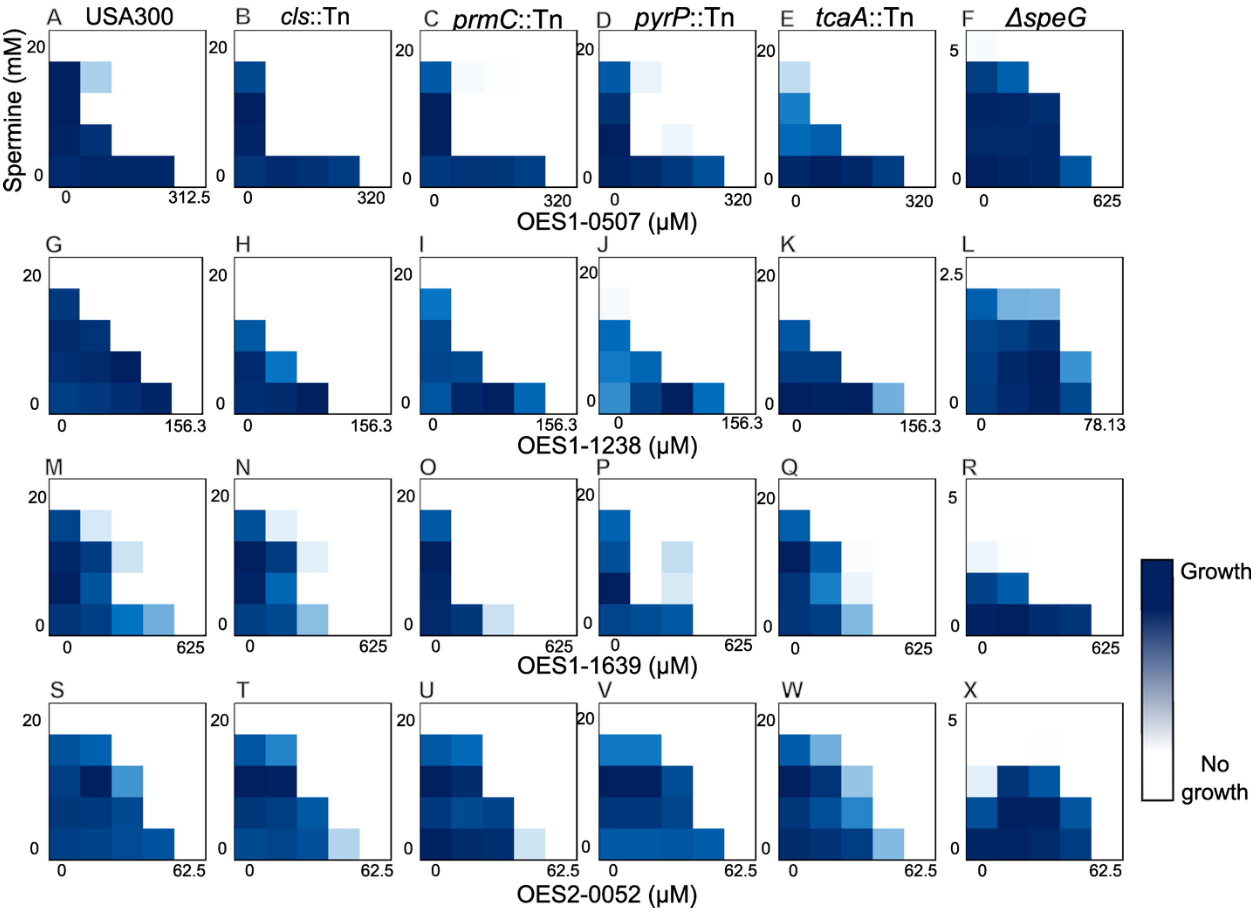


Supplementary Figure 4. **Mini-checkerboard assays of genetic determinants against hit compounds**. Representative mini-checkerboard assays of hit compounds from a chemical screen against spermine resistance determinants uncovered in a chemogenomic screen against the NTML. Checkerboards depict interactions between spermine and **A-F)** OES1-0507 and **G-L)** OES1-1238**, M-R**) OES1-1639, and **S-X)** OES2-0052 against *S. aureus* USA300 wild-type (**A, G, M, S**), *cls*::Tn **(B, H, N, T)**, *prmC*::Tn (**C, I, O, U)**, *pyrP*::Tn (**D, J, P, V)**, *tcaA*::Tn (**E, K, Q, W)**, and Δ*speG* (**F, L, R, X)**.

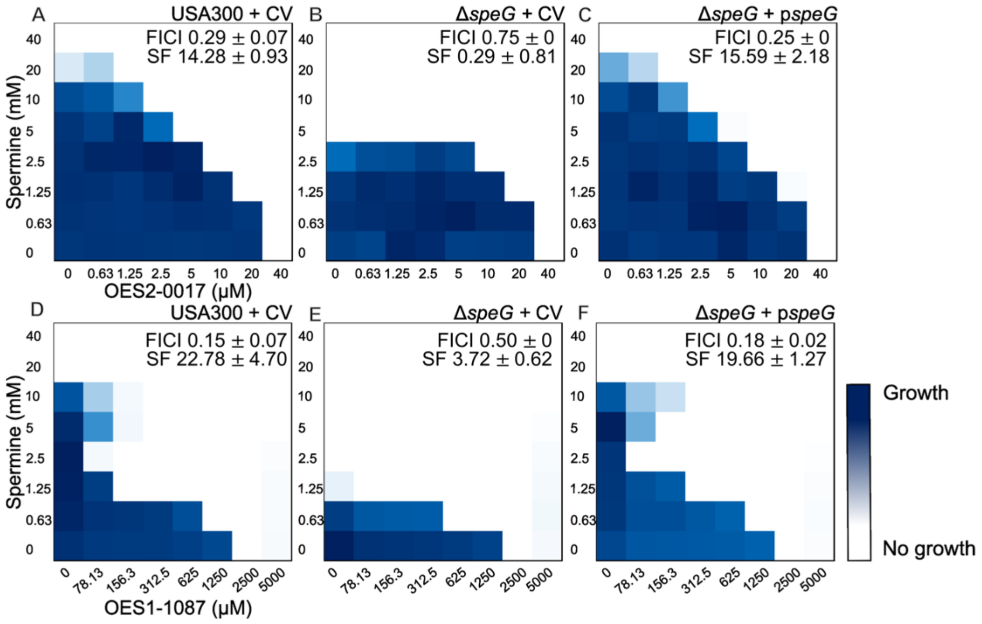


Supplementary Figure 5. **The spermine sensitivity and loss of synergy with OES2-0017 and OES1-1087 of Δ*speG* is restored by *speG* complementation.** Representative checkerboard assays of spermine and OES2-0017 or OES1-1087 against **A** and **D)** USA300 with the control vector (pKK22), **B** and **E)** Δ*speG* with the control vector, and **C** and **F)** Δ*speG* with *speG* cloned on pKK22 under the expression of the *fba* promoter. FICI values and Synergy Finder (SF) scores are reported as mean ± standard deviation of three replicates, except for **D)** n=6.

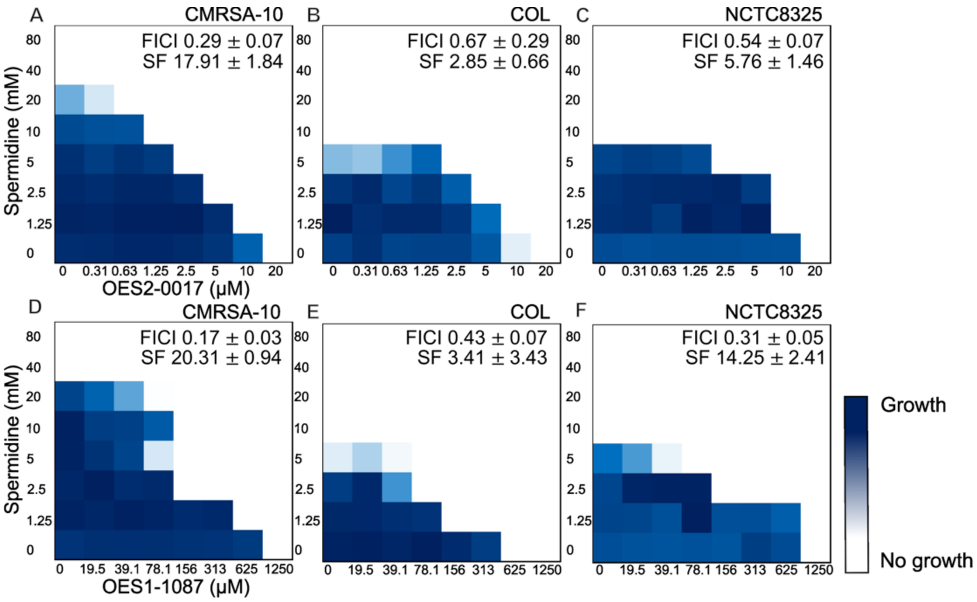


Supplementary Figure 6. **A synergistic phenotype between spermidine and OES2-0017 or OES1-1087 is reduced in *S. aureus* strains not encoding *speG.*** Representative checkerboard assays of spermidine and OES2-0017 against *S. aureus* **A)** CMRSA-10 (n=3), **B)** COL (n=3), and **C)** NCTC8325 (n=3) and of spermidine and OES1-1087 against *S. aureus* **D)** CMRSA-10 (n=3), **E)** COL (n=3), and **F)** NCTC8325 (n=3). CMRSA-10, which encodes *speG,* exhibit synergy, while the strains that do not encode *speG* (COL and NCTC8325) exhibit a reduction in the synergistic phenotype. FICI values and Synergy Finder (SF) scores are reported as mean ± standard deviation of the indicated number of replicates.

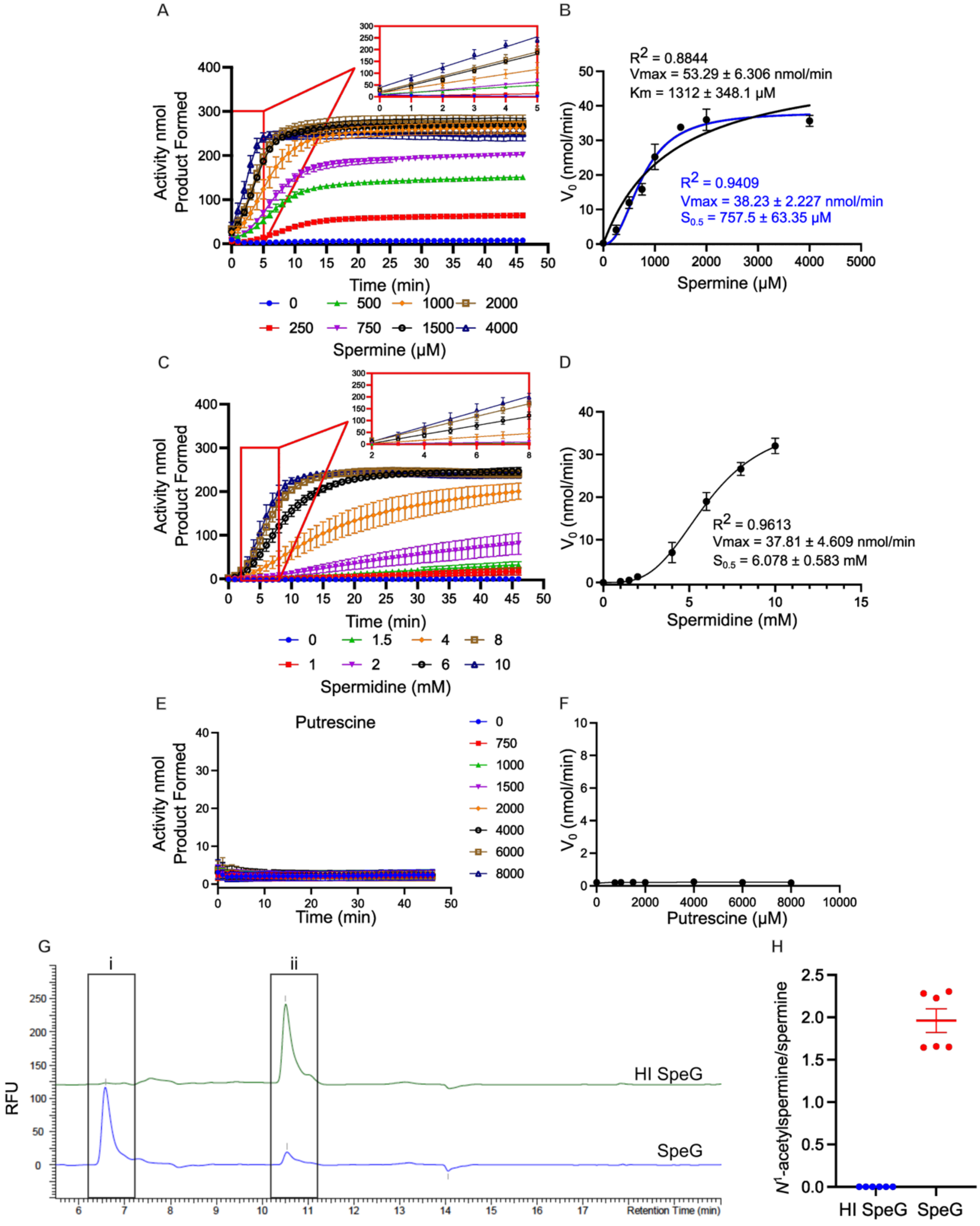


Supplementary Figure 7. ***N*¹-acetyltransferase activity of SpeG with spermine, spermidine, and putrescine. A)** Colorimetric SSAT time course with purified SpeG and spermine. **B)** Spermine substrate saturation curve generated by calculating initial velocity from the linear portion of **A)**, data were fit with both the Michaelis Menten (black) and allosteric sigmoidal (blue) equation in GraphPad Prism 10 with corresponding kinetic parameters displayed on the graph. **C)** Colorimetric SSAT time course with purified SpeG and spermidine. **D)** Spermidine substrate saturation curve generated by calculating initial velocity from the linear portion of **C)**, data was fit with an allosteric sigmoidal equation in GraphPad Prism 10. **E)** Colorimetric SSAT time course with purified SpeG and putrescine. **F)** Putrescine substrate saturation curve generated by calculating initial velocity from **E)** and fit with Michaelis-Menten equation in GraphPad Prism 10. Time course data is presented as background subtracted by a heat-inactivated enzyme control. **G)** Representative chromatogram of a 10-minute reaction of heat inactivated SpeG and SpeG with 1 mM spermine and acetyl-CoA. Primary amines were derivatized with OPA/NAC and detected by fluorescence, represented in relative fluorescence units (ex 340nm/em 450nm). i $N^1$-acetylspermine and ii spermine. **H)** Ratio of quantified $N^1$-acetylspermine/spermine by HPLC in **G)**. Results are grouped from three independent experiments **(A and B,** n=3**)** and two independent experiments (**C**-**F**, n=4; **H,** n=6) and reported as mean $\pm$ standard error of the mean**.**

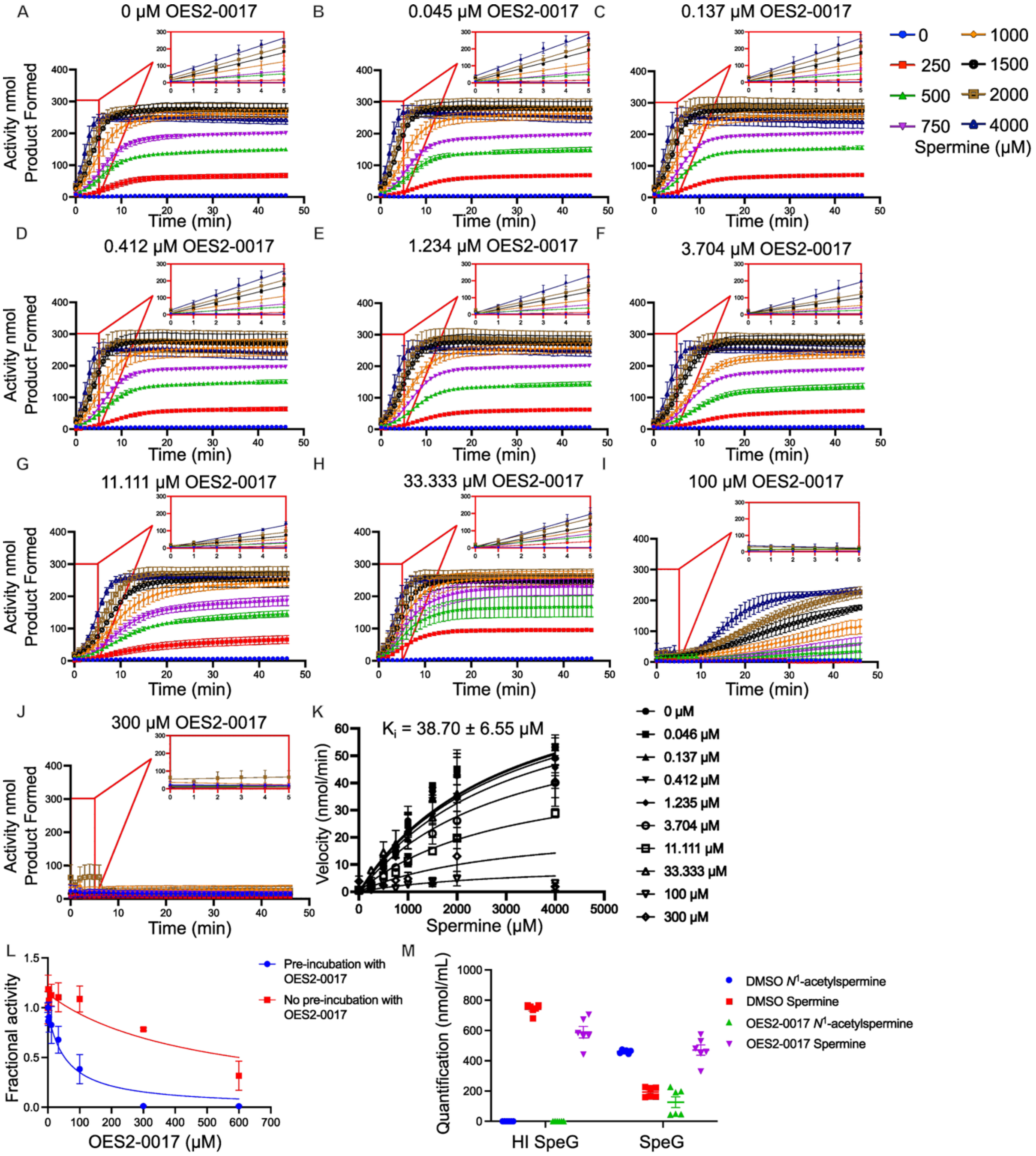


Supplementary Figure 8. **OES2-0017 activity as a SpeG inhibitor. A-J)** Colorimetric enzymatic assay of SpeG activity with increasing spermine concentrations in the presence of OES2-0017 at **A)** 0 µM, **B)** 0.045 µM, **C)** 0.137 µM, **D)** 0.412 µM, **E)** 1.234 µM, **F)** 3.704 µM, **G)** 11.11 µM, **H)** 33.333 µM, **I)** 100 µM, and **J)** 300 µM. **K)** Substrate saturation curves of SpeG with a fixed concentration of acetyl-CoA and variable spermine concentrations in the presence of OES2-0017. Data was fit with the noncompetitive inhibitor equation in GraphPad Prism 10. **L)** $IC_{50}$ plot of a colorimetric SSAT enzymatic reaction of SpeG inhibition by OES2-0017 where SpeG was pre-incubated with OES2-0017 before adding spermine and acetyl CoA as substrates (blue line) or without pre-incubation with OES2-0017 and spermine, acetyl CoA, and OES2-0017 added to the reaction at the same time. Data is represented as mean ± SEM for two independent experiments n=4. **M)** Quantification of $N^1$-acetylspermine and spermine by HPLC analysis of a 10-minute reaction of SpeG and heat inactivated SpeG with 1 mM spermine, 1 mM Acetyl-CoA and 300 μM OES2-0017. Primary amines were derivatized with OPA/NAC and detected by fluorescence, represented in relative fluorescence units (ex 340nm/em 450nm). Data are grouped from three independent experiments **(A-K,** n=3**)** and two independent experiments (**L**, n=4; **M**, n=6) and reported as mean ± standard error of the mean**.**

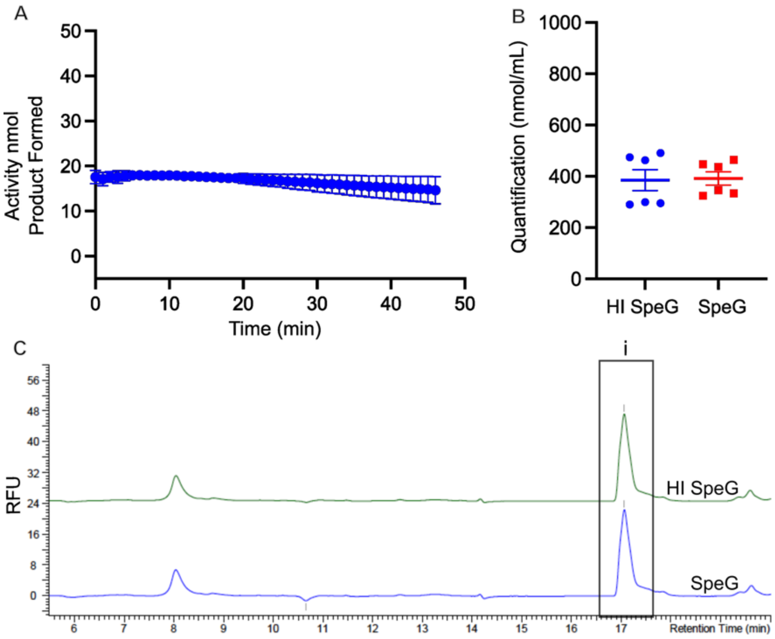


Supplementary Figure 9. **SpeG exhibits no *N*[1]-acetyltransferase activity towards the polyamine analog OES2-0017. A)** Colorimetric SSAT assay of SpeG with 1 mM Acetyl-CoA and 300 μM OES2-0017 detecting the amount of Coenzyme A liberated during the acetyl-transfer from Acetyl-CoA. **B)** Quantification of OES2-0017 by HPLC analysis of a 10-minute reaction of SpeG and heat inactivated SpeG with 1 mM Acetyl-CoA and OES2-0017. Primary amines were derivatized with OPA/NAC and detected by fluorescence, represented in relative fluorescence units (ex 340nm/em 450nm). **C)** Representative chromatograms quantifying (i) OES2-0017 of heat inactivated SpeG and SpeG. Results are grouped from two independent experiments **A**) n=2 **B)** n=6 and reported as mean ± SEM.

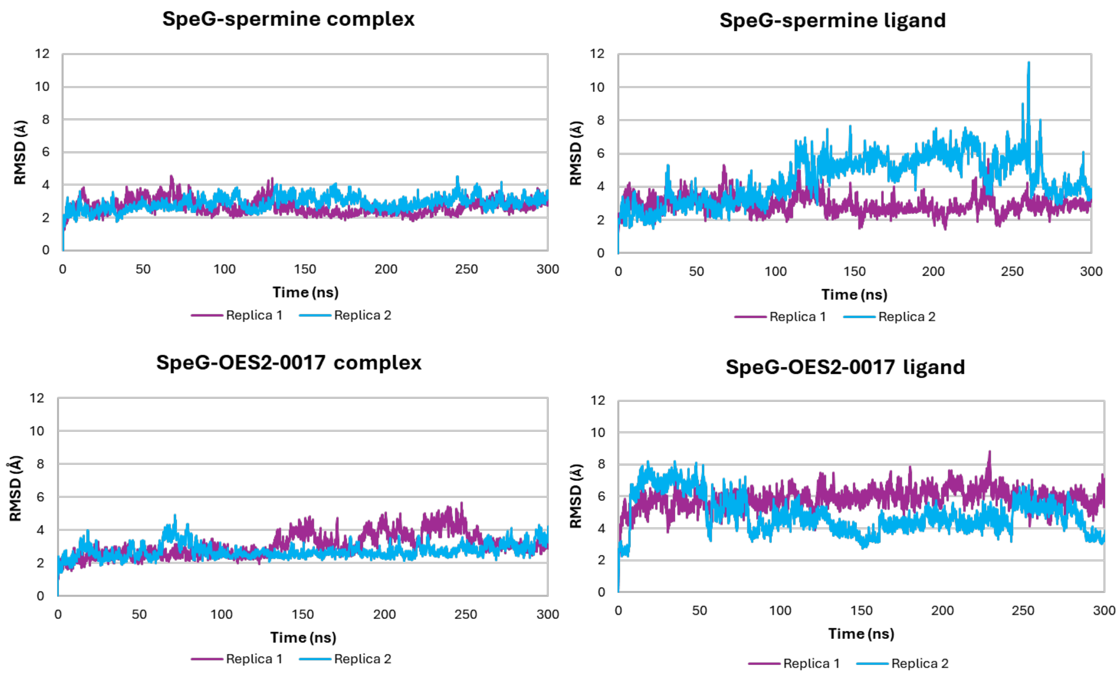


Supplementary Figure 10. Root Mean Square Deviation (RMSD) of the SpeG dimer (PDB ID 8fv1) in complex with spermine (top) and OES2-0017 (bottom). RMSD has been calculated for the whole complex (left) and only for the ligands (right).

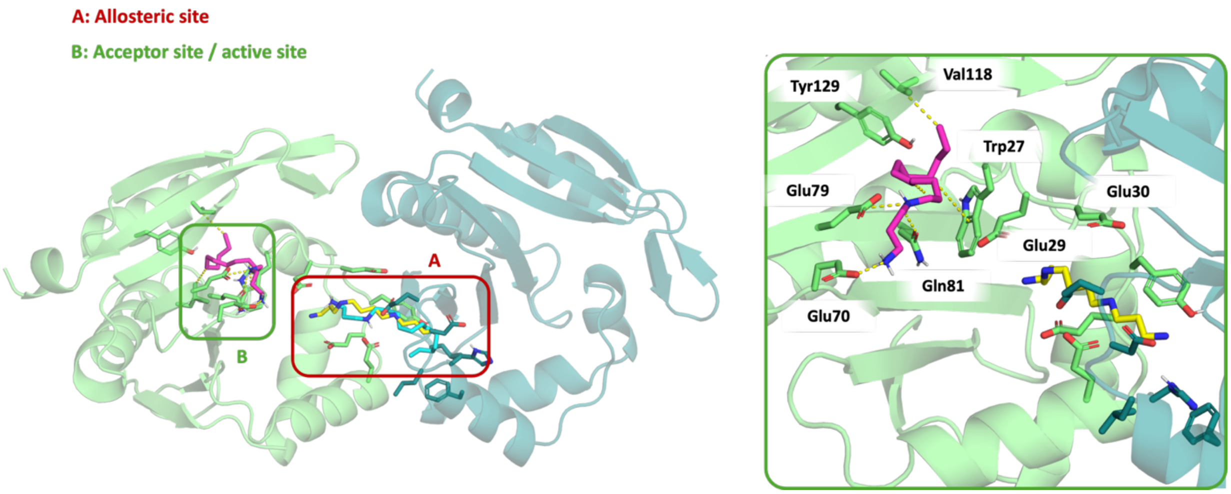


Supplementary Figure 11. **Docking of OES2-0017 into the allosteric and acceptor sites of SpeG. (left)** General view of the two binding sites studied by docking for compound OES2-0017. The allosteric site is marked with a red box (A) and acceptor site with a green box (B). Crystallographic spermine (8fv1) is depicted as yellow sticks, the predicted docked pose of OES2-0017 in the allosteric site is depicted as cyan sticks and the predicted docked pose of OES2-0017 in the acceptor site is depicted as magenta sticks. **(Right)** Detail of the docked binding mode of OES2-0017 in the acceptor site. Non-polar hydrogens are hidden for clarity.

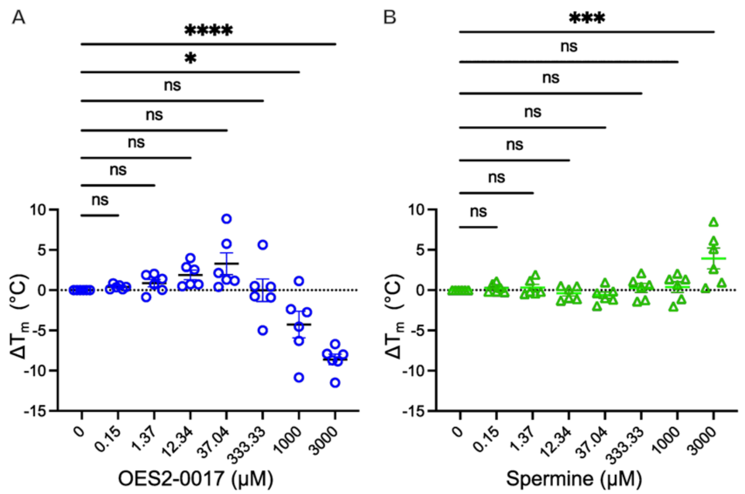


Supplementary Figure 12. **OES2-0017 and spermine bind SpeG.** Thermal shift assay measuring melting temperature ($\Delta T_m$) in response to **A)** OES2-0017 or **B)** spermine treatment. Results are grouped from six independent experiments represented as mean ± SEM, n=6. Statistical significance was determined by ordinary one-way ANOVA with Dunnett's multiple comparisons test. * represents p ≤ 0.05, ** p ≤ 0.01, *** p ≤ 0.001, **** p ≤ 0.0001, and ns p > 0.05.

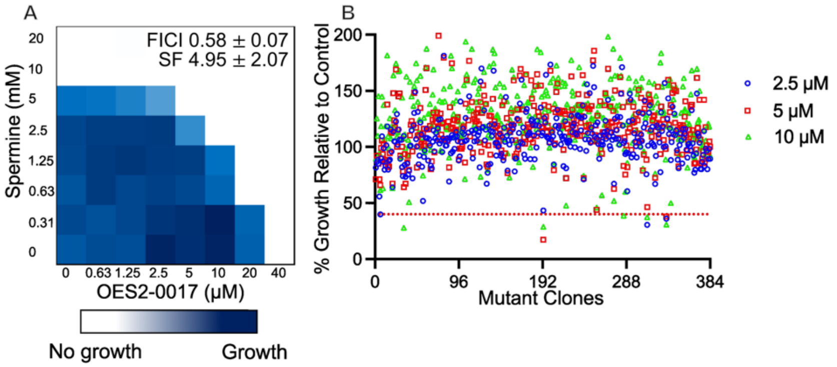


Supplementary Figure 13. **Essential genomic screen did not uncover essential protein targets for OES2-0017. A)** Representative checkerboard assay of Spermine and OES2-0017 against *B. subtilis* 168 (n=3). **B)** Screen of OES2-0017 at 1/4th (blue circles), 1/8th (red squares), and 1/16th (green triangles) the growth inhibitory concentration against the *B. subtilis* knockdown collection of essential genes with 0.05% xylose induction.

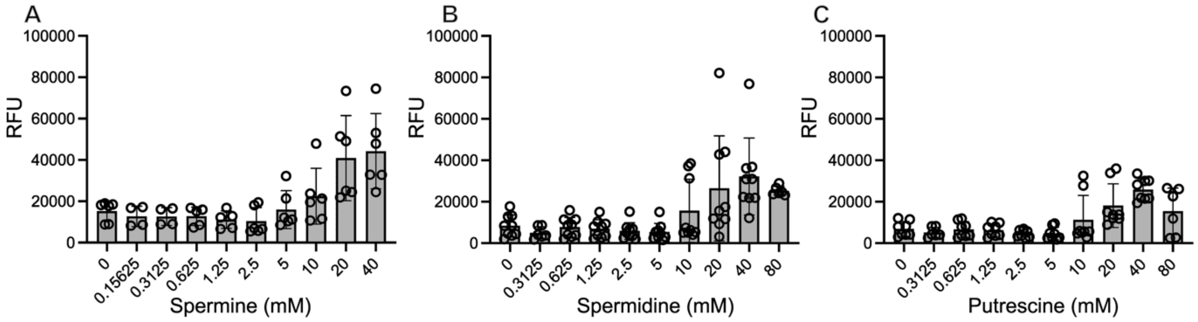


Supplementary Figure 14. **Polyamines disrupt membrane integrity of *S. aureus*.** DiSC3(5) assays against *S. aureus* USA300 with **A)** spermine, **B)** spermidine, and **C)** putrescine. Increases in fluorescent signal of DiSC3(5) dye suggests disruption in membrane integrity. Individual data points are represented from 4 (spermine n= 6, and spermidine n=9) or 3 (putrescine, n=8) independent experiments.

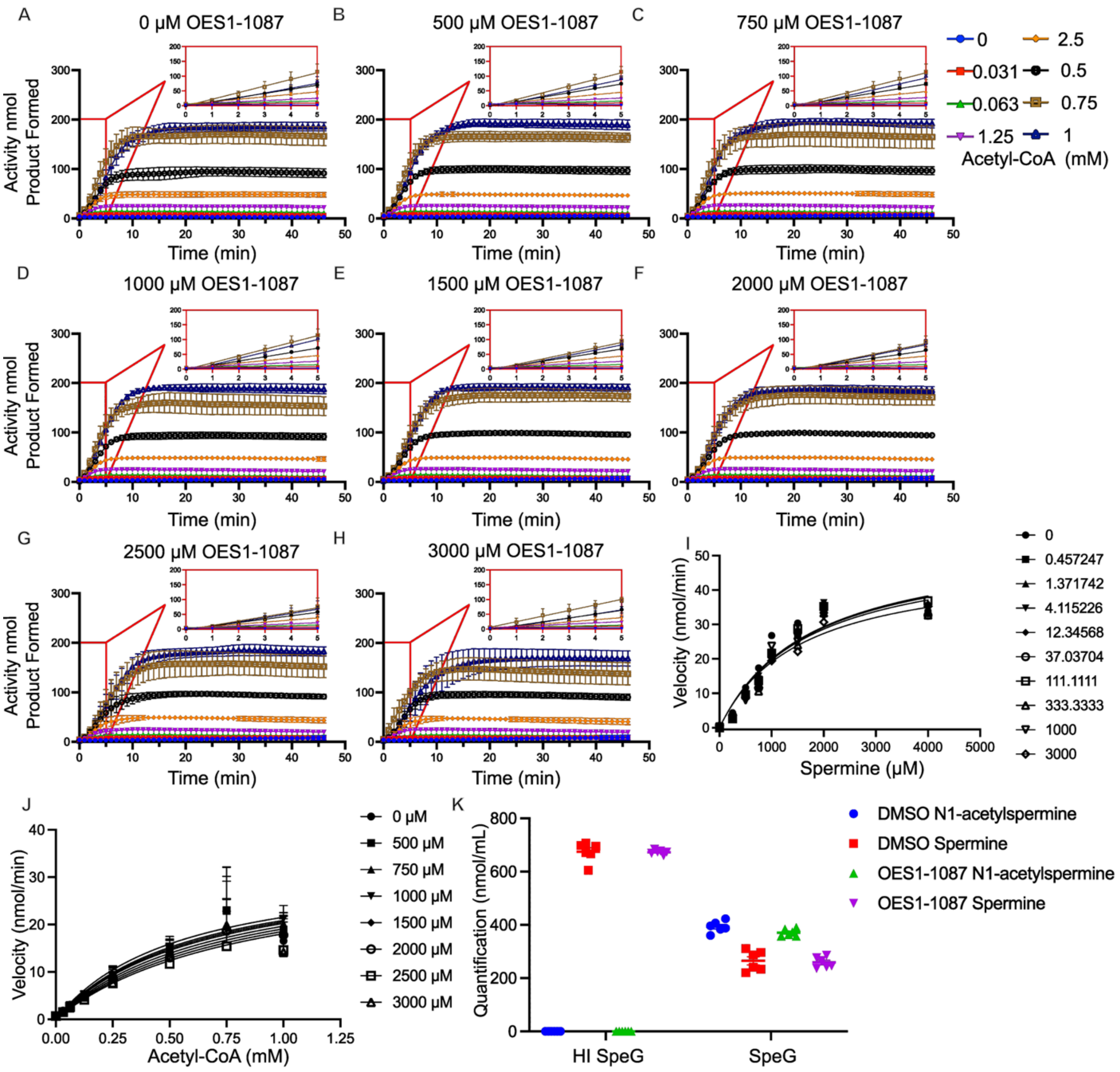

Supplementary Figure 15. **OES1-1087 activity as a SpeG inhibitor. A-H)** Colorimetric enzymatic assay of SpeG activity with increasing acetyl-CoA concentrations in the presence of OES1-1087 at **A)** 0 μM, **B)** 500 μM, **C)** 750 μM, **D)** 1000 μM, **E)** 1500 μM, **F)** 2000 μM, **G)** 2500 μM, and **H)** 3000 μM. **I)** Substrate saturation curve of SpeG with a fixed concentration of acetyl-CoA and variable spermine concentrations in the presence of OES1-1087. **J)** Substrate saturation curve of SpeG with a fixed concentration of spermine and variable acetyl-CoA concentrations in the presence of OES1-1087. Data was fit with the competitive inhibitor equation in GraphPad Prism 10. **K)** Quantification of $N^1$-acetylspermine and spermine by HPLC analysis of a 10-minute reaction of SpeG and heat inactivated SpeG pre-incubated with 3 mM OES1-1087 for 30 min before initiating the reaction with 1 mM spermine and 0.75 mM Acetyl-CoA. Primary amines were derivatized with OPA/NAC and detected by fluorescence, represented in relative fluorescence units (ex 340nm/em 450nm). Data is represented as mean ± SEM for three independent experiments n=3, except for **K)** which represents two independent experiments n=6, and **I)** where n=1.

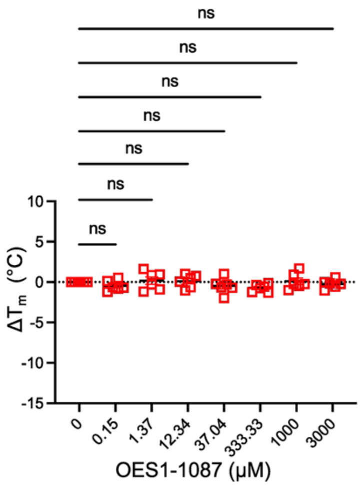


Supplementary Figure 16. **OES1-1087 does not alter the $T_m$ of SpeG.** Thermal shift assay measuring melting temperature ($\Delta T_m$) in response to OES1-1087 treatment. Results are grouped from six independent experiments represented as mean ± SEM, n=6. Statistical significance was determined by ordinary one-way ANOVA with Dunnett's multiple comparisons test. * represents $p \leq 0.05$, ** $p \leq 0.01$, *** $p \leq 0.001$, **** $p \leq 0.0001$, and ns $p > 0.05$.

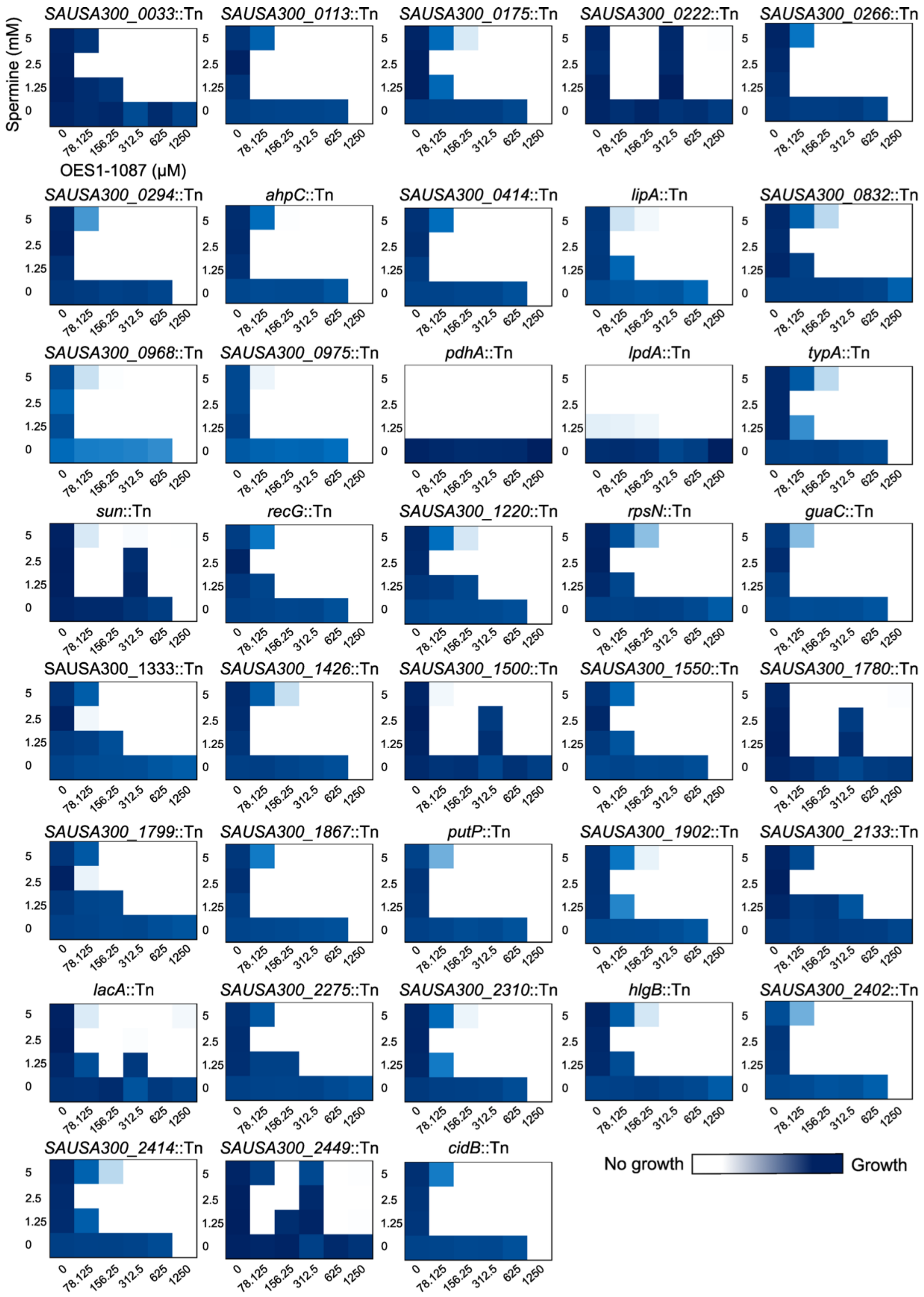


Supplementary Figure 17. **Mini-checkerboard assays of genetic determinants identified in a OES1-1087 screen**. Representative mini-checkerboard assays of OES1-1087 and spermine against determinants identified in a 312.5 and 625 μM OES1-1087 chemogenomic screen against the NTML.

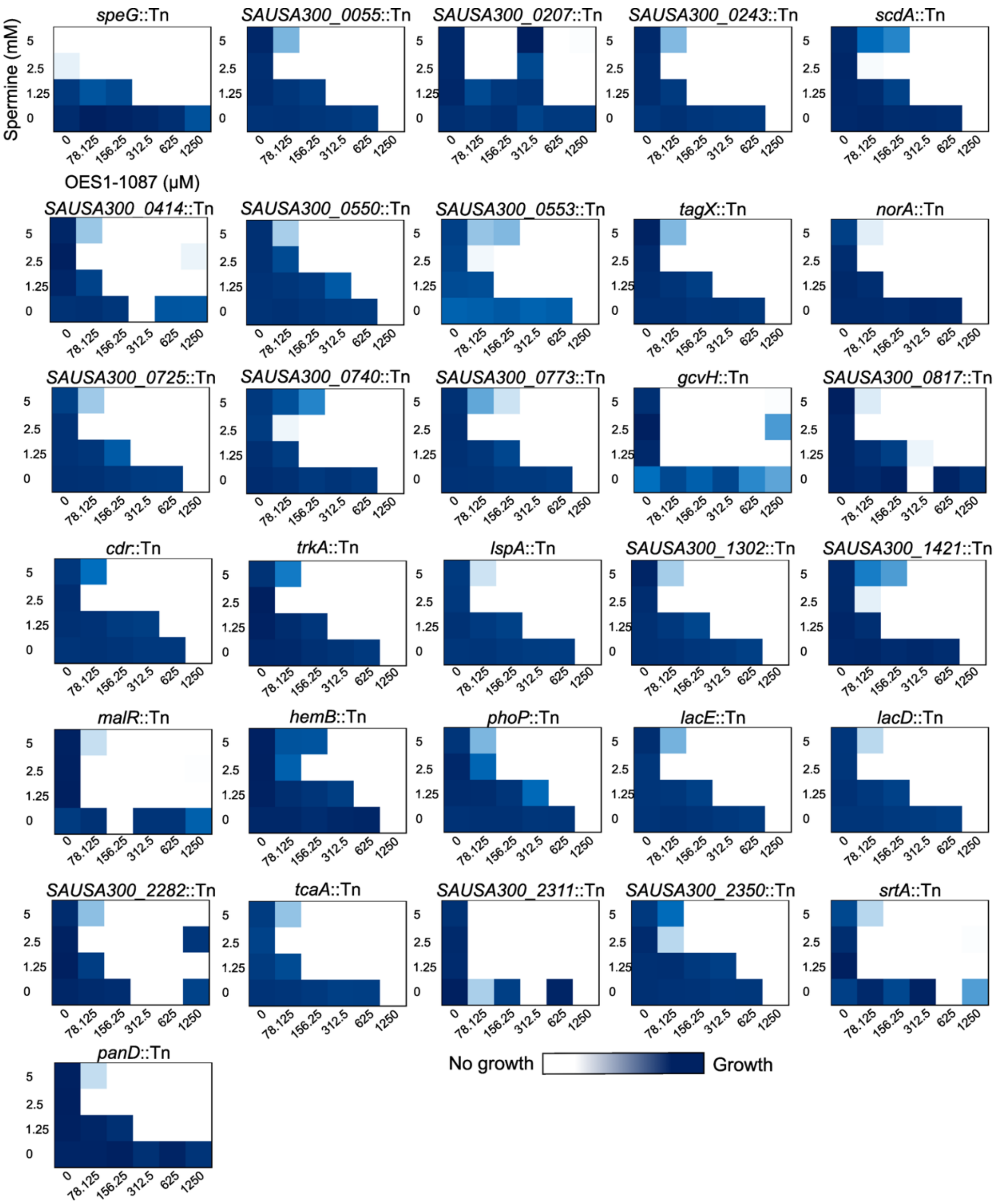


Supplementary Figure 18. **Mini-checkerboard assays of genetic determinants identified in a spermine screen**. Representative mini-checkerboard assays of OES1-1087 and spermine against determinants identified in a 5 mM spermine chemogenomic screen against the NTML.

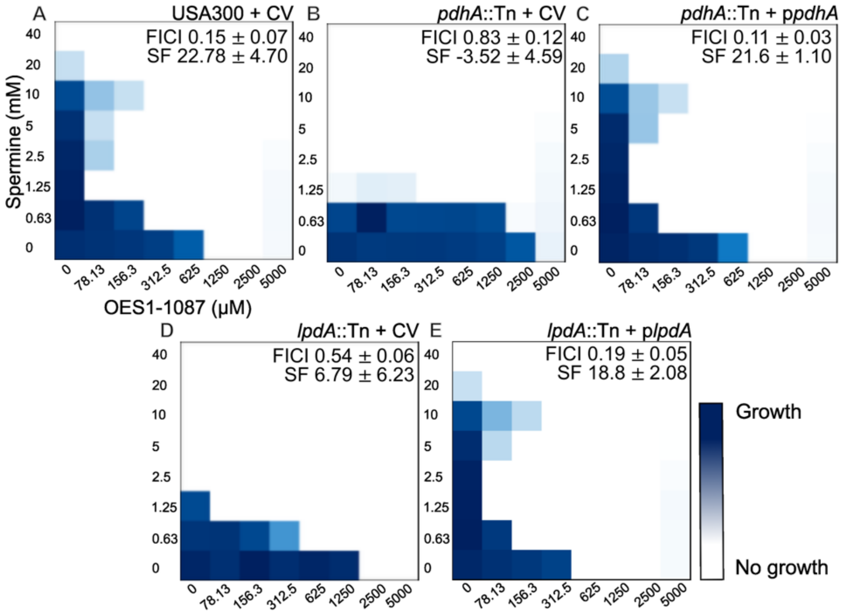


Supplementary Figure 19. **The spermine sensitivity and loss of synergy with OES1-1087 of *pdhA*::Tn and *lpdA*::Tn are restored by *pdhA* and *lpdA* complementations.** Representative checkerboard assays of spermine and OES1-1087 against **A)** USA300 with the control vector (pKK22) **B)** *pdhA*::Tn with the control vector, **C)** *pdhA*::Tn with *pdhA* cloned on pKK22 under the expression of the *fba* promoter, **D)** *lpdA*::Tn with the control vector, **E)** *lpdA*::Tn with *lpdA* cloned on pKK22 under the expression of the *fba* promoter. FICI values and Synergy Finder (SF) scores are reported as mean ± standard deviation of three replicates, except for **A)** n=6.

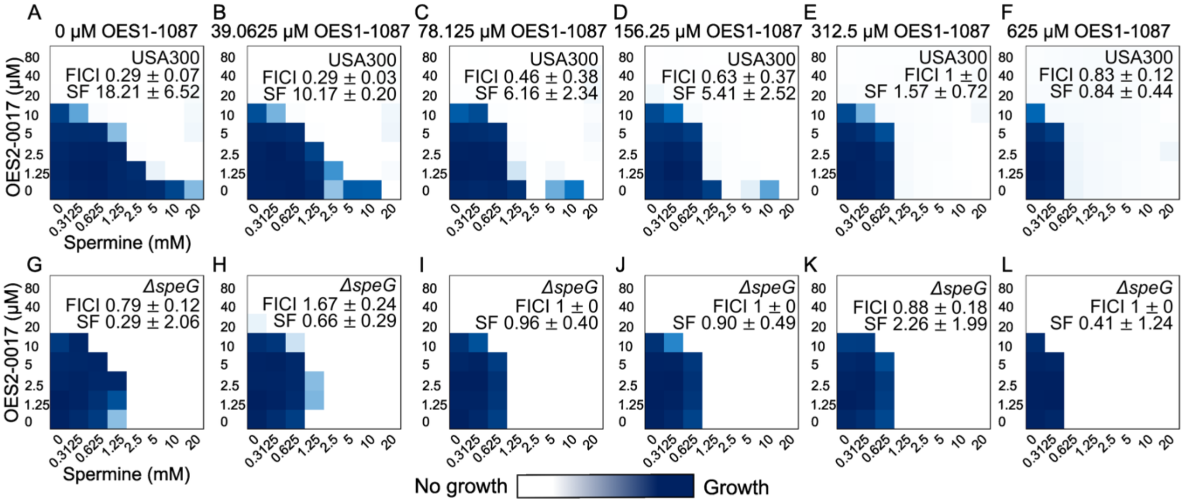


Supplementary Figure 20. **OES2-0017 and OES1-1087 do not synergize with each other in the presence of spermine suggesting they act in the same pathway.** Representative checkerboard assays of OES2-0017 and spermine with OES1-1087 added to media against *S. aureus* **A-F)** USA300 and **G-L)** Δ*speG*. FICI and Synergy Finder (SF) scores are reported as mean ± standard deviation**,** n=3.

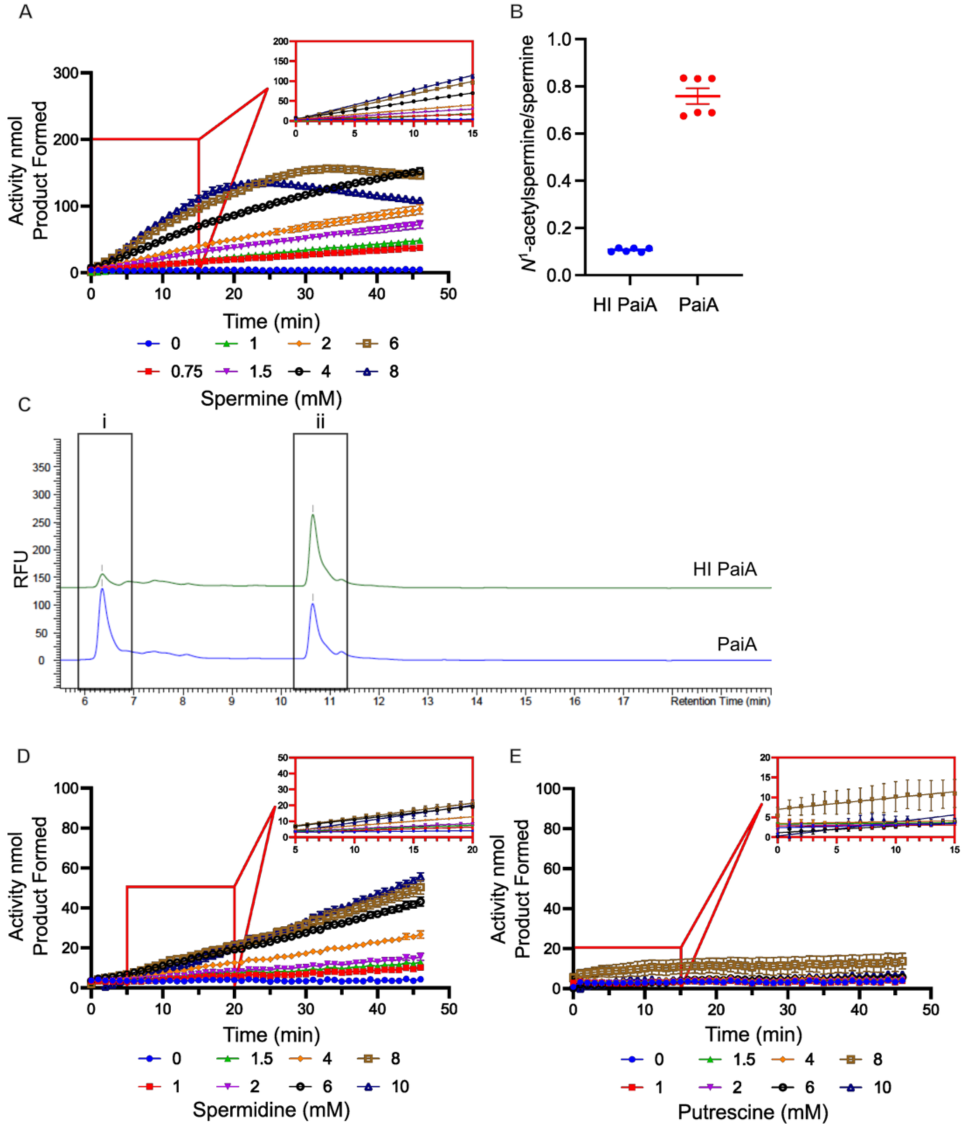


Supplementary Figure 21. **Polyamine acetyltransferase activity of *S. aureus* PaiA$_{Sa}$. A)** Colorimetric enzymatic assay of PaiA$_{Sa}$ with spermine measuring the production of Coenzyme A over time. **B)** Ratio of quantified $N^1$-acetylspermine and spermine by HPLC analysis of a 1-hour reaction of PaiA$_{Sa}$ and heat inactivated PaiA$_{Sa}$ with 1 mM spermine and 1 mM Acetyl-CoA. Primary amines were derivatized with OPA/NAC and detected by fluorescence, represented in relative fluorescence units (ex 340nm/em 450nm). **C)** Representative chromatograms of heat-inactivated PaiA$_{Sa}$ and PaiA$_{Sa}$, i $N^1$-acetylspermine and ii spermine. Colorimetric enzymatic assays of PaiA$_{Sa}$ with **D)** spermidine and **E)** putrescine as substrates measuring the production of Coenzyme A over time. Results are grouped from two independent experiments reported as mean $\pm$ SEM **A)**, **D)**, and **E)** n=4, **B)** n=6.

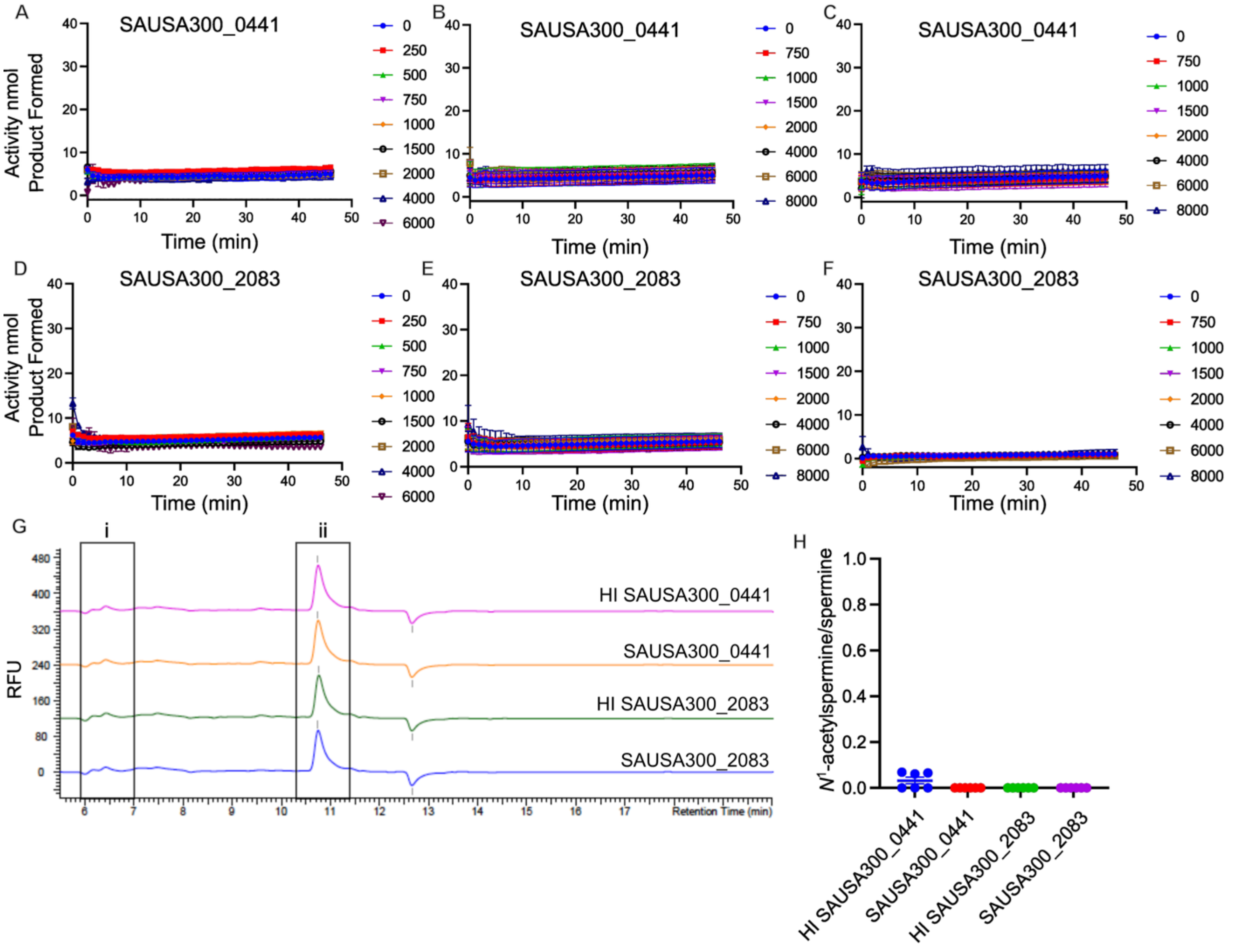


Supplementary Figure 22. **Polyamine acetyltransferase activity of BltD homologs. A-C)** Colorimetric enzymatic assays of SAUSA300_0441 with **A)** spermine, **B)** spermidine, and **C)** putrescine measuring the production of Coenzyme A over time. **D-F)** Colorimetric enzymatic assays of SAUSA300_2083 with **D)** spermine, **E)** spermidine, and **F)** putrescine. **G)** Representative chromatograms of heat-inactivated SAUSA300_0441, SAUSA300_0441, heat-inactivated SAUSA300_2083, and SAUSA300_2083 of a 1-hour reaction with 1 mM spermine and 1 mM Acetyl-CoA detecting i $N^1$-acetylspermine and ii spermine. Primary amines were derivatized with OPA/NAC and detected by fluorescence, represented in relative fluorescence units (ex 340nm/em 450nm). **H)** Ratio of quantified $N^1$-acetylspermine and spermine by HPLC analysis of a 1-hour reaction with 1 mM spermine and 1 mM Acetyl-CoA. Results are grouped from two independent experiments reported as mean ± SEM **A-F)** n=4 and **H)** n=6.

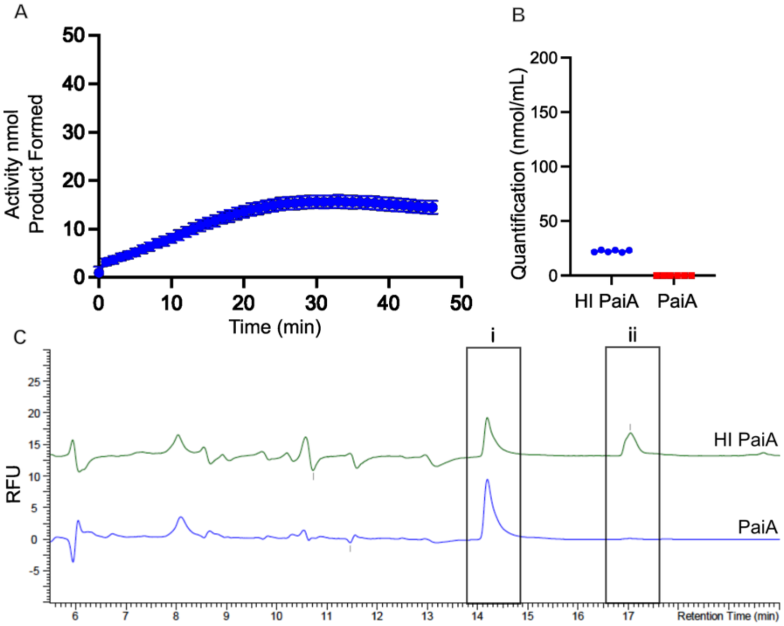


Supplementary Figure 23. **PaiA exhibits *N*[1]-acetyltransferase activity towards the polyamine analog OES2-0017. A)** Colorimetric SSAT assay of $PaiA_{Sa}$ with 1 mM Acetyl-CoA and 300 μM OES2-0017 detecting the amount of Coenzyme A liberated during the acetyl-transfer from Acetyl-CoA. **B)** Quantification of OES2-0017 by HPLC analysis of a 1-hour reaction of $PaiA_{Sa}$ and heat inactivated $PaiA_{Sa}$ with 1 mM Acetyl-CoA and OES2-0017. Primary amines were derivatized with OPA/NAC and detected by fluorescence, represented in relative fluorescence units (ex 340nm/em 450nm). **C)** Representative chromatograms of reactions containing heat-inactivated $PaiA_{Sa}$ and $PaiA_{Sa}$ with 1 mM each Acetyl-CoA and OES2-0017 revealed i, unidentified peak, hypothesized to be acetylated OES2-0017 and ii, OES2-0017. Results are grouped from two independent experiments **A**) n=4 **B)** n=6 and reported as mean ± SEM.

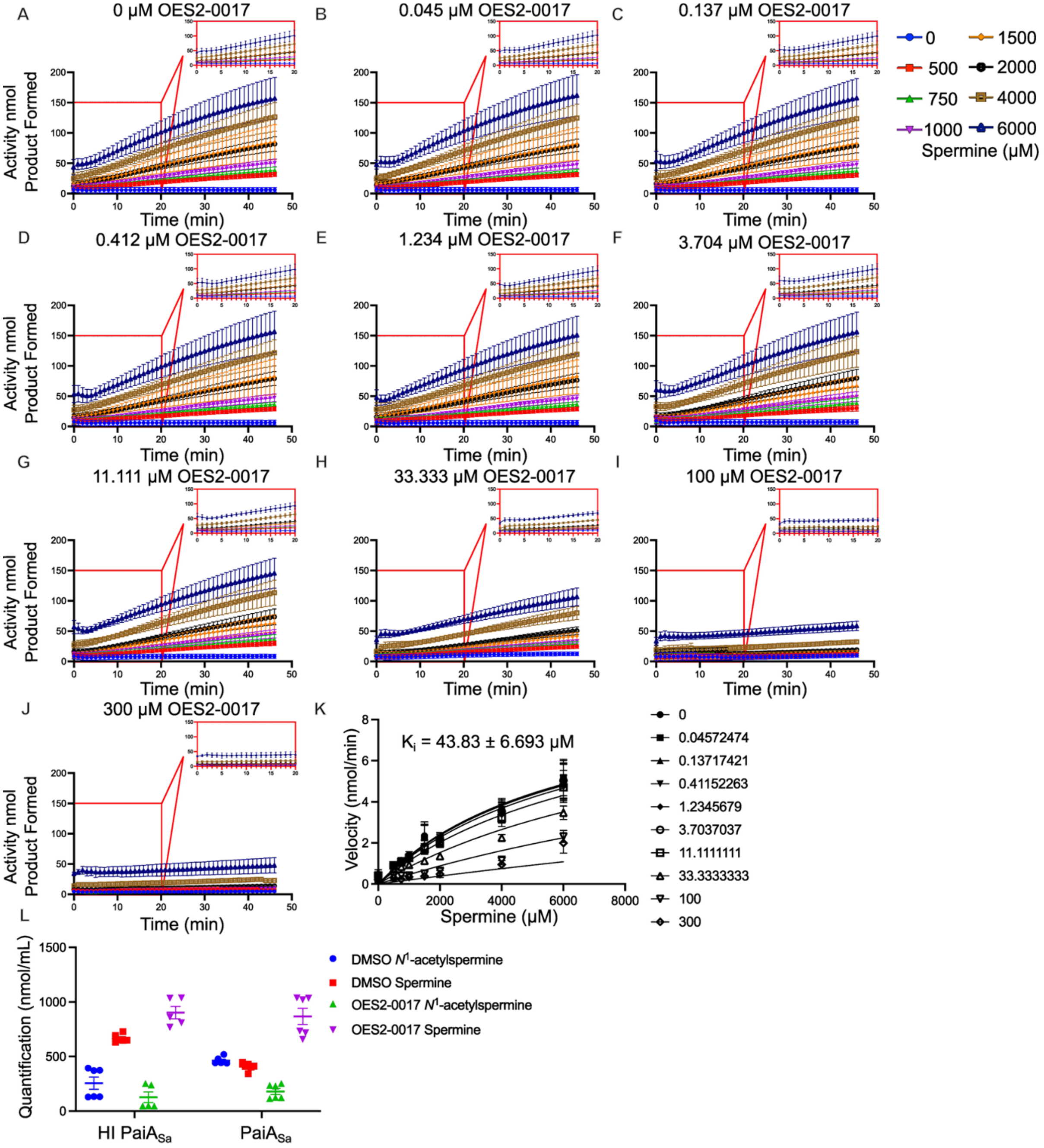


Supplementary Figure 24. **OES2-0017 activity as a PaiA$_{Sa}$ inhibitor. A-J)** Colorimetric enzymatic assay of PaiA$_{Sa}$ activity with increasing spermine concentrations in the presence of OES2-0017 at **A)** 0 μM, **B)** 0.045 μM, **C)** 0.137 μM, **D)** 0.412 μM, **E)** 1.234 μM, **F)** 3.704 μM, **G)** 11.11 μM, **H)** 33.333 μM, **I)** 100 μM, and **J)** 300 μM. **K)** Substrate saturation curves of PaiA$_{Sa}$ with a fixed concentration of acetyl-CoA and variable spermine concentrations in the presence of OES2-0017. Data was fit with the competitive inhibitor equation in GraphPad Prism 10. **L)** quantification of $N^1$-acetylspermine and spermine by HPLC of a 1-hour reaction of heat-inactivated PaiA$_{Sa}$ and PaiA$_{Sa}$ with 1 mM each acetyl-CoA and spermine with and without 1.2 mM OES2-0017. Primary amines were derivatized with OPA/NAC and detected by fluorescence, represented in relative fluorescence units (ex 340nm/em 450nm). Data is represented as mean ± SEM for three independent experiments **(A-K)** n=3, and two independent experiments **L)** n=6.

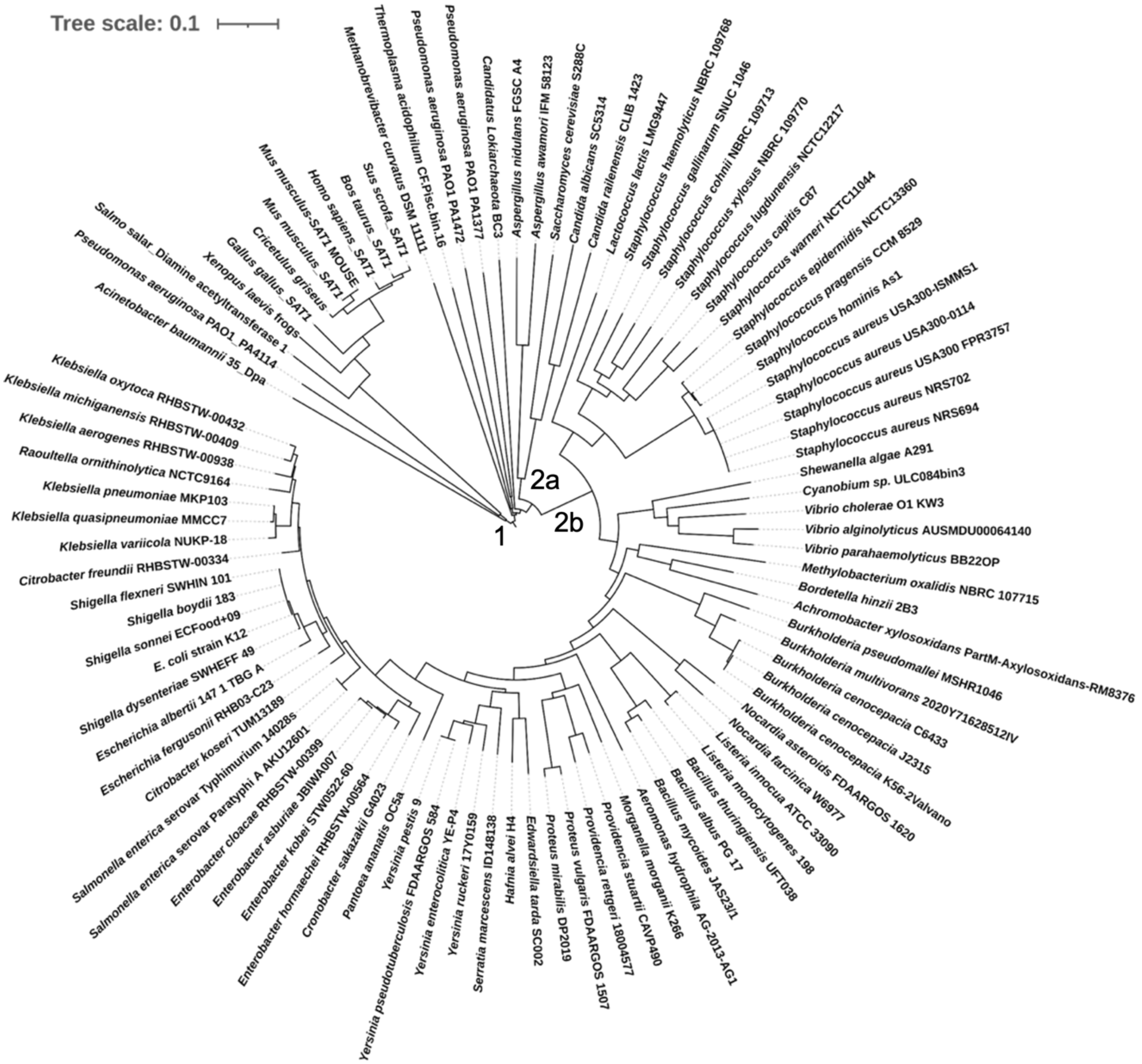


Supplementary Figure 25. **Phylogenetic analysis of polyamine acetyltransferases.** The evolutionary history was inferred using the neighbor-joining method [7]. The phylogenetic tree is drawn to scale, with branch lengths in the same units as those of the evolutionary distances used to infer the phylogenetic tree. The evolutionary distances were computed using the p-distance method [8] and are in the units of the number of amino acid differences per site. The rate variation among sites was modeled with a gamma distribution (shape parameter = 1). This analysis involved 94 amino acid sequences (Supplementary Table 4). All ambiguous positions were removed for each sequence pair (pairwise deletion option). There were a total of 384 positions in the final dataset, positions represent the amino acid sites compared across all sequences. Evolutionary analyses were conducted in MEGA11 [9].

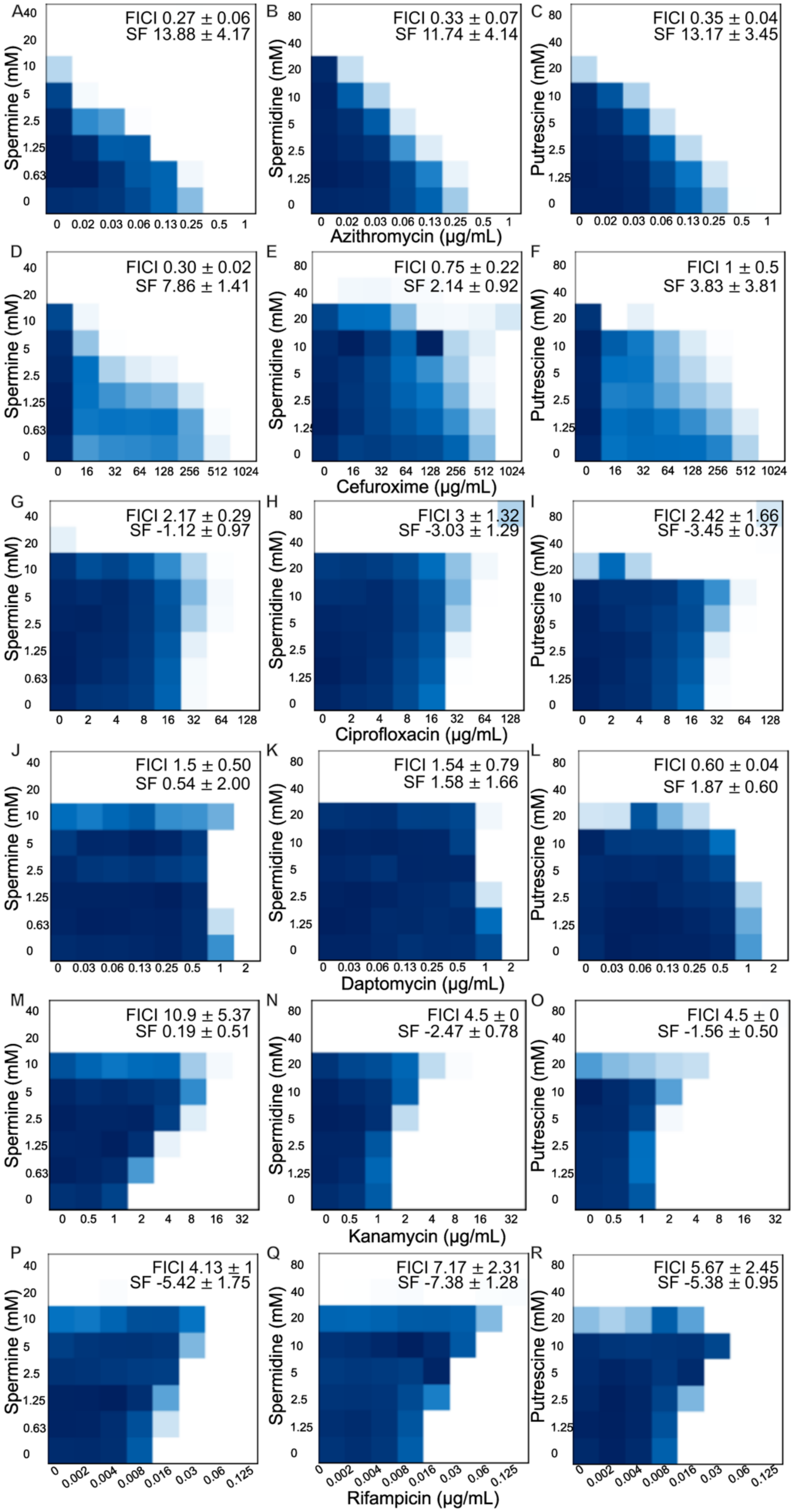
A
FICI 0.27 ± 0.06
SF 13.88 ± 4.17
B
FICI 0.33 ± 0.07
SF 11.74 ± 4.14
C
FICI 0.35 ± 0.04
SF 13.17 ± 3.45
Azithromycin (μg/mL)
D
FICI 0.30 ± 0.02
SF 7.86 ± 1.41
E
FICI 0.75 ± 0.22
SF 2.14 ± 0.92
F
FICI 1 ± 0.5
SF 3.83 ± 3.81
Cefuroxime (μg/mL)
G
FICI 2.17 ± 0.29
SF -1.12 ± 0.97
H
FICI 3 ± 1.32
SF -3.03 ± 1.29
I
FICI 2.42 ± 1.66
SF -3.45 ± 0.37
Ciprofloxacin (μg/mL)
J
FICI 1.5 ± 0.50
SF 0.54 ± 2.00
K
FICI 1.54 ± 0.79
SF 1.58 ± 1.66
L
FICI 0.60 ± 0.04
SF 1.87 ± 0.60
Daptomycin (μg/mL)
M
FICI 10.9 ± 5.37
SF 0.19 ± 0.51
N
FICI 4.5 ± 0
SF -2.47 ± 0.78
O
FICI 4.5 ± 0
SF -1.56 ± 0.50
Kanamycin (μg/mL)
P
FICI 4.13 ± 1
SF -5.42 ± 1.75
Q
FICI 7.17 ± 2.31
SF -7.38 ± 1.28
R
FICI 5.67 ± 2.45
SF -5.38 ± 0.95
Rifampicin (μg/mL)
Spermine (mM)
Spermidine (mM)
Putrescine (mM)

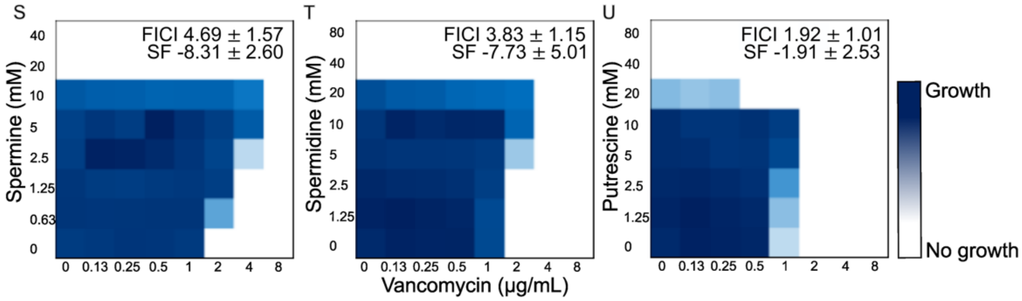


Supplementary Figure 26. **Polyamines influence antibiotic susceptibility of *S. aureus* USA300.** Representative checkerboard assays with spermine (column 1), spermidine (column 2), putrescine (column 3), and representative antibiotics against *S. aureus* USA300 **A-C)** azithromycin, **D-F)** cefuroxime, **G-I)** ciprofloxacin, **J-L)** daptomycin, **M-O)** kanamycin, **P-R)** rifampicin, and **S-U)** vancomycin. FICI values and Synergy Finder (SF) scores are reported as mean ± standard deviation for three independent experiments with the exceptions of **A)** n=6, **M)** n=5, **P)** n=8, and **S)** n=10.

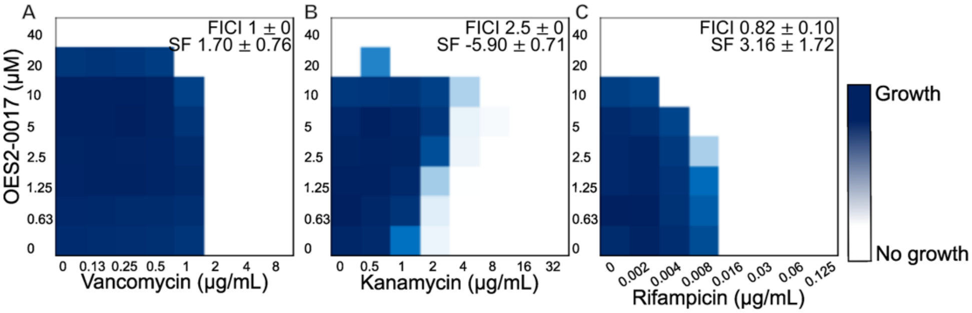


Supplementary Figure 27. **Activity of OES2-0017 in combination with antibiotics that antagonize with polyamines**. Representative checkerboard assays of OES2-0017 and **A)** vancomycin, **B)** kanamycin, and **C)** rifampicin. FICI values and Synergy Finder (SF) scores are reported as mean ± standard deviation of two independent experiments.

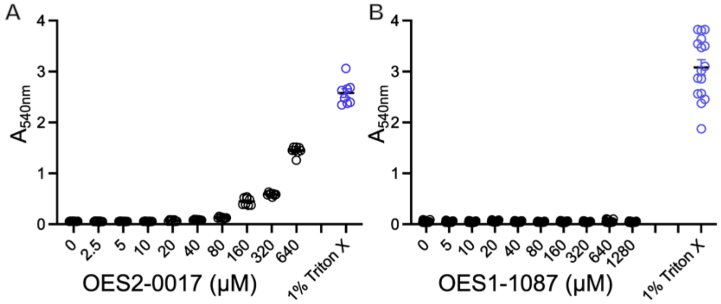


Supplementary Figure 28. **Hemolytic activity of OES2-0017 and OES1-1087.** Effects of **A)** OES2-0017 (n=24) and **B)** OES1-1087 (n=16) on the lysis of sheep red blood cells. Cells begin to lyse at concentrations of OES2-0017 greater than those required to inhibit bacterial growth, and OES1-1087 shows no observable lysis at the tested concentration range. Data is represented as mean ± SEM for three independent experiments.

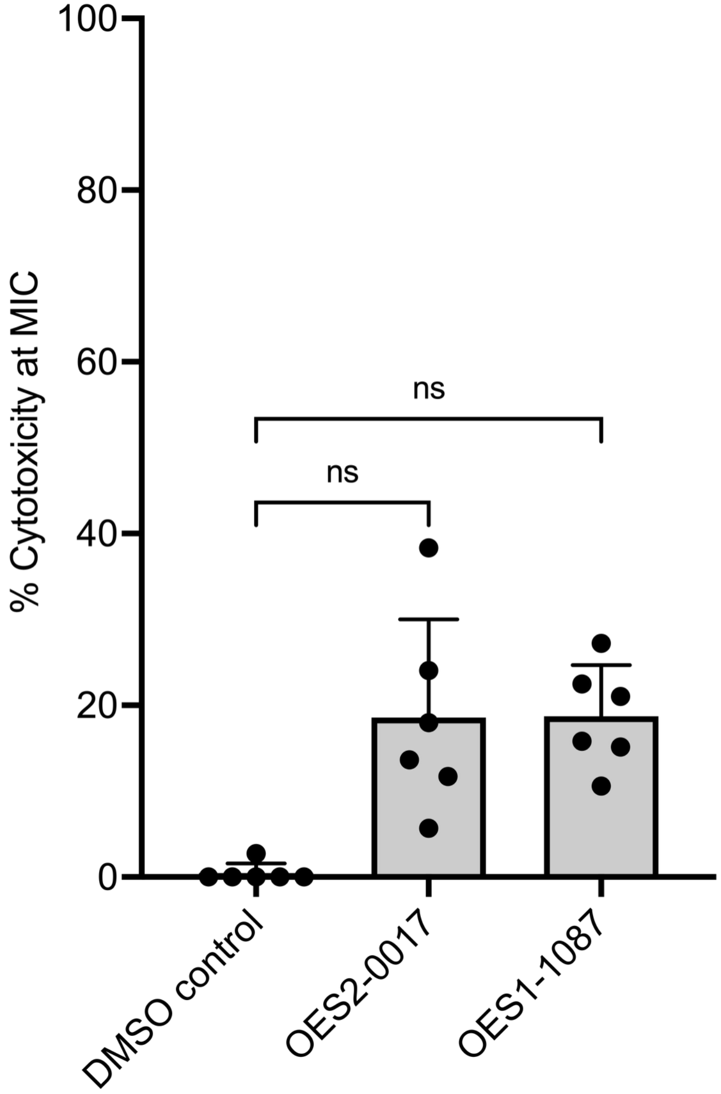


Supplementary Figure 29. **Effects of compound treatment on HepG2 cell damage**. Lactate dehydrogenase leakage from cells was measured after 4 h treatment with compounds at concentrations relative to their respective MICs against *S. aureus* USA300. Data is represented as mean $\pm$ SD for two independent experiments n=6. Statistical significance was determined by ordinary one-way ANOVA using a Dunnett's multiple comparison test using the vehicle control as the comparator *p<0.05, **p<0.01, p<0.001, ****p<0.0001

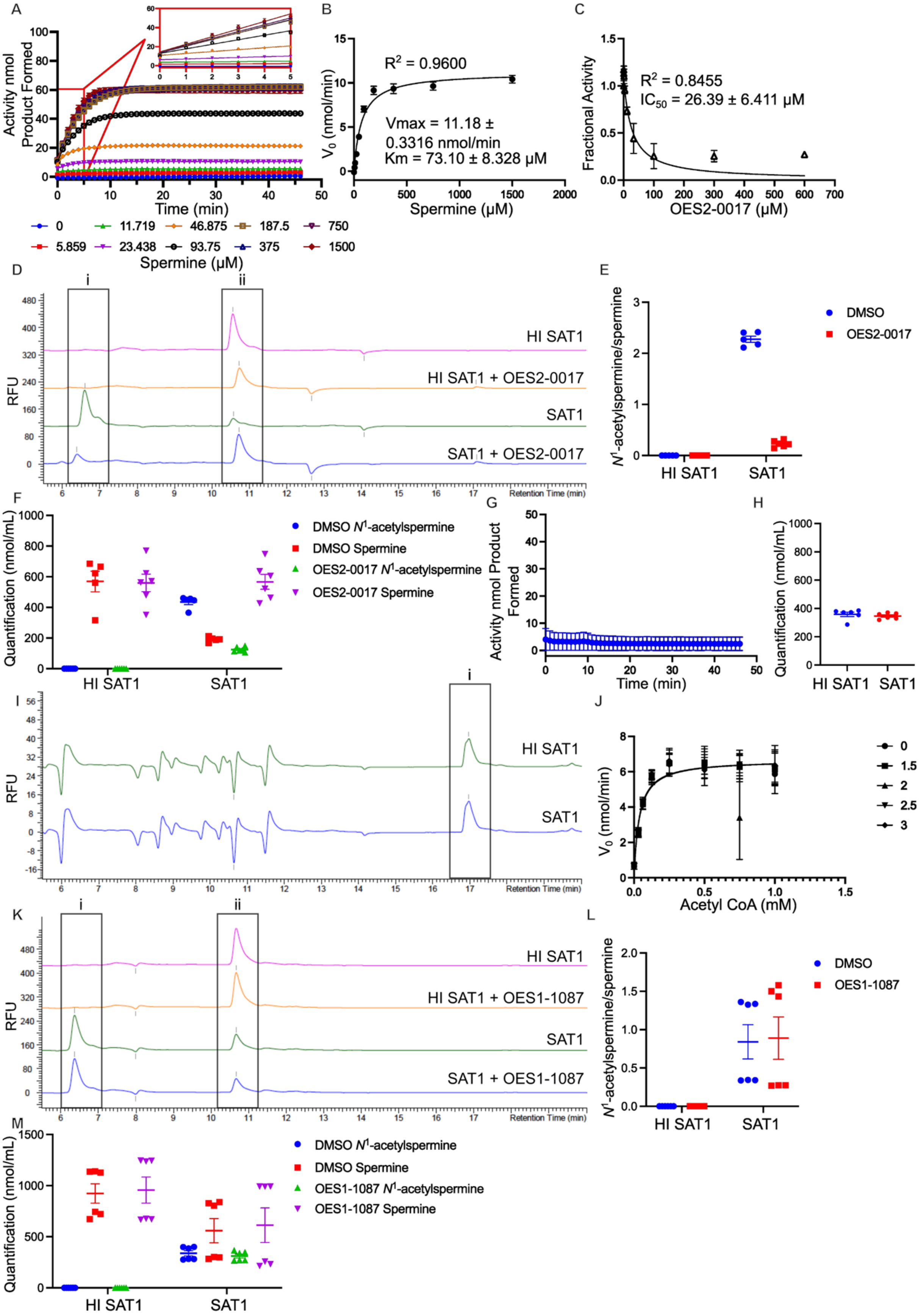
A
B
C
D
E
F
G
H
I
J
K
L
M
Activity nmol Product Formed
Time (min)
0
5.859
11.719
23.438
46.875
93.75
187.5
375
750
1500
Spermine (μM)
R² = 0.9600
Vmax = 11.18 ± 0.3316 nmol/min
Km = 73.10 ± 8.328 μM
V0 (nmol/min)
Spermine (μM)
R² = 0.8455
IC50 = 26.39 ± 6.411 μM
Fractional Activity
OES2-0017 (μM)
i
ii
HI SAT1
HI SAT1 + OES2-0017
SAT1
SAT1 + OES2-0017
RFU
Retention Time (min)
N1-acetylspermine/spermine
DMSO
OES2-0017
Quantification (nmol/mL)
DMSO N1-acetylspermine
DMSO Spermine
OES2-0017 N1-acetylspermine
OES2-0017 Spermine
Acetyl CoA (mM)
0
1.5
2
2.5
3
HI SAT1 + OES1-1087
SAT1 + OES1-1087
OES1-1087
OES1-1087 N1-acetylspermine
OES1-1087 Spermine

Supplementary Figure 30. **Spermine acetyltransferase activity of human SAT1. A)** colorimetric enzymatic SSAT assay of SAT1 with spermine measuring the production of Coenzyme A over time. **B)** Spermine substrate saturation curve generated by calculating initial velocity from the linear portion of **A)**, data was fit with the Michaelis Menten equation in GraphPad Prism 10 with corresponding kinetic parameters displayed on the graph. The acetyltransferase activity of SAT1 was measured under steady-state condition with an excess of AcCoA (0.25 mM) and variable concentrations of spermine. **C)** $IC_{50}$ plot of a colorimetric SSAT enzymatic reaction of SAT1 inhibition by OES2-0017 where SAT1 was pre-incubated with OES2-0017 before adding spermine (75 μM) and Acetyl-CoA (0.25 mM) as substrates. **D)** Representative chromatograms of a 10 min reaction of heat-inactivated SAT1 and SAT1 with and without 300 μM OES2-0017 added to the reaction mixture along with 75 μM spermine and 1 mM Acetyl-CoA to quantify i, $N^1$-acetylspermine and ii, spermine. Primary amines were derivatized with OPA/NAC and detected by fluorescence, represented in relative fluorescence units (ex 340nm/em 450nm). **E)** Ratio of quantified $N^1$-acetylspermine and spermine by HPLC analysis of a 10-min reaction with 300 μM OES2-0017, 75 μM spermine, and 1 mM Acetyl-CoA. **F)** Quantification of $N^1$-acetylspermine and spermine from HPLC analysis of a 10-min reaction with 300 μM OES2-0017, 75 μM spermine, and 1 mM Acetyl-CoA. **G)** Colorimetric SSAT assay of SAT1 with 1 mM Acetyl-CoA and 300 μM OES2-0017 detecting the amount of Coenzyme A liberated during the acetyl-transfer from Acetyl-CoA. **H)** Quantification of OES2-0017 by HPLC analysis of a 10 min reaction of SAT1 and heat inactivated SAT1 with 1 mM Acetyl-CoA and OES2-0017. Primary amines were derivatized with OPA/NAC and detected by fluorescence, represented in relative fluorescence units (ex 340nm/em 450nm). **I)** Representative chromatograms of reactions containing heat-inactivated SAT1 and SAT1 with 1 mM each Acetyl-CoA and OES2-0017, i, OES2-0017. **J)** Substrate saturation curves of SAT1 with a fixed concentration of spermine (75 μM) and variable concentrations of acetyl-CoA and OES1-1087 (mM). **K)** Representative chromatograms of HPLC quantification of *N*1-acetylspermine and spermine by analysis of a 10-minute reaction of SAT1 and heat-inactivated SAT1 with 1 mM spermine and 0.75 mM Acetyl-CoA with or without 3 mM OES1-1087 added to the reaction at the same time as the substrates, i, $N^1$-acetylspermine and ii, spermine. Primary amines were derivatized with OPA/NAC and detected by fluorescence, represented in relative fluorescence units (ex 340nm/em 450nm). **L)** Ratio of quantified $N^1$-acetylspermine and spermine by HPLC analysis of a 10-min reaction with 1 mM spermine and 0.75 mM Acetyl-CoA with or without 3 mM OES1-1087. **M)** Quantification of $N^1$-acetylspermine and spermine from HPLC analysis of a 10-min reaction 1 mM spermine and 0.75 mM Acetyl-CoA with or without 3 mM OES1-1087. Data is represented as mean ± SEM from three independent experiments **C)** n=3 and two independent experiments **A,** and **B,** n= 4, **E, F, H, L,** and **M)** n=6, **G** and **J)** n=2.

**Spermine**

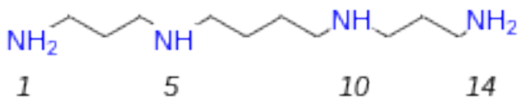


**OES2-0045**

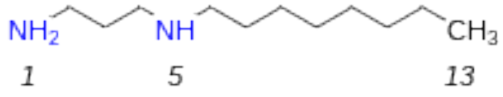


**OES2-0017**

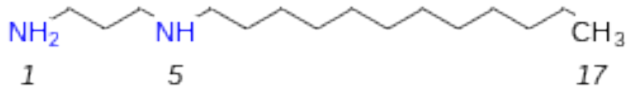


**OES2-0085**

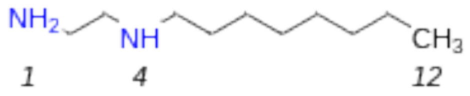


**OES2-0086**

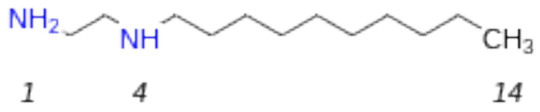


**OES2-0047**

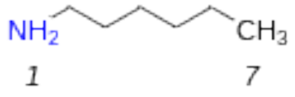


**OES2-0046**

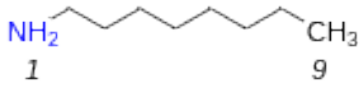


**OES2-0077**

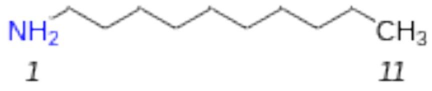


**OES2-0052**

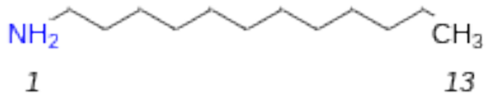


Supplementary Figure 31. 2D chemical structure of OES2 series compounds and spermine.

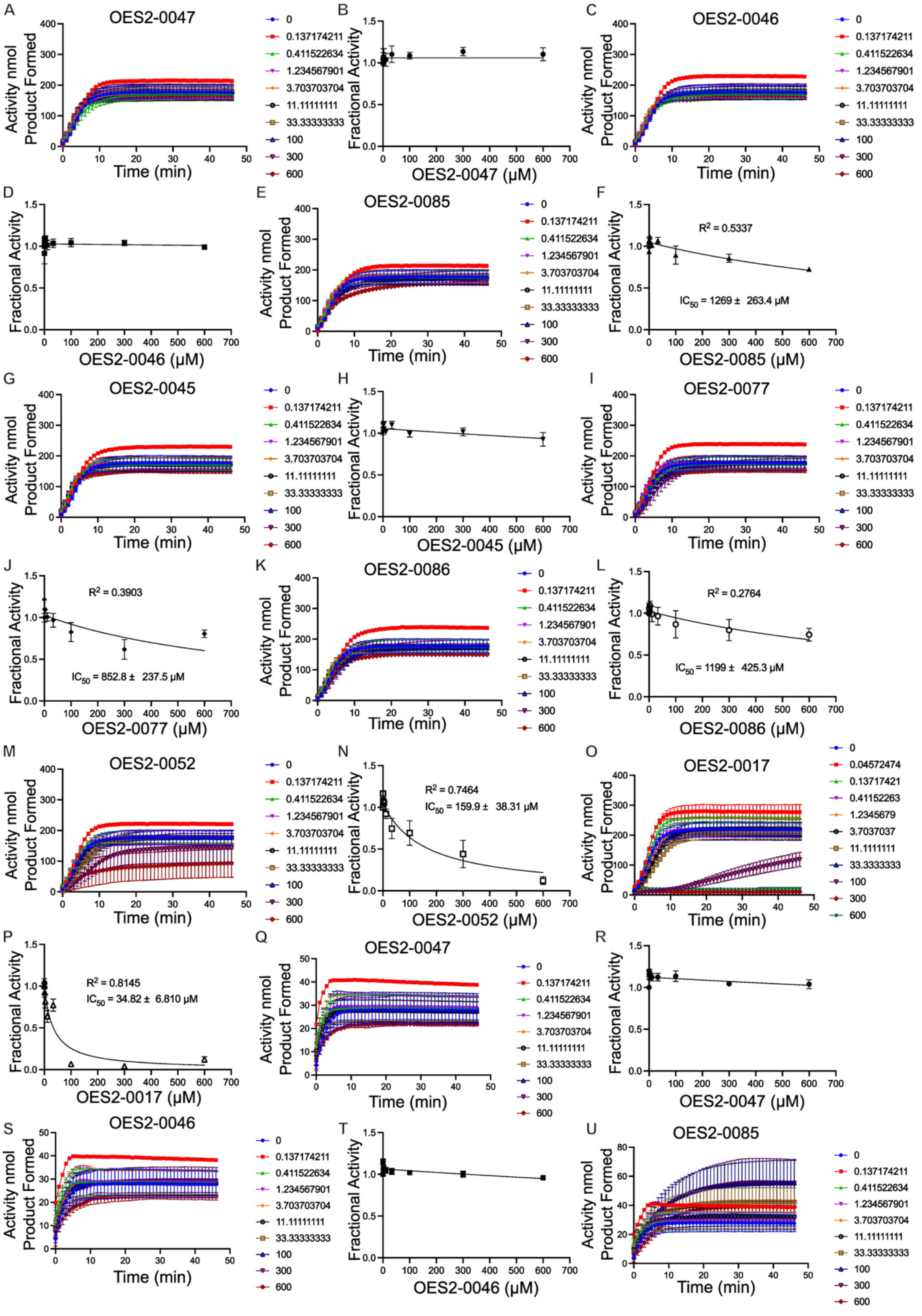
A
OES2-0047
Activity nmol Product Formed
Time (min)
0
0.137174211
0.411522634
1.234567901
3.703703704
11.11111111
33.33333333
100
300
600
B
Fractional Activity
OES2-0047 (μM)
C
OES2-0046
Activity nmol Product Formed
Time (min)
D
Fractional Activity
OES2-0046 (μM)
E
OES2-0085
Activity nmol Product Formed
Time (min)
F
Fractional Activity
R² = 0.5337
IC50 = 1269 ± 263.4 μM
OES2-0085 (μM)
G
OES2-0045
Activity nmol Product Formed
Time (min)
H
Fractional Activity
OES2-0045 (μM)
I
OES2-0077
Activity nmol Product Formed
Time (min)
J
Fractional Activity
R² = 0.3903
IC50 = 852.8 ± 237.5 μM
OES2-0077 (μM)
K
OES2-0086
Activity nmol Product Formed
Time (min)
L
Fractional Activity
R² = 0.2764
IC50 = 1199 ± 425.3 μM
OES2-0086 (μM)
M
OES2-0052
Activity nmol Product Formed
Time (min)
N
Fractional Activity
R² = 0.7464
IC50 = 159.9 ± 38.31 μM
OES2-0052 (μM)
O
OES2-0017
Activity nmol Product Formed
Time (min)
0
0.04572474
0.13717421
0.41152263
1.2345679
3.7037037
11.1111111
33.3333333
100
300
600
P
Fractional Activity
R² = 0.8145
IC50 = 34.82 ± 6.810 μM
OES2-0017 (μM)
Q
OES2-0047
Activity nmol Product Formed
Time (min)
R
Fractional Activity
OES2-0047 (μM)
S
OES2-0046
Activity nmol Product Formed
Time (min)
T
Fractional Activity
OES2-0046 (μM)
U
OES2-0085
Activity nmol Product Formed
Time (min)

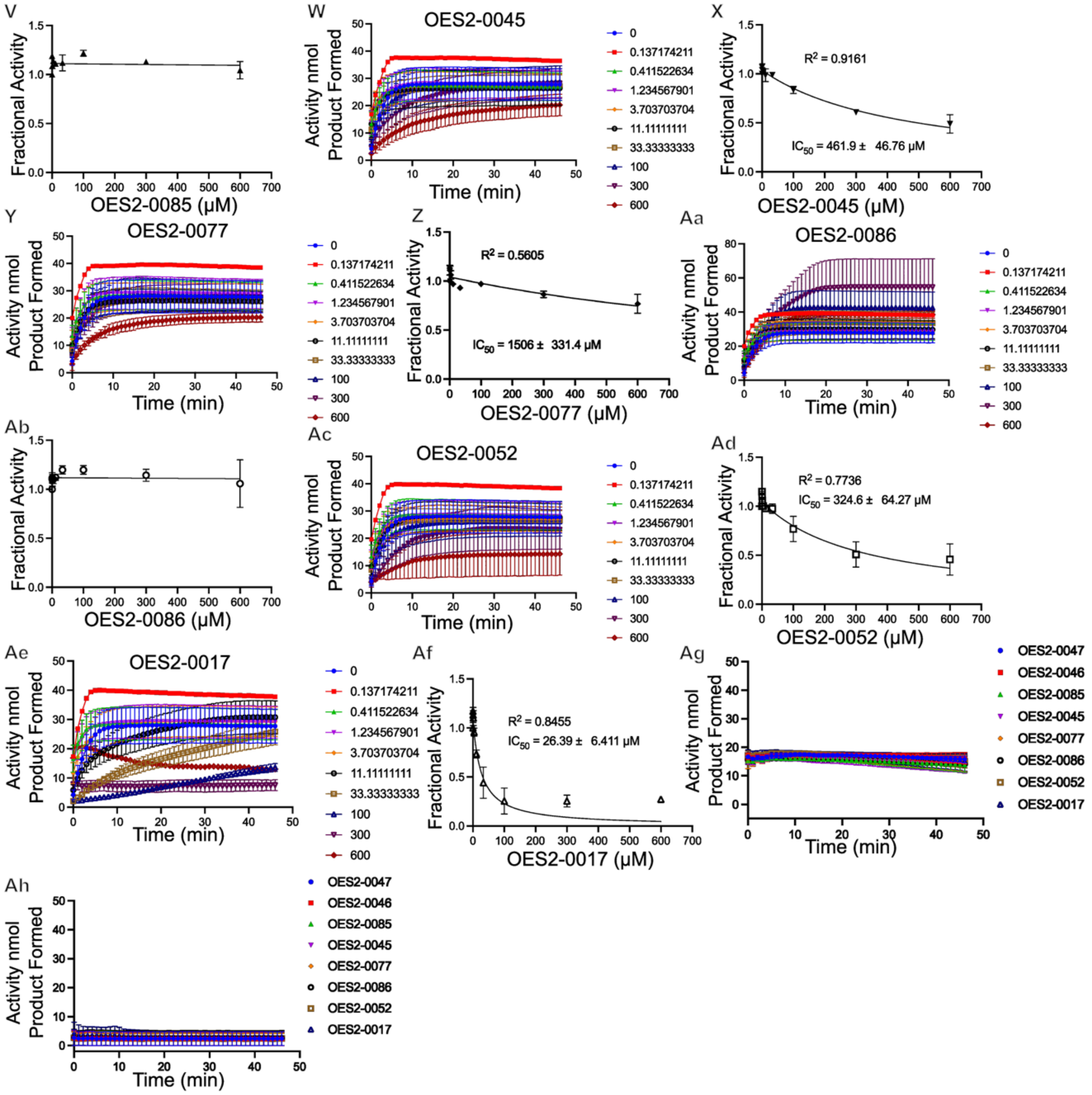


Supplementary Figure 32. **Inhibitory effects of OES2-0017 analogs on SpeG and SAT1.** Colorimetric enzymatic assays representing **A-P)** inhibition of SpeG by **A** and **B)** OES2-0047, **C** and **D)** OES2-0046, **E** and **F)** OES2-0085, **G** and **H)** OES2-0045, **I** and **J)** OES2-0077, **K** and **L)** OES2-0086, **M** and **N)** OES2-0052, and **O** and **P)** OES2-0017. **Q-Af)** Inhibition of SAT1 by **Q** and **R)** OES2-0047, **S** and **T)** OES2-0046, **U** and **V)** OES2-0085, **W** and **X)** OES2-0045, **Y** and **Z)** OES2-0077, **Aa** and **Ab)** OES2-0086, **Ac** and **Ad)** OES2-0052, and **Ae** and **Af)** OES2-0017. **Ag-Ah)** Colorimetric enzymatic assays containing 300 μM of each analog and 1 mM or 0.25 mM AcCoA to assess if analogs can be acetylated by **Ag)** SpeG and **Ah)** SAT1, respectively. Data are represented as the mean ± SEM from four independent experiments **A-N**) n=4, seven independent experiments **O** and **P**, n=7, three independent experiments **Q-Af)** n=3, and two independent experiments **Ag)** and **Ah)**, n=2.

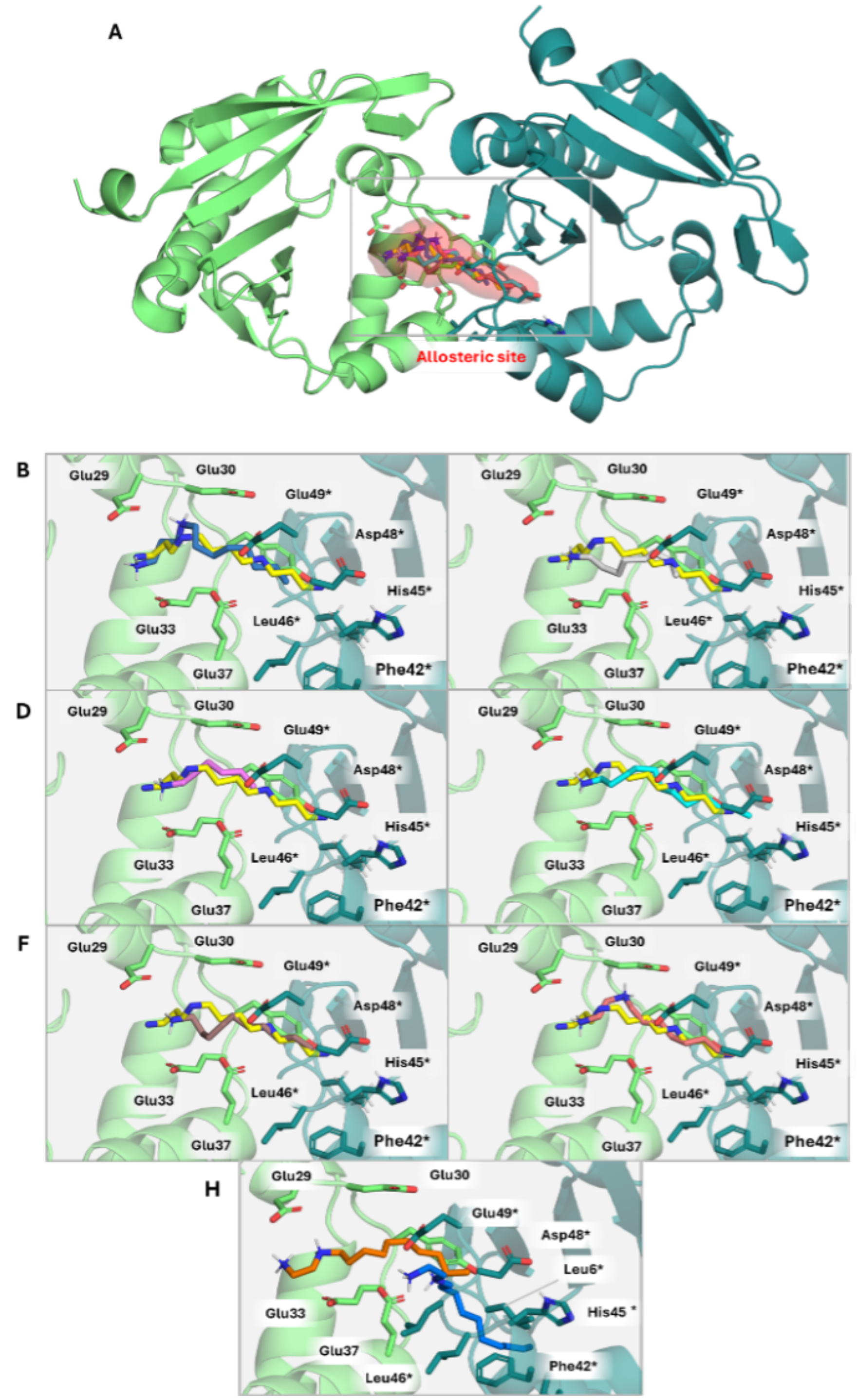


Supplementary Figure 33. **Selected binding poses obtained from docking calculations of OES2 compounds into SpeG dimer. A)** General view of the SpeG dimer (PDB ID 8fv1). The allosteric site is represented with a red surface and selected poses are superimposed. **B-G)** Detail of best docked pose and main interactions into the spermine binding site in SpeG dimer of compounds OES2-0045 (**B**, blue), OES2-0046 (**C**, light grey) and OES2-0047 (**D**, pink), OES2-0052 (**E**, cyan), OES2-0077 (**F**, brown), OES2-0085 (**G**, orange), and OES2-0086 (**H**, orange). The crystallographic pose of spermine is shown in yellow as reference. Monomers A and B are depicted in green and aquamarine blue, respectively.

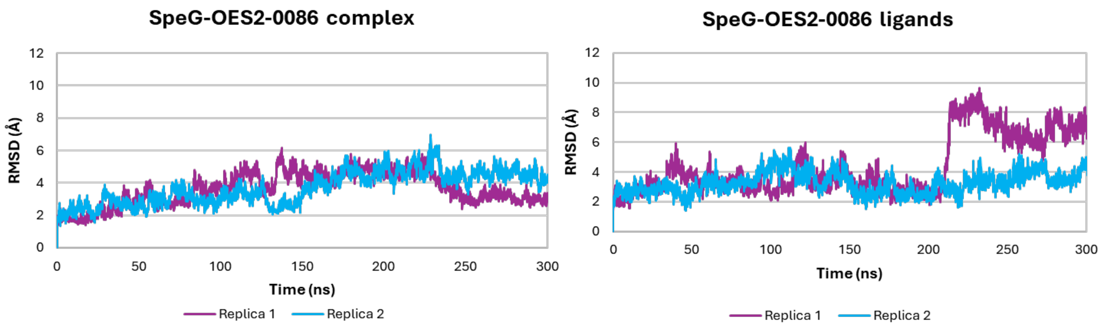


Supplementary Figure 34. Root Mean Square Deviation (RMSD) of the SpeG dimer (PDB ID 8fv1) in complex with OES2-0086. RMSD has been calculated for the whole complex (left) and only for the ligands (right).

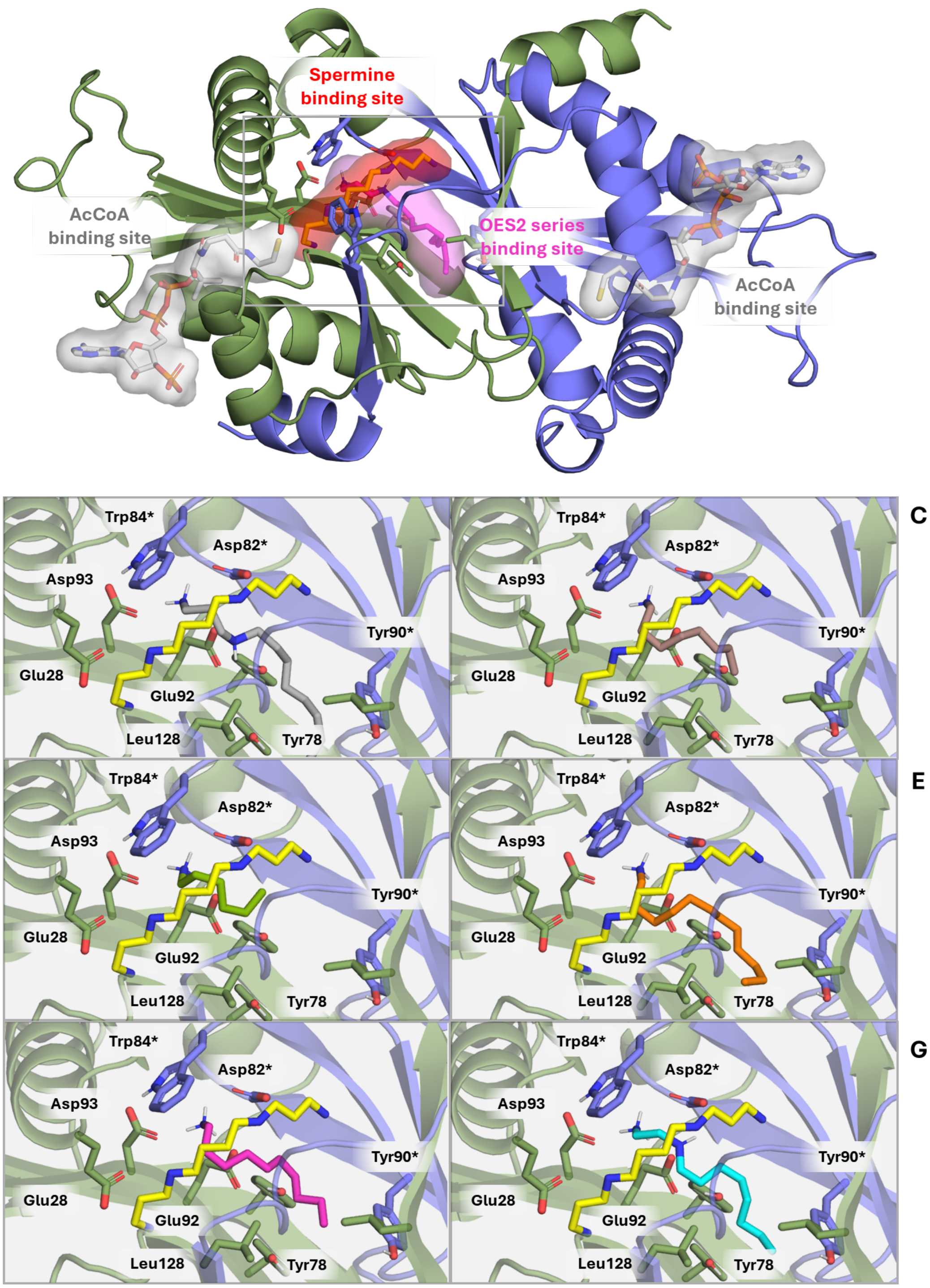


Supplementary Figure 35. Selected binding poses obtained from docking calculations of OES2 series compounds into human SAT1 dimer (PDB ID 2b58). **A)** General view of the SAT1 dimer. The spermine binding area is represented with a red surface and that of the OES2 series is represented as a pale pink surface. **B-G)** Detail of the interactions of the crystallographic structure of spermine (yellow) in comparison with OES2-0045 (**B**, light grey), OES2-0046 (**C**, brown) and OES2-0047 (**D**, green), OES2-0052 (**E**, orange), OES2-0077 (**F**, pink) and OES2-0085 (**G**, cyan). Monomers A and B are depicted in green and blue, respectively.

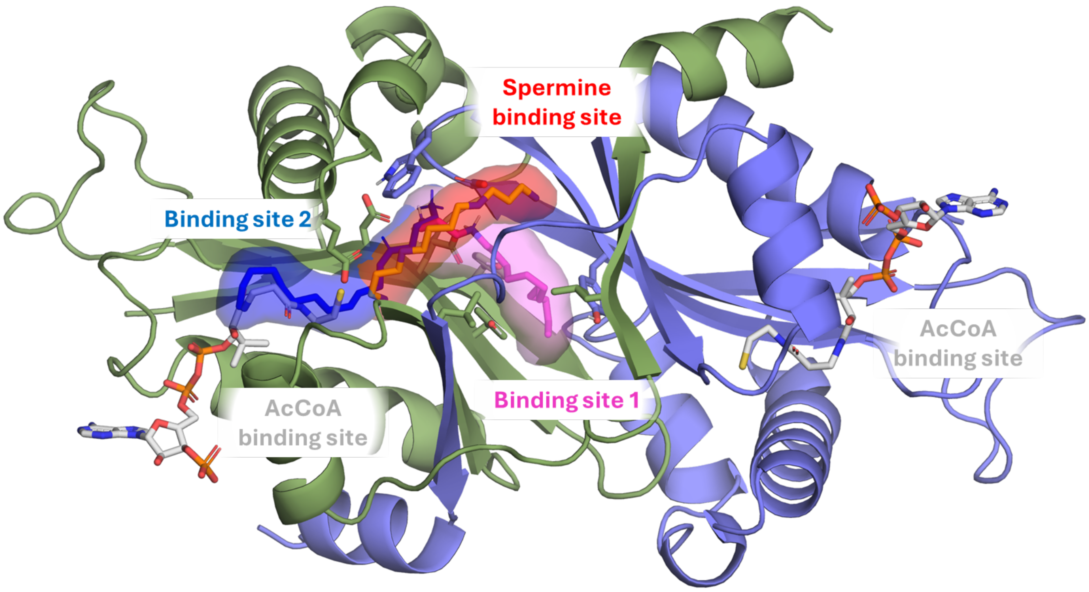


Supplementary Figure 36. Description of the binding sites in the human SAT1 dimer (PDB ID 2b58). Spermine binding site (red surface) shares the acidic region with binding sites 1 and 2 (pink and blue, respectively. AcCoA is depicted as white sticks, monomer A in green and monomer B in blue.

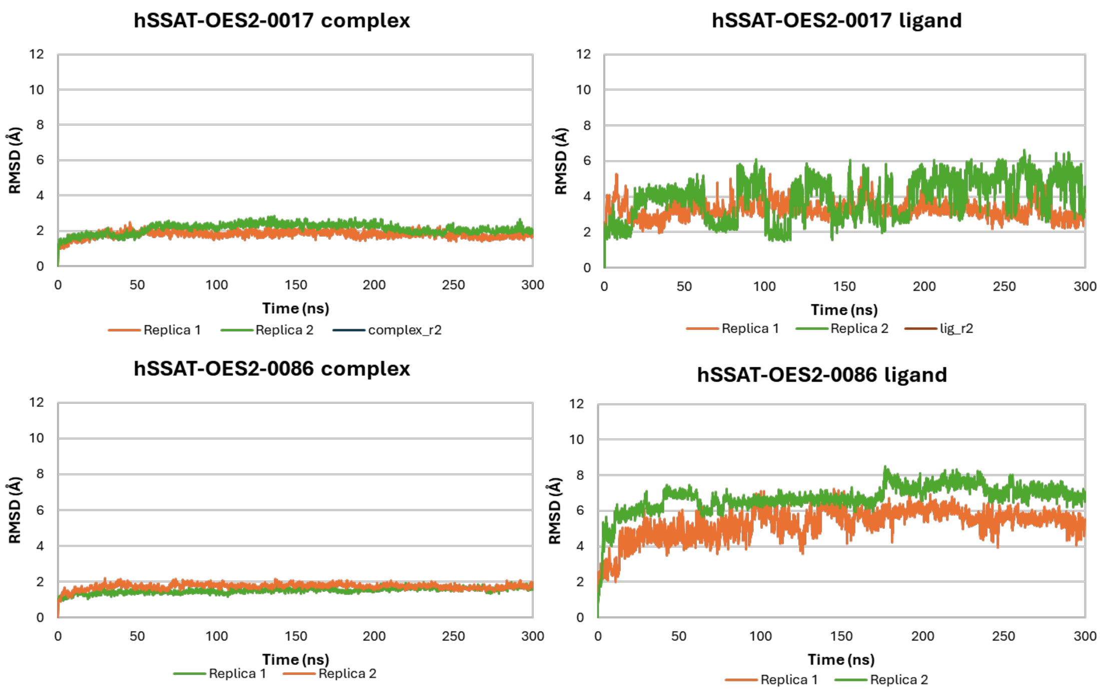


Supplementary Figure 37. Root Mean Square Deviation (RMSD) of the human SAT1 (hSSAT) dimer (PDB ID 2b58) in complex with OES2-0017 (top) and OES2-0086 (bottom) in binding site 1. RMSD has been calculated for the whole complex (left) and only for the ligands (right).

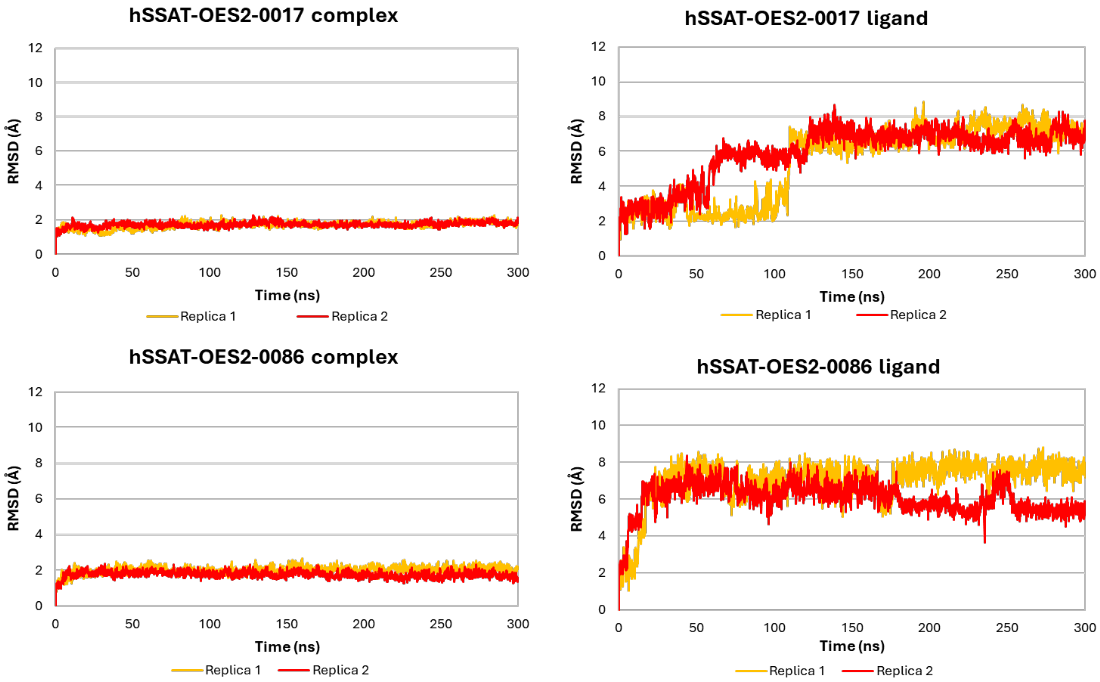


Supplementary Figure 38. Root Mean Square Deviation (RMSD) of the human SAT1 dimer (PDB ID 2b58) in complex with OES2-0017 (top) and OES2-0086 (bottom) in binding site 2. RMSD has been calculated for the whole complex (left) and only for the ligands (right).

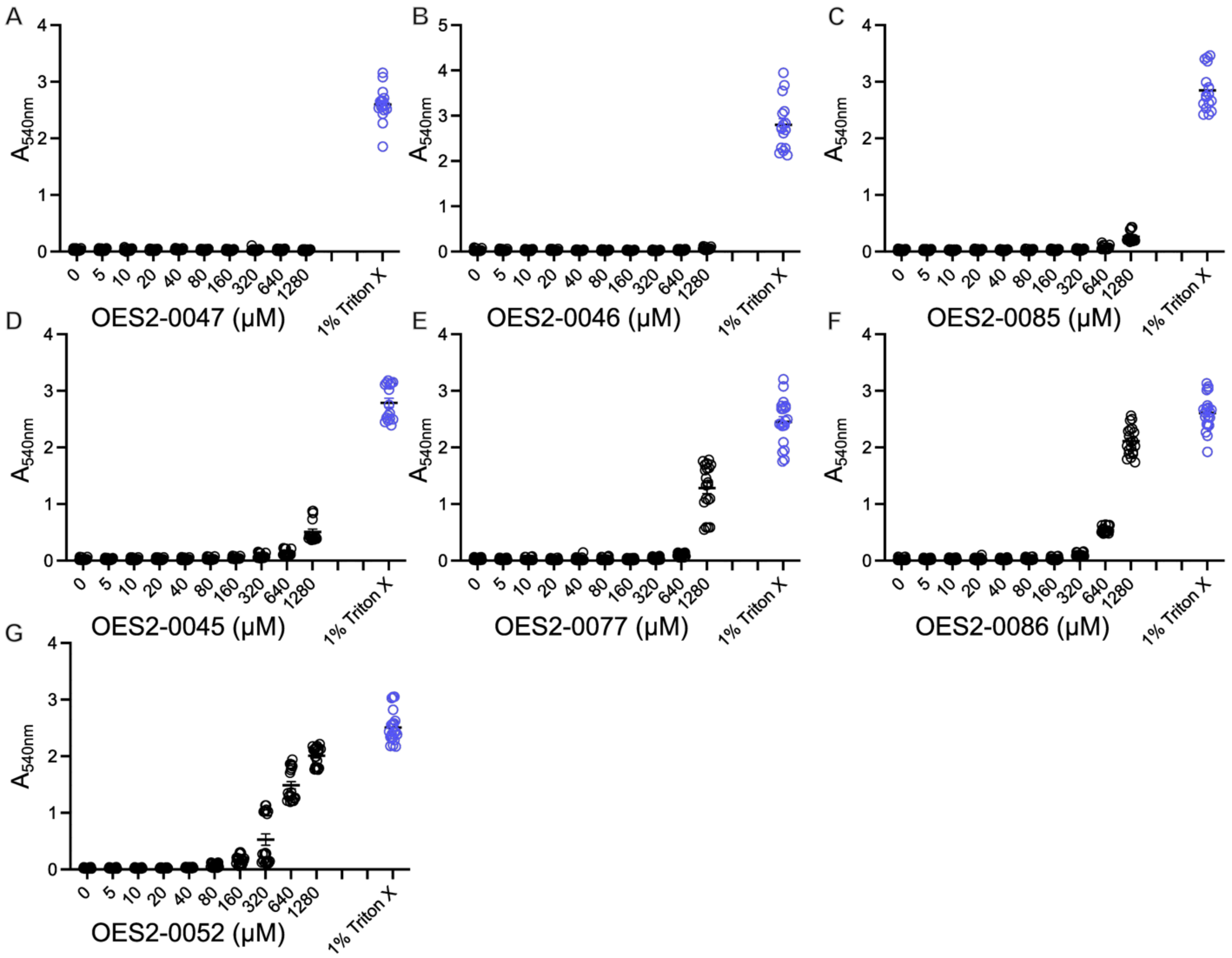


Supplementary Figure 39. **Hemolytic effects of OES2-0017 analogs on sheep red blood cells.** Hemolytic effects of **A)** OES2-0047, **B)** OES2-0046, **C)** OES2-0085, **D)** OES2-0045, **E)** OES2-0077, **F)** OES2-0086, and **G)** OES2-0052. Data are represented as the mean ± SEM from three independent experiments **A-D)** n=16 and **E-F)** n=20.

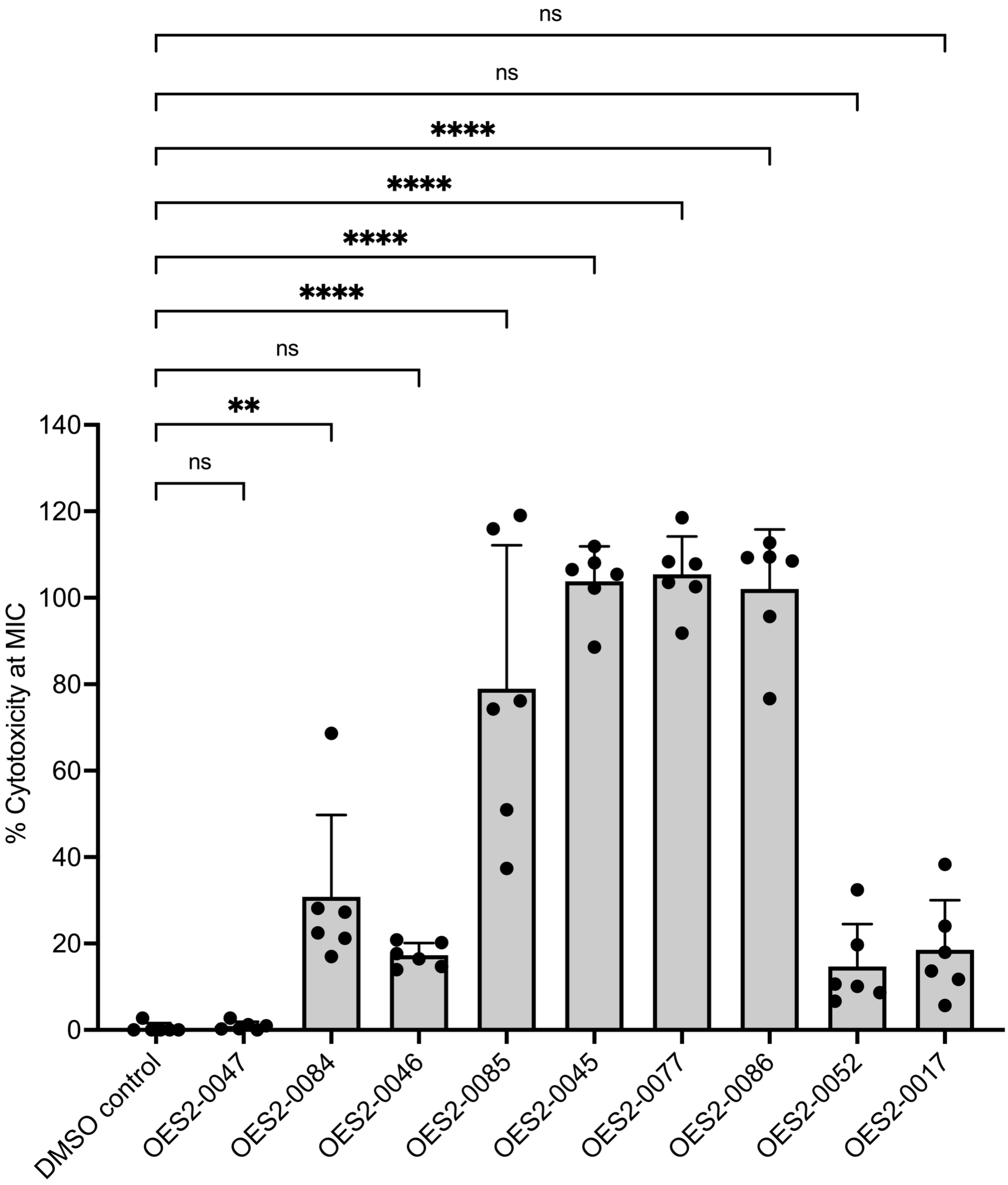


Supplementary Figure 40. **Effects of compound treatment on HepG2 cell damage**. Lactate dehydrogenase leakage from cells was measured after 4 h treatment with compounds at concentrations equivalent to their respective MICs against *S. aureus* USA300. Data is represented as mean ± SD for two independent experiments n=6. Statistical significance was determined by ordinary one-way ANOVA using a Dunnett's multiple comparison test using the vehicle control as the comparator *p<0.05, **p<0.01, p<0.001, ****p<0.0001.

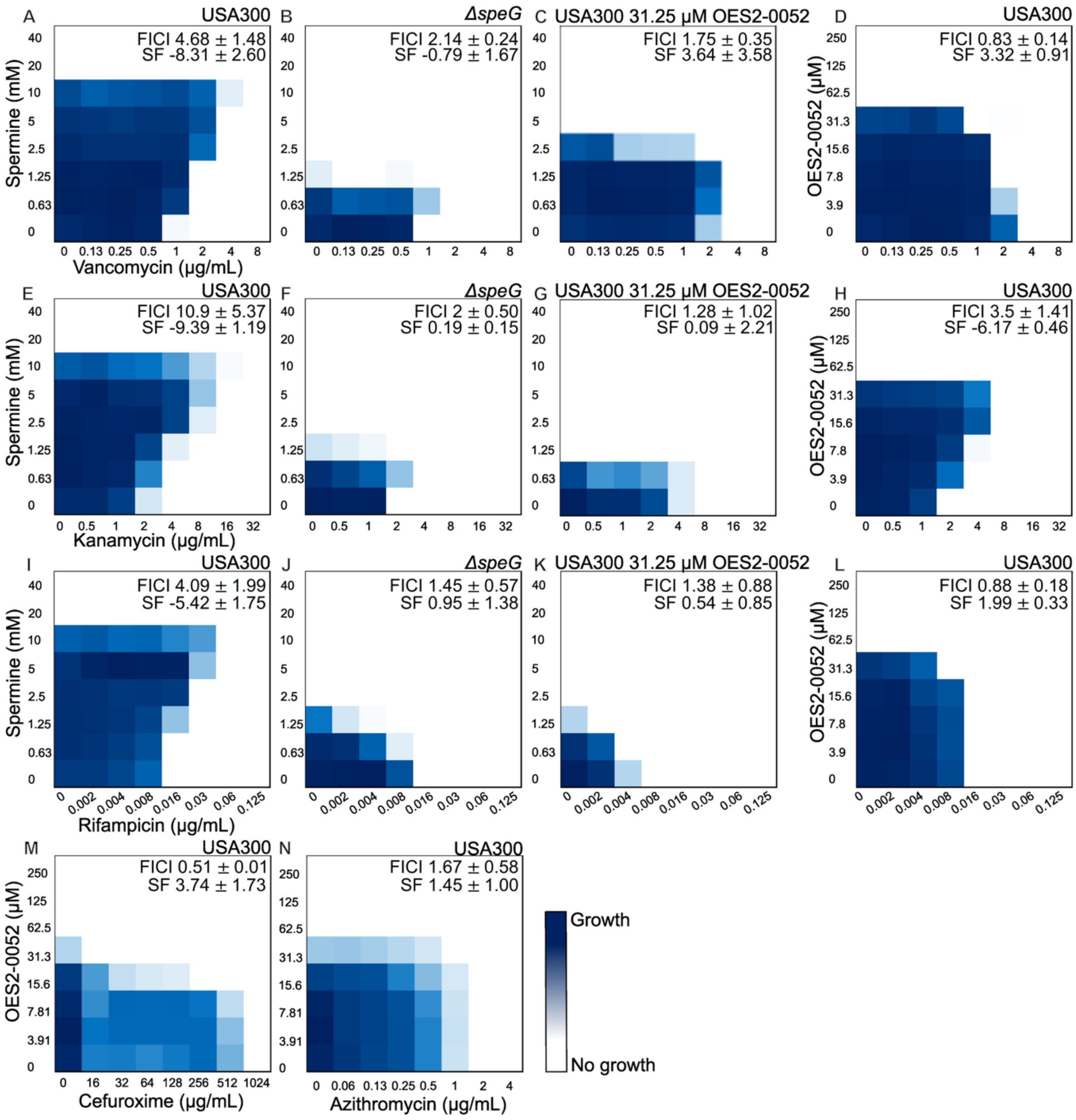


Supplementary Figure 41. **The potential of OES2-0052 as an antibiotic adjuvant.** Representative checkerboard assays of spermine and vancomycin **(A-C)**, OES2-0052 and vancomycin **(D)**, spermine and kanamycin **(E-H)**, OES2-0052 and kanamycin **(I)**, spermine and rifampicin **(J-L)**, and OES2-0052 and rifampicin against *S. aureus* **A, D, E, H, I, and L)** USA300, **B, F, J)** Δ*speG*, and **C, G, K)** USA300 supplemented with a sub-inhibitory concentration of OES2-0052 where OES2-0052 abolishes the protective effect of polyamines from the antibiotics. Representative checkerboard assay of OES2-0052 and cefuroxime **(M)** and azithromycin **(N)** against *S. aureus* USA300, where OES2-0052 potentiates the antibiotic activity. FICI values and Synergy Finder (SF) scores are represented as mean ± standard deviation n=10 **(A)**, n=7 **(B)**, n=2 **(C, G, H, K, and L)**, n=8 **(I),** n=5 **(E, F, and J)**, and n=3 **(D, M and N).**

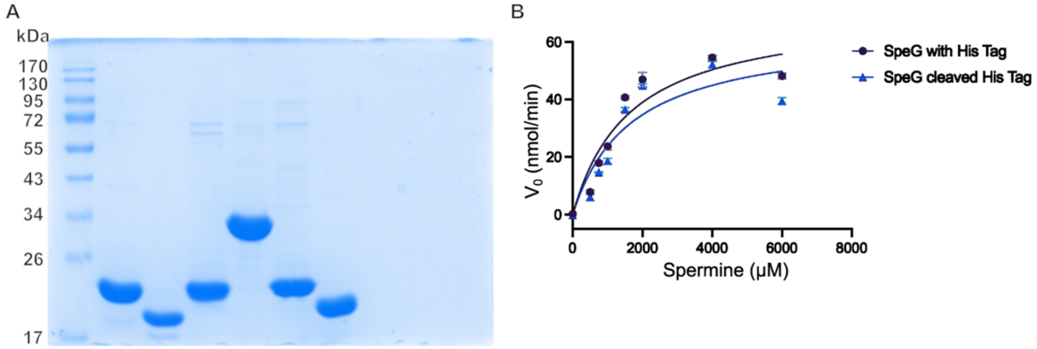


Supplementary Figure 42. **Purified proteins used in this study. A)** 12% SDS-PAGE gel of proteins used in this study Lane 1 EZ-run protein ladder, Lane 2 SpeG, Lane 3 SpeG cleaved His tag, Lane 4 SAT1, Lane 5 SAUSA300_2083, Lane 6 PaiA$_{Sa}$, Lane 7 SAUSA300_0441. **B)** Colorimetric enzymatic assay with SpeG and SpeG with cleaved His tag shows no appreciable difference in spermine acetylation. Data is represented as mean ± SEM from one experiment n=2.
